# A platform of robust patient-derived leukemia models covering subgroups for which no cell lines exist

**DOI:** 10.1101/2025.09.26.677299

**Authors:** Binje Vick, Vindi Jurinovic, Kristina Kuhbandner, Lena Lagally, Lisa Latzko, Chiara Arnreich, Gerulf Hänel, Amelie Muth, Maja Rothenberg-Thurley, Annika M. Dufour, Stephanie Schneider, Lesca M. Holdt, Liliana Mura, Fabian Klein, Annette Frank, Maya C. André, Claudia D. Baldus, Martin Carroll, Christine Dierks, Martin Ebinger, Katharina S. Götze, Pablo Menéndez, Christian Récher, Ambrine Sahal, Jean-Emmanuel Sarry, Christian Thiede, Talía Velasco-Hernández, Xiaoyan Wei, Jan Henning Klusmann, Michael von Bergwelt-Baildon, Wolfgang Hiddemann, Klaus H. Metzeler, Philipp A. Greif, Marion Subklewe, Sebastian Vosberg, Tobias Herold, Karsten Spiekermann, Irmela Jeremias

## Abstract

Preclinical cancer research requires robust model systems, especially for poor prognosis entities like acute myeloid leukemia (AML), a highly aggressive blood cancer. Here, primary tumor cells from 137 AML patients of all age groups were transplanted into immune compromised mice to generate patient-derived xenografts (PDX). From these, 23 models enable robust, virtually endless serial re-transplantation and are amenable to lentiviral genetic engineering (^*^PDX AML models). These models primarily originate from patients with highly aggressive, relapsed disease. Comprehensive genomic, transcriptomic, and epigenomic analyses confirmed that they replicate primary AML biology more faithfully than conventional cell lines. Notably, ^*^PDX AML models include AML subgroups that are underrepresented or absent in existing model systems, such as cytogenetically normal or *IDH1/2*-mutant AML. They withstand freeze-thaw cycles, making them suitable for broad distribution and reproducibility across research institutions. Luciferase-based *in vivo* imaging enables real-time monitoring of tumor progression and treatment responses in preclinical trials. Surprisingly, long-term treatment, including repeated cytarabine therapy over a period of one year, showed a gradual reduction in leukemia cell proliferation, which decreased continuously after each treatment block. Collectively, our ^*^PDX models represent a robust, versatile, and relevant platform that holds great promise to accelerate translational research for the benefit of cancer patients.

**Visual Abstract:** 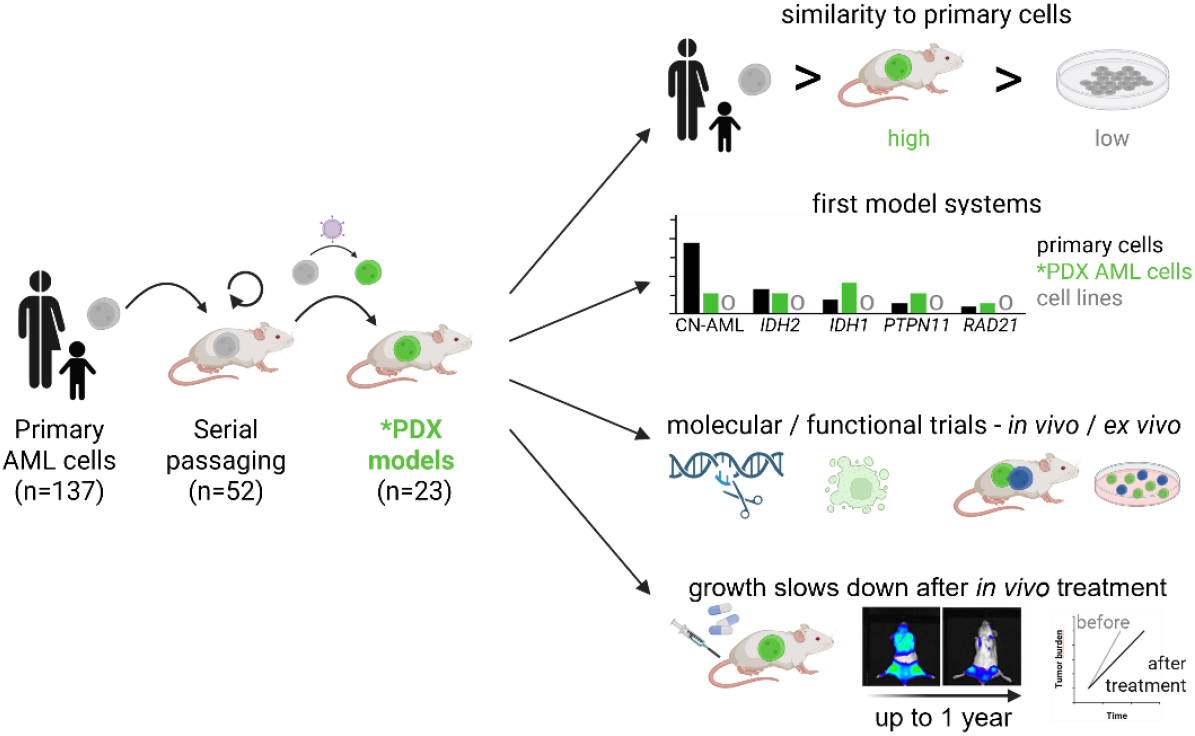

**Key Points:**

- We present new robust AML PDX models covering subgroups for which no cell lines exist for use in various *ex vivo* and *in vivo* applications.
- ^*^PDX models enable serial transplantation, genetic engineering and better representation of primary AML biology than cell lines.
- One-year *in vivo* trials mimicking clinical chemotherapy showed surprising gradual decline in leukemia growth after each treatment block.

## Introduction

Acute myeloid leukemia (AML) is an aggressive and heterogenous hematologic malignancy characterized by the clonal expansion of immature myeloid cells^1^. Initial chemotherapy can induce complete remission, but high relapse rates lead to a persistently unfavorable prognosis in AML, highlighting an urgent need for better treatment strategies^2^. A major challenge to develop novel therapies for treatment of cancer is the lack of convenient and faithful model systems. Primary patient samples are rare and often low in cell numbers, especially in children suffering from hematologic malignancies, and cells don’t consistently grow *ex vivo*^3,4^. Cell lines are convenient to culture, but poorly model important features of primary tumor cells as they are heavily biased towards unstable karyotypes and certain genetic abnormalities and show altered drug response^5-7^. While establishing a cell line from a primary tumor sample is often successful in solid tumors, this is rarely the case in liquid tumors and often accompanied by acquired confounding features^8^. Recent 3D organoid models of the bone marrow (BM) niche with multiple cell types^9^ lack, for example, mechanical forces by blood flow and cannot model tumor cell migration into the BM.

Patient-derived xenograft (PDX) models are highly attractive tools for cancer research. By transplanting primary human cancer cells into immunodeficient mice, they proliferate in an *in vivo* microenvironment. Orthotopically growing tumors as in AML considerably facilitate our understanding of disease pathogenesis, support reliable drug testing and more accurately reflect the genetic and biological heterogeneity of the human disease^10-12^. PDXs have offered critical insights into cancer stem cell and AML biology, including mechanisms driving disease initiation, progression and therapy resistance, (epi)genetic clonal evolution, the biology of leukemia-initiating cells and interaction with the BM niche^13-16^ (and many more).

Most PDX AML studies focus on primary engraftment of patient cells. As limitation, primary cells are only available in limited cell numbers in many countries, restricting their availability and use in repetitive assays. A decade ago, we published our first serially transplantable PDX AML models, pioneered genetic engineering of PDX AML cells and shared the models across multiple laboratories worldwide^17^. Building on these initial results, we have since increased the number of available models, characterized the models in depth and were able to generate PDX models for AML subgroups for which no cell lines exist. These <u>s</u>erially transplantable <u>a</u>nd robust (STAR), genetically engineered PDX (^*^PDX) AML models represent an attractive tool for future research in the field.

## Results

### Primary AML engraftment in NSG mice is favored by aggressive disease and female mice

Primary samples of 102 adult AML patients were transplanted into mice over a period of 12 years (Table S1); of these, 74 samples were collected at the University Hospital at LMU Munich (cohort 1, Table S2), and 28 samples were collected either at hospitals in Germany in the framework of the German Cancer Consortium (n=23) or at the hospital of the University of Pennsylvania (n=5) (together cohort 2). From 22 patients, PDX models were established by collaboration partners, among them five from pediatric patients (cohort 3). In an independent approach, 13 primary AML samples from children, mainly with *KMT2A*-rearranged AML, were transplanted at Hannover Medical School or University Hospital Halle (cohort 4). In total, cells from 137 patients were included into the study (Figure 1A; Table S1). Using non-irradiated NSG mice and intra-venous injection, between 40% and 60% of samples across all cohorts engrafted, in line with published data (Figure 1A)^18,19^.

**Figure 1.**
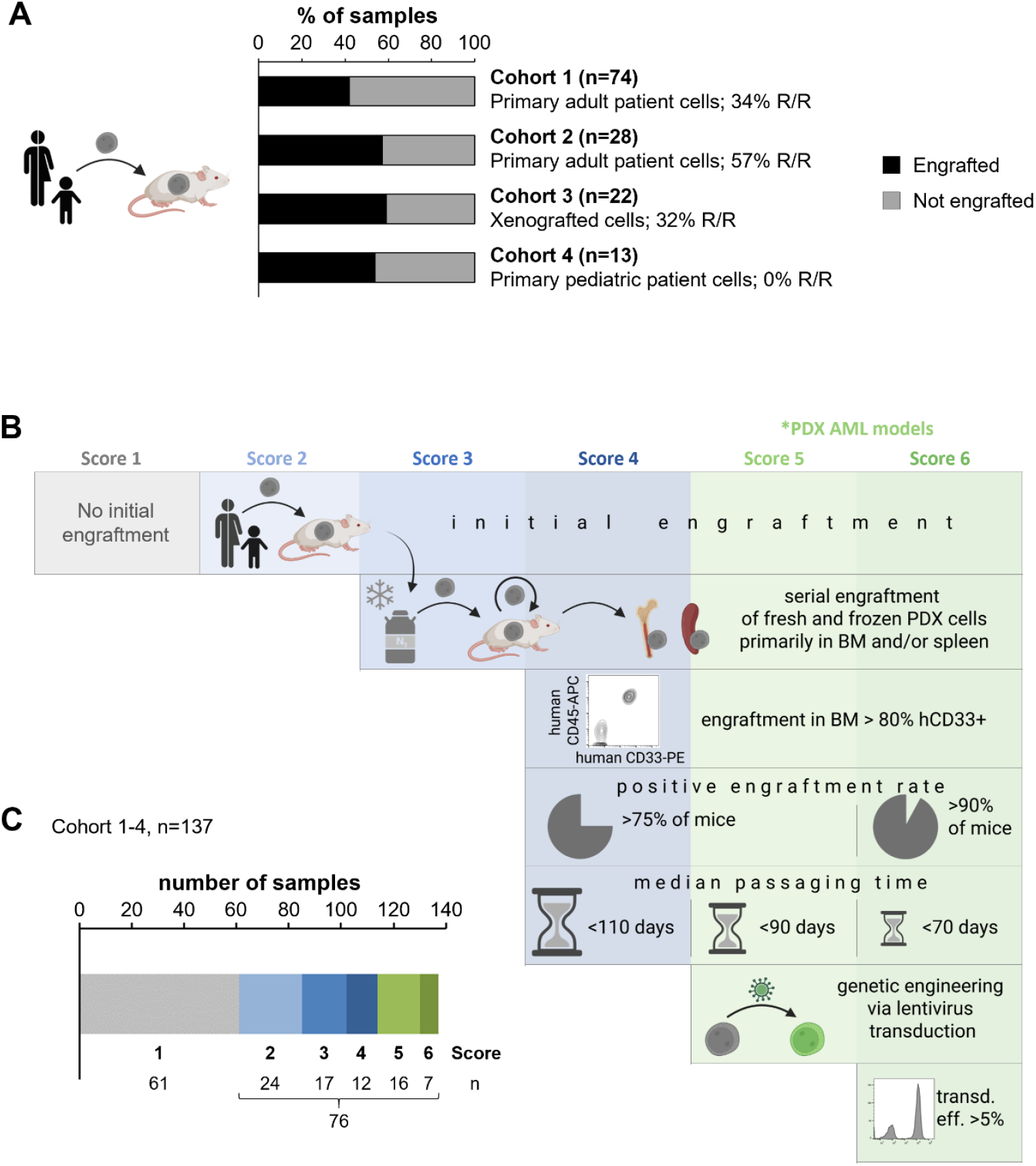
Both primary engraftment and robust serial engraftment are confined to a minority of PDX models. **A** Primary samples of adult AML patients (cohort 1, n=74 and cohort 2, n=28), xenografted models of adult or pediatric AML patients (cohort 3, n=22) or primary samples of pediatric AML patients (cohort 4, n=13) were transplanted into one to eight NSG mice via tail vein injection (2.5E5 – 2E7 cells per mouse) and positive engraftment was analyzed. If at least one mouse showed positive engraftment, the sample was classified as engrafted. R/R: relapsed/primary refractory. **B**,**C** Primary AML samples or AML PDX models were serially transplanted into NSG mice, and positive engraftment was analyzed. **B** Scheme illustrating the scoring system based on specific features. **C** Distribution of samples within Score 2 – Score 6.

To assess predictive parameters for primary engraftment, we analyzed concomitant clinical data available for cohort 1 and reproduced known prediction factors such as relapse/primary refractory (R/R) disease and presence of *FLT3*-ITD mutations (Table S2; Figure S1A)^18,20-24^. Additionally, *BCOR*-mutated samples, although low in numbers, showed a trend towards higher engraftment capacity (Figure S1A). Combining the effect of these two mutations by creating a composite binary variable did not improve the prediction model (not shown). Other examined variables such as patient age, gender, white blood cell count, or French-American-British (FAB) group did not predict engraftment (Table S1 and data not shown). In accordance with literature, xenograft potential negatively correlated with overall survival (OS) probability, with a major difference in 1-year OS between patients whose cells did or did not engraft (Figure S1B)^20,21^. This is consistent with the extremely poor prognosis for patients who suffer relapse, e.g., after allogeneic BM transplant, when treatment may focus on supportive care.

Conflicting results have been published concerning engraftment capacity in male versus female NSG mice^25,26^. In our study, primary cells from male patients transplanted into male mice showed the lowest engraftment rates, while primary cells from female patients transplanted into female mice showed the highest engraftment rates; age of recipient mice, however, did not affect engraftment efficiency (Figure S1C,D).

### A rare subfraction of primografts allows generation of PDX models with robust serial engraftment and genetic engineering

Serial re-transplantation of PDX AML cells beyond two passages has been scarcely studied, except for our own previous data^15,17,24,27,28^. Here, we aimed to establish robust, serially transplantable and fast engrafting PDX AML models which allow freeze/thaw cycles and genetic engineering as new tools for complex downstream applications *ex vivo* and *in vivo*. To identify appropriate PDX models, we defined subjective characteristics based on prior experience to categorize PDX AML models into six groups, considering the following features: (i) positive primary engraftment, (ii) serial engraftment capacity of frozen/thawed cells, (iii) high engraftment rate in the BM and in a majority of mice, (iv) passaging time (period from transplantation to end-stage leukemia) below 16 weeks, and (v) ability for lentiviral transduction (Figure 1B). Of the 137 AML samples included in the study, 76 (55%) showed positive primary engraftment (score ≥2), but only 52 (38%) showed positive serial engraftment (score ≥3). This contrasts with the closely related disease acute lymphoblastic leukemia (ALL), where both primary engraftment and serial passaging is feasible for virtually all samples^29^. Robust and fast serial engraftment was achieved in 35 samples (25.5%, score ≥4). While genetic manipulation was not conducted in samples of cohort 4, 23/124 (18.5%) samples of cohort 1-3 allowed genetic engineering (score ≥5, Figure 1B,C and S2A). We called those score 5/6 models ^*^PDX AML models to indicate their ability for <u>s</u>erial transplantation <u>a</u>nd robust engraftment (STAR) and genetic engineering. In summary, ^*^PDX AML models were generated from only around one-fifth of primary AML samples.

### *In vivo* characteristics of ^*^PDX AML models

^*^PDX AML models showed re-engraftment in up to 200 mice, up to 18 consecutive passages and thereby *in vivo* growth for up to 800 consecutive days until today, indicating the presence of non-exhausting leukemia-initiating cells (Table S1; Figure S2B,C). The passaging time was highly heterogeneous between and within models, with median passaging times ranging from 42 to 105 days, dependent on the passage number, and other factors in individual models (Figure S2D,E). The positive engraftment rate (PER) remained constant over sequential passaging and was independent of the number of injected cells, use of freshly isolated versus thawed cells, or recipient mouse age for most models (Figure S2F). On purpose, we included mice of both sexes into our studies, to prevent gender bias and ensure ethical animal use. While genetic engineering mildly affected PER, mouse gender had a more pronounced effect with higher PER in female than in male mice in some models, in line with published data (Figure S2F,G)^26^.

The amount of PDX cells that could be re-isolated from BM or spleen at an end-stage leukemia was highly heterogenous within and across ^*^PDX AML models; splenic engraftment associated with splenomegaly was observed in a subfraction and at late disease stages (Figure S2H), while we could not define any factors predicting splenic involvement (data not shown). Percentage of PDX cells in PB at advanced-stage leukemia differed between the models (Figure S2I). A particular advantage of the model is that, upon careful observation, most experiments can be terminated with mice showing no or mild symptoms (Figure S2J).

### ^*^PDX AML models originate from patients with highly aggressive disease

^*^PDX AML models might represent a biased subfraction of AML in patients; thus, we next examined variables associated with these models. As for primary engraftment (Figure S1A), disease status emerged as the most robust predictor for the ability to generate ^*^PDX AML models (Figure 2A and S3A) accordingly, patients whose cells gave rise to a ^*^PDX AML model showed dismal OS (Figure 2B and S3B) and thus represent a population of high need for better treatment. Most PDX models originating from ID samples failed to fulfill the prerequisites of score 5/6 PDX models, with four exceptions: AML-1177 and AML-1186, as well as AML-388 and AML-1226. The latter were generated from patients with primary refractory disease and are counted as “R/R samples”. Disease stage of AML-955 is unknown. Therefore, while around 60% of score 2-4 PDX models originated from ID, about 90% of score 5/6 PDX models originated from R/R disease (Figure S3A). The 23 ^*^PDX AML models were retrieved from patients of all age groups and from both sexes and originated from five distinct AML World Health Organization (WHO) subgroups at representative proportions: 8/23 models harbor *NPM1* mutations (30-35% of AML patients), 6/23 display a myelodysplasia-related phenotype (10-15% of AML cases) or have *KMT2A* rearrangement (KMT2Ar; about 5% in adults, 20-25% in children, respectively), and one model has a BCR::ABL1 fusion, which is also rare in patients (Figure 2C)^30-34^. As another peculiar feature, AML-491 and AML-661 are paired models derived from cells from the same patient collected at first and second relapse, respectively, allowing to study specific questions such as tumor recurrence. Interestingly, upon second relapse, a gain of an EZH2 LOF mutation was detected in this model (Table S3: Reiter 2018).

**Figure 2.**
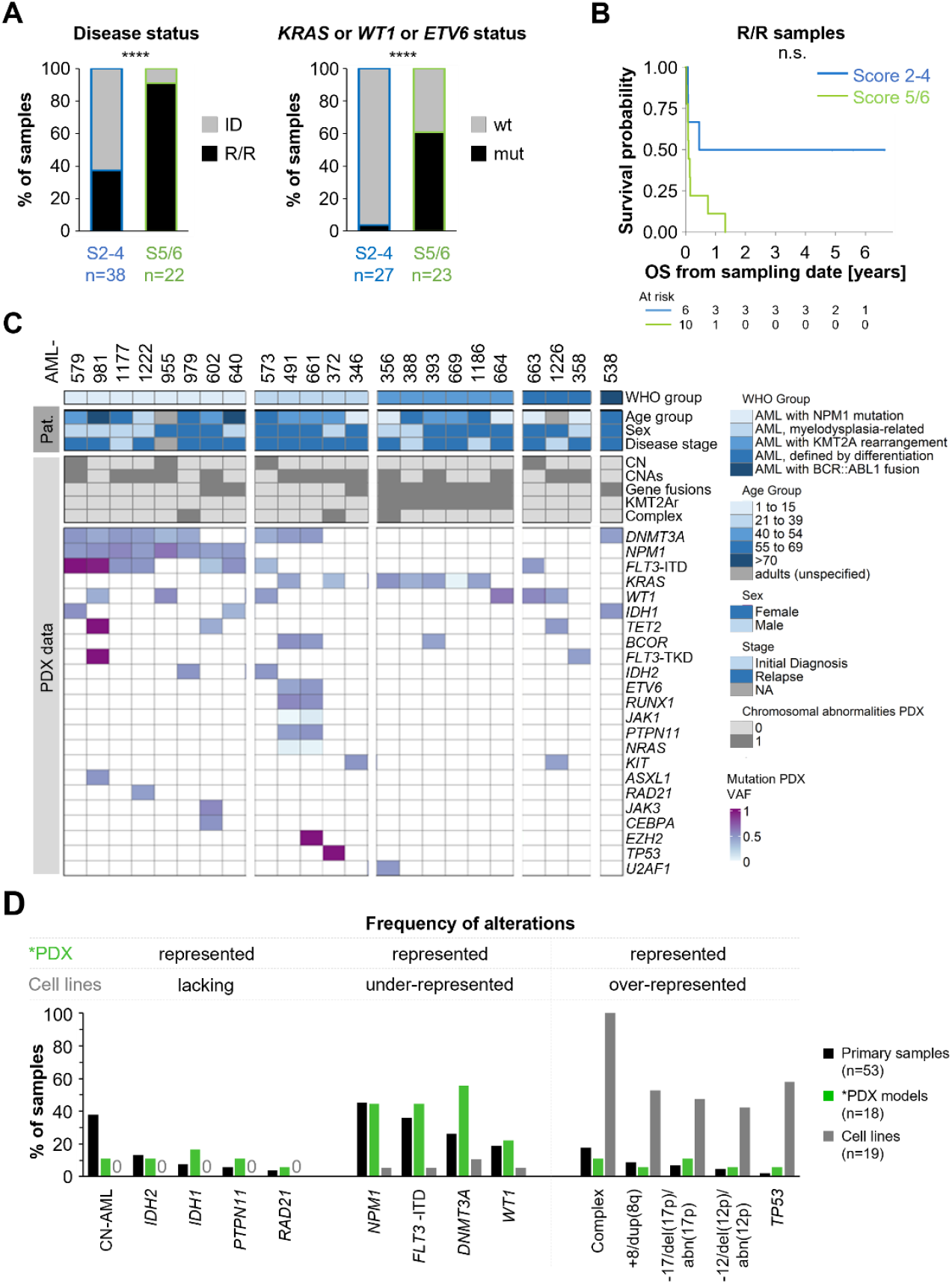
Genomic characteristics of ^*^PDX AML models better resemble primary AML cells than cell lines. **A** Prediction model for score 5/6 (n=23) versus Score 2-4 (n=46) was calculated by backward selection based on the Akaike information criterion, including samples of cohort 1-3. Left: percentage of models originating from initial diagnosis (ID, grey) or primary refractory / relapse (R/R, black) patient samples are depicted (*p=*8.7E-4). Right: percentage of models with no (grey) or detected mutation in *KRAS* or *WT1* or *ETV6* (black) are depicted (*p=*1.8E-05). The *p*-values were calculated with Fisher’s exact test. **B** Kaplan-Meier graph showing the survival probability of patients with R/R disease whose cells produced score 5/6 PDX models (green) or did not meet the criteria of high score models (score 2-4, blue). Log-rank test, *p*=0.053. **C Genomic landscape of** ^**\***^**PDX models**. ^*^PDX AML models are presented and categorized according to their WHO group. Patient (Pat.) data indicate age group, sex and disease status at time of sample collection. PDX models were characterized by cytogenetics, low coverage whole genome sequencing, RNA sequencing, methylation profiling, and DNA panel sequencing. PDX data depict cytogenetically normal (CN) models, detection of copy number alterations (CNAs), gene fusions, *KMT2A* rearrangements, and complex karyotype, as identified with at least two independent methods. Recurrent mutations are presented by variant allele frequency (VAF). **D** Relative frequencies of samples/models with specific mutational or chromosomal alterations are depicted for primary adult patient samples at relapsed disease of BEAT cohort (n=53, black), ^*^PDX AML models generated from adult patients (n=18, green), and AML cell lines ^36^ generated from adult patients and having more than 500 hits in Google Scholar (n=19, grey). Events are included which are present in a minimum of 5% in one of the three cohorts and are found in primary patients. Alterations are categorized according to prevalence in cell line models and sorted according to relative frequencies in patients. Refer to Figure S3C for further categories.

### Genomic characteristics of ^*^PDX AML models replicate AML biology better than cell lines

Molecular alterations within serially passaged ^*^PDX AML cells were studied using whole genome sequencing, panel sequencing, RNA sequencing, cytogenetics, methylation profiling and fluorescence in situ hybridization (Table S1).

At the level of genomics and in contrast to primary engraftment, mutations in *WT1, KRAS* or *ETV6*, but not in *FLT3-*ITD or *BCOR*, were associated with an increased likelihood to generate ^*^PDX AML models. 60% of score 5/6 models harbor mutations in these genes compared to 5% of score 2-4 PDX models, although the presence of *WT1, KRAS* or *ETV6* mutations does not correlate with patient OS probability (Figure 2A and Figure S3C,D).

Our ^*^PDX AML models exhibited a wide range of chromosomal alterations and/or mutations known to be associated with AML. Transcriptome analysis revealed both known fusions and undescribed fusion genes (Figure 2C and S3E,F; Table S1).

We compared the frequency of alterations between primary AML cells, PDX models, and cell lines. In the initial analysis, we included primary AML samples from adult patients at relapse from the BEAT cohort (n=53)^35, *^PDX AML models from adults (n=18), and AML cell lines from adults which were most frequently used in research (n=19)^36^. Importantly, cytogenetically normal samples and mutations in *IDH1, IDH2, PTPN11* and *RAD21* can be modelled in ^*^PDX AML models but are absent in AML cell lines or in most frequently applied cell lines, respectively. Additionally, mutations most prevalent in primary patient cells – *NPM1, FLT3-* ITD, *DNMT3A* and *WT1* – are recapitulated in PDX models at similar frequencies but are severely underrepresented in cell lines. Conversely, cell lines have a strong bias towards *TP53* mutations and complex karyotypes, while primary samples and ^*^PDX AML models display these alterations at low frequencies (Figure 2D and S3H,I). Several alterations were represented by both ^*^PDX AML models and cell lines, with ^*^PDX AML models overrepresenting *KRAS* mutations and *KMT2A* rearrangements. Some rare AML mutations were not found in ^*^PDX AML models, explainable by the limited number of primary samples studied. Unfortunately, some mutations are still not represented, neither in ^*^PDX AML models nor in cell lines, e.g. biallelic *CEBPA, SF3B1* or *SRSF2*, although for the latter we were able to establish a serially transplantable, although less robust PDX model of score 3 (Figure S3G-I; Table S1).

When we further broadened the analyses by including additional primary patient data sets^37,38^, scores 3-6 AML PDX models derived from adult patients (n=45), and all adult AML cell lines (n=54)^36^, we could reproduce the results described above and additionally found that the number of AML-specific mutations was similar in ^*^PDX AML models and primary samples (Figure S3I,J). In summary, PDX models were clearly superior to cell lines in mimicking the genetic and chromosomal alterations found in primary AML cells, including alterations for which no cell lines exist.

### Methylome, transcriptome and surface protein characteristics show that ^*^PDX AML models resemble primary AML cells

To further investigate the molecular classification of our models, we compared DNA methylation and gene expression (GE) patterns from ^*^PDX AML models (Table S1) with available own data from AML patient samples at initial diagnosis (AML-CG, n=433)^39,40^ and from AML cell lines (n=6). In the t-distributed Stochastic Neighbor Embedding (t-SNE) plots, PDX cells predominantly segregated from primary samples in both methylome and transcriptome (Figure 3A and S4A). Of note, both methylation and GE of AMLCG samples were conducted from bulk samples with variable percentage of blasts, while pure AML cells of PDX models were analyzed; furthermore, the analyses were not available from patient cells originating from relapsed disease. Interestingly, most differentially expressed pathways were associated with immunity, possibly induced by the immunocompromised environment of ^*^PDX AML models (Figure S4B-D).

**Figure 3.**
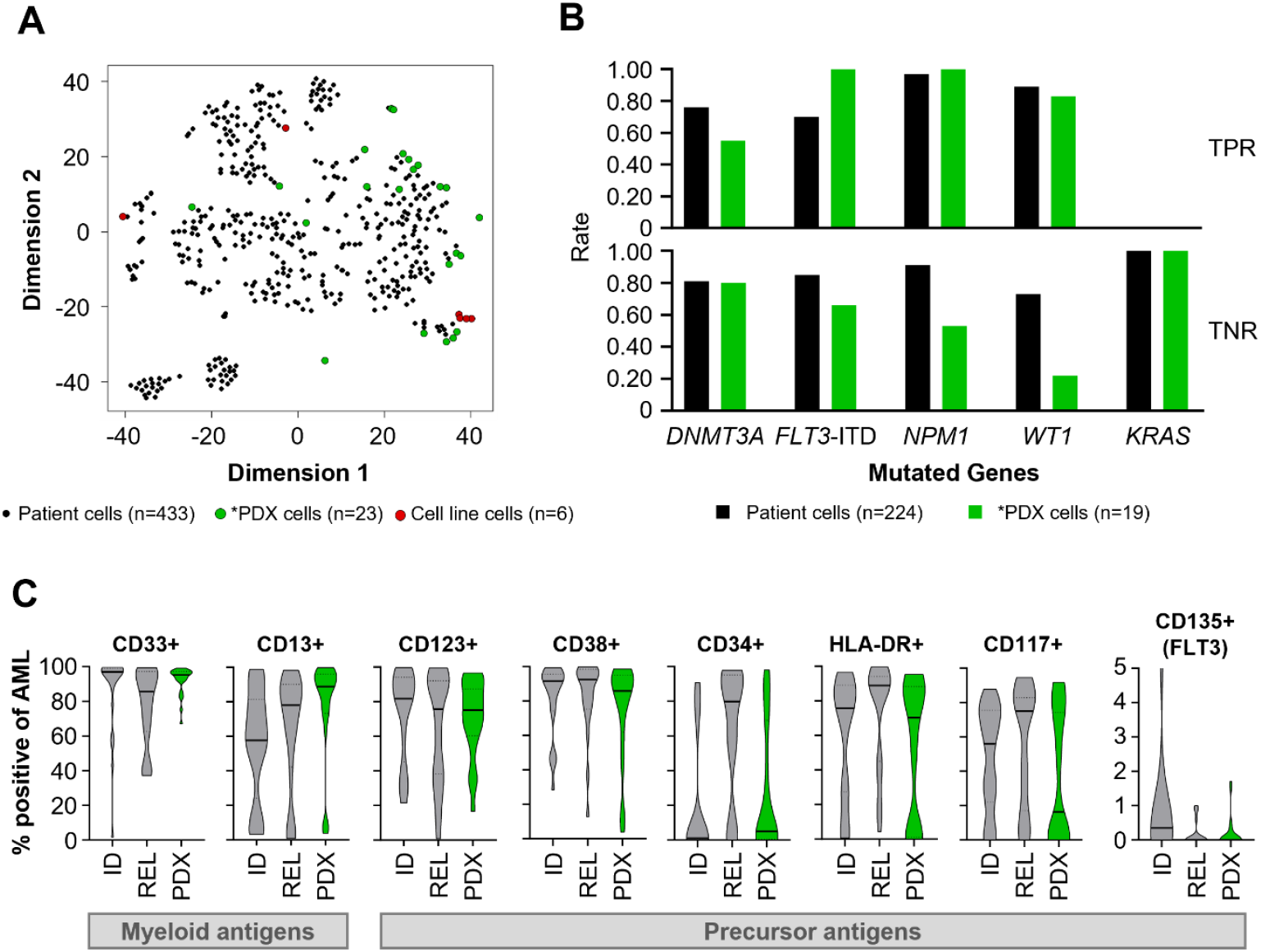
Methylome, transcriptome and surface protein characteristics show that ^*^PDX AML models resemble primary AML cells. **A t-SNE plot** showing the methylation profile of primary patient samples at ID (209 patients from the AML-CG-1999 or AML-CG-2008 study and 224 patients from the AML-CG Registry (DRKS00020816), black), ^*^PDX AML models (n=23, green), and AML cells lines (Leibniz Institute DSMZ, n=6, red). **B Prediction Analysis of Microarrays (PAM) models**. PAM classifiers based on DNA methylation profiling evaluated on AML-CG Registry patient samples (n=224) and ^*^PDX AML models (n=19). Displayed are the true positive rate (TPR) and true negative rate (TNR) for genes mutated in at least 20% of PDX samples. **C Immunophenotype of** ^**\***^**PDX AML models resembles primary patient samples**. Primary patient samples at initial diagnosis (ID, n=20), primary patient samples at relapse (REL, n=20), and ^*^PDX AML models (n=23) were analyzed for surface molecule expression via flow cytometry. Primary patient samples are independent of PDX models. Violin plots depicting the percentage of antigen-positive cells within blast gate; median indicated by solid line and 25^th^ and 75^th^ percentile by dotted lines.

We performed Prediction Analysis for Microarrays (PAM) with five genes mutated in at least 20% of ^*^PDX AML models. The predictive outcomes were largely comparable between patient samples and PDX models, with no consistent trend favoring either cohort in terms of model performance, highlighting a major similarity between ^*^PDX AML models and primary AML cells (Figure 3B and S4E). Additionally, ^*^PDX AML models maintained key immunophenotypic characteristics of primary AML cells, expressing myeloid and precursor antigens at heterogeneous levels and in similar percentages across all cohorts (CD33, CD123 and HLA-DR) or more biased towards samples from initial diagnosis (CD34 and CD33/CD7) or relapse (CD135 (FLT3) and CD33/CD64) (Figure 3C and S4F). In summary, PDX models largely mimic primary patient AML cells on the level of transcriptome, methylome and immunophenotype.

### ^*^PDX AML models allow a wide range of *ex vivo* and *in vivo* experiments

Until today, ^*^PDX AML models have been successfully used for a variety of applications, including functional and molecular assays and treatment trials, and were included in 45 scientific publications from collaboration partners across the globe and us (Figure 4A and S5A-C; Table S3). To further broaden their scientific use, these models and related characteristics will be shared via the openly accessible repository CancerModels.org^41^.

**Figure 4:**
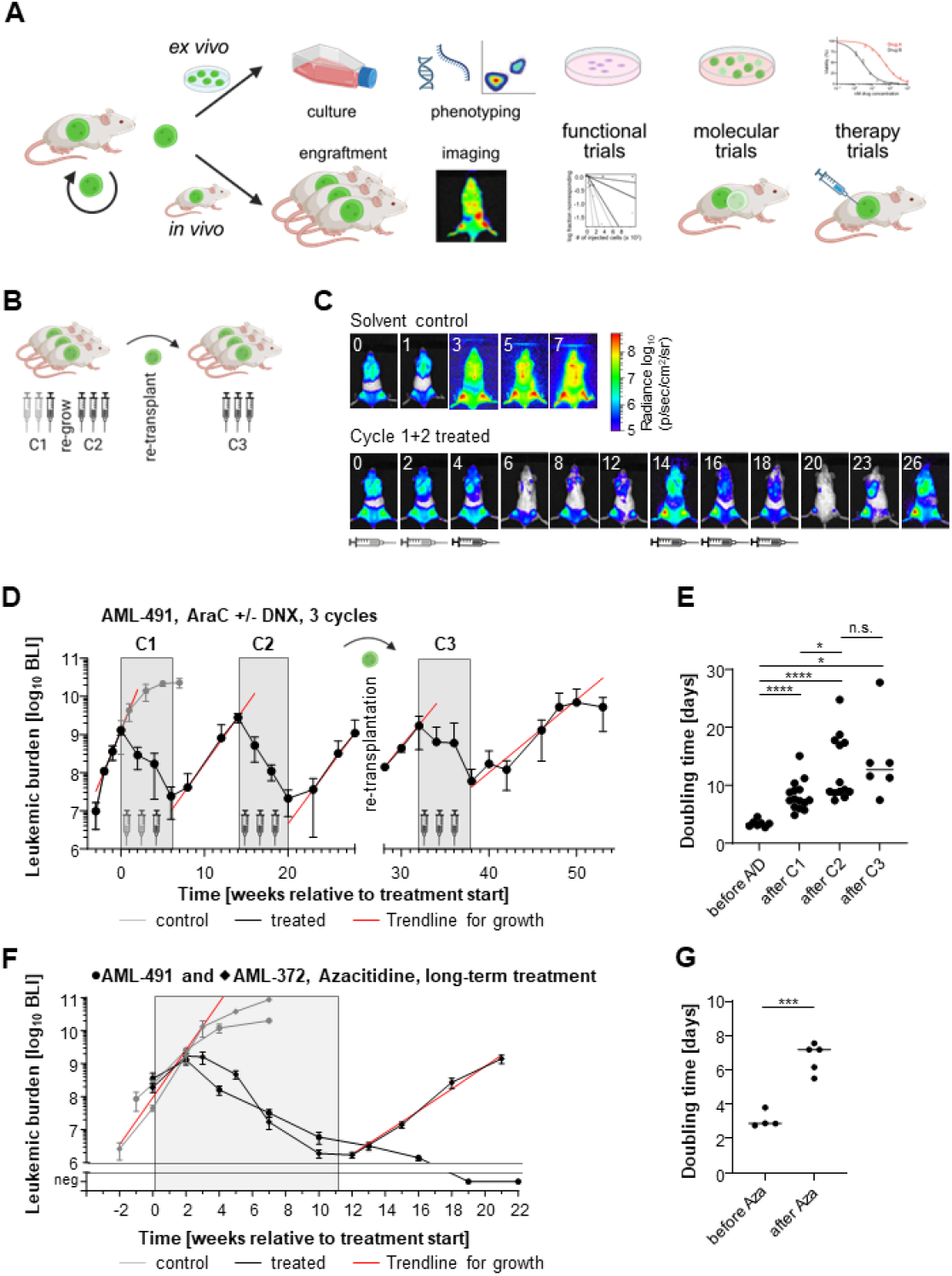
^*^PDX AML models allow a wide range of *ex vivo* and *in vivo* experiments, including long-term *in vivo* treatment trials. **A Scheme** illustrating published *in vivo* and *ex vivo* trials conducted with ^*^PDX AML models. **B-E Repetitive treatment to model induction plus consolidation chemotherapy**. AML-491 PDX cells were injected into mice and tumor burden was monitored by BLI. At a total flux of around 1E9 P/sec, mice were either treated with PBS (n=7, grey) or two cycles of chemotherapy (C1 and C2, n=15, black). After re-growth, animals were sacrificed and PDX cells were re-injected into next recipient mice receiving a third cycle of chemotherapy (n=6, C3). **B** Scheme. Grey syringe: AraC 50 mg/kg, d1-4 + DNX 1mg/kg, d1+4, 10 days rest; black syringe: AraC 100 mg/kg, d1-4, 10 days rest. **C** BLI pictures of two exemplary mice are shown; numbers indicate weeks after treatment start. **D** Quantification of BLI. Grey box indicates treatment period, and red lines show exponential trend line during growth. **E** Growth rate, measured by increase in BLI signal, is depicted for untreated mice (before A/D, n=7), after stop of 1^st^ (C1, n=15), 2^nd^ (C2, n=13) or 3^rd^ (C3, n=6) cycle of chemotherapy. Dots indicate values of individual mice, line indicates median. n.s. not significant, ^*^ *p*<0.05, ^****^ *p*<0.0001 as tested by two-sided unpaired t-test with Welch’s correction. **F**,**G Long-term therapy with Azacitidine**. AML-372 (n=4-5 per group, diamonds) or AML-491 (n=4 per group, circles) PDX cells were injected into mice and tumor burden was monitored by BLI. Mice were either treated with PBS (grey) or Azacitidine (black) for 11 weeks. **F** Quantification of BLI. Grey box indicates treatment period, and red lines show exponential trend line during growth. **G** Growth rate, measured by increase in BLI signal, is depicted for untreated mice (before Azacitidine, n=4) or after stop of Azacitidine therapy (n=5). Dots indicate values of individual mice. ^***^ *p*<0.001 as tested by two-sided unpaired t-test with Welch’s correction.

PDX cells allow functional characterization of leukemia cells and investigation of different tumor cell subpopulations. In AML primografts and with the help of secondary transplantation assays, the cancer stem cell concept had been elucidated^13^. Our serially transplantable models enable stem cell-associated tests such as limited dilution transplantation assays to determine the frequency of leukemia-initiating stem cells^17,42^ (Table S3: Tahk 2021, Ghalandary 2023, Bahrami 2023, Zhou 2023). Moreover, they also facilitate functional assays allowing to investigate homing (Table S3: Bahrami 2023), (re-) engraftment capacity (Table S3: Yankova 2021, Zhou 2023, Able 2025) and proliferation status^42^ of leukemia cells (Figure 4A and S5A).

^*^PDX AML cells freshly isolated from donor mice display a higher viability than frozen/thawed cells, allowing complex *ex vivo* studies with high cell numbers and over several weeks (Figure S5E). However, *ex vivo* studies could only be performed with PDX cells amplified in donor mice, as ^*^PDX AML models show only minor proliferation rates *ex vivo*.

For cutting edge *in vivo* experiments, ^*^PDX AML models were genetically engineered to stably express enhanced firefly luciferase for sensitive and repetitive monitoring of tumor burden using BLI^17,43^. For molecular studies, derivative ^*^PDX AML models were generated which express various transgenes, such as fluorochromes, Cre-ER^T2^ or Cas9. This allows fluorescent labelling for cell enrichment^42^, genetic barcoding to track (single cell) clones (Table S3: Zeller 2022), (inducible) knock-down (Table S3: Garg 2019, Jensen 2020, Carlet 2021, Sun 2021, Sollier 2025), CRISPR/Cas9 knock out (Table S3: Salik 2020, Ghalandary 2023, Bahrami 2023, Barbosa 2024), marker-guided competitive *in vivo* assays (Table S3: Carlet 2021, Zeller 2023, Ghalandary 2023, Bahrami 2023) and CRISPR/Cas9 *in vivo* and *in vitro* screens (Table S3: Ghalandary 2023, Bahrami 2023) (Figure S5B,D).

Serially transplantable, luciferase-transgenic PDX models are particularly suited to perform comprehensive pre-clinical *in vivo* therapy trials. Importantly, in-depth characterization allows the selection of samples representing specific patient subgroups which might benefit most from specific (targeted) drugs; for example, *FLT3*-ITD positive samples were specifically selected for testing a novel multi-kinase inhibitor (Table S3: Lopez-Millan 2022). The value of our PDX models has also been recognized by the leukemia community and sparked numerous collaborations with research groups from Germany, Europe and across the globe focusing on the development of new treatment options. In the last years, multiple therapeutic strategies were tested in our PDX models including targeted therapies (Table S3: Garz 2017, Lynch 2019, Janssen 2022, Lopez-Millan 2020, Yankova 2021, Shi 2025), immunotherapies (Table S3: Gottschlich 2023, Tahk 2021, Baroni 2020) and antibody-based therapies (Table S3: Krupka 2015, Roas 2022, Augsberger 2022, Able 2025; Figure S5C).

### Long-term *in vivo* experiments show reduced growth rates after treatment

As a concrete example of their application, we show here that ^*^PDX AML models and repetitive *in vivo* imaging enable long-term *in vivo* treatment trials and thus provide new insights into the disease biology. Spontaneous tumor growth cannot be studied in patients as immediate and continuous treatment is required upon cancer diagnosis. The influence of treatment on spontaneous growth rates remains largely elusive in leukemia, while certain solid tumors showed accelerated growth after chemotherapy^44,45^.

Most treatment regimens are based on repetitive therapy cycles or constant drug application over months. These schedules are difficult to reproduce in current experimental models, among others due to the aggressiveness of cell line models or the lack of options for continuous tumor monitoring. Using an adapted form of a mouse equivalent to the “7+3” induction therapy in patients^46,47^, we mimicked induction- and consolidation therapy and treated mice transplanted with luciferase-transgenic ^*^PDX AML-491 cells with repetitive cycles of chemotherapy (Figure 4B-E). Of note, NSG mice only tolerate low-doses of DNA-damaging agents due to their *scid*-background^47^. We have previously shown that PDX models differed in their response towards a 3 week-treatment with Cytarabine and DaunoXome, a liposomally encapsulated version of the anthracycline Daunorubicin, which is currently not commercially available. While therapy only had a minor effect on tumor burden in AML-393 and AML-661, AML-491 showed response (Table S3: Stief 2020, Kempf 2021, Zeller 2022). Therefore, we chose this model to simulate induction and consolidation therapy. Mice were transplanted with PDX AML-491 cells and treated with two cycles of chemotherapy (Figure 4B). While solvent-treated control mice rapidly progressed to advanced leukemia as monitored by BLI, the treatment significantly reduced tumor burden but – similar to what is commonly observed in the clinics – tumor re-growth occurred upon therapy cessation, both after first and second treatment cycle (Figure 4CD). PDX cells were isolated in the logarithmic growth phase, around 30 weeks after cell transplantation, and re-transplanted due to mouse aging into secondary recipients which received a third treatment cycle (Figure 4B). Repeatedly, the tumor load was effectively reduced with re-occurrence after stopping treatment (Figure 4D). Despite expected reduction in blood cell counts^48^, the treatment was well tolerated over prolonged periods of time (Figure S5F,G). After a total of 53 experimental weeks with nine blocks of chemotherapy in three cycles, we were surprised to find that response to treatment remained unchanged, with no signs of acquired resistance, while tumor growth rate decreased continuously with each treatment block (Figure 4D,E and S5H). Importantly, only continuous imaging allowed to capture this additional, essential piece of information.

Unfit AML patients are typically treated with hypomethylating agents^49^. Treatment of ^*^PDX AML-491 cells with Azacitidine monotherapy for three months reduced tumor burden below the imaging detection limit, while mice carrying ^*^PDX AML-372 showed tumor re-growth after end of treatment with an increase in BLI doubling time, indicating that a reduced growth rate after treatment represents a shared feature across different treatment modalities and PDX models (Figure 4F,G and S5I,J).

In summary, we established ^*^PDX AML models - serially transplantable and robust genetically engineered PDX AML models - for translational and preclinical trials, harboring the potential to foster future AML research for the ultimate benefit of patients suffering AML.

## Discussion

To advance cancer research, we established and characterized ^*^PDX models as a sharable tool according to PDX-MI guidelines^50^. They fulfill clearly defined criteria regarding the ability of serial transplantation, robust engraftment and genetic engineering. ^*^PDX AML models represent alterations found in primary AML patient cells, both at initial diagnosis and relapse, and include subgroups that cannot be modelled with cell lines, such as cytogenetically normal and *IDH1/2*-mutant AML. They are used for complex molecular and functional studies, both *ex vivo* and *in vivo*, and are especially suitable for long-term preclinical treatment trials.

^*^PDX AML models complement existing model systems such as primary tumor cells, primografts, and cell lines. Primary patient cells are typically available mostly in limited numbers, especially from pediatric patients and in certain countries, complicating data reproducibility and standardization of experiments. Primary patient cells and primografts don’t consistently grow *ex vivo*, while it will be interesting to study whether PDX AML cells do so without major skewing. Compared to primary patient samples, primografts undergo clonal selection with a bias towards “relapse-committed” clones^16,51-53^, complicating co-clinical trials for individual patients. Additionally, clones might be further enriched during serial passaging^21^.

Nevertheless, our ^*^PDX AML models, even from advanced passages and after genetic engineering, showed high overall concordance with primary patient samples regarding mutational landscape, chromosomal alterations, and methylation patterns. As shown before, neither serial passaging nor genetic engineering had a major impact on characteristics of ^*^PDX AML models, including mutations, transcriptome, or therapy response^17,54,55^. Importantly, we do not anticipate that each PDX model represents the corresponding primary patient sample. Instead, we have established robust AML models representing various specific disease subgroups, facilitating scientific discoveries relevant for groups of patients, not for individual patients.

One limitation of our cohort is that it only includes samples collected from patients in Europe and North America, thus introducing an ethnic bias. Furthermore, ^*^PDX-AML models are biased towards high-risk subgroups. Primary leukemia cells from AML subtypes with less dismal prognosis showed positive primary engraftment in immune compromised mice only after long latency and at low percentage^22^, making it impossible to develop ^*^PDX models as defined here. Still, generating robust PDX models from highly aggressive samples represents a valuable tool to model the clinically most challenging subtypes of the disease with most urgent need for new treatment options. The advantage of ^*^PDX AML models lies in their ability to provide highly patient-related cells repetitively and reproducibly for a wide variety of different AML subtypes and clones across the broad heterogeneity of AML.

Compared to cell lines, PDX models are time- and resource-intensive and logistically complex^56,57^. Nevertheless and importantly, PDX AML models harbour major advantages over cell lines which include (i) orthotopic engraftment due to a persistent dependence on the physiologic tumor microenvironment^42,58^; (ii) maintained stem cell hierarchy^17,59,60^; (iii) higher similarity to primary AML cells; (iv) availability of AML subtypes for which no cell lines exist; and (v) improved predictive value for drug responses^48 61^.

^*^PDX models allow performing prolonged *in vivo* treatment trials. Repetitive *in vivo* imaging facilitates close monitoring of tumor burden under treatment and thereby enabled, to our knowledge, the first preclinical *in vivo* trial in acute leukemia which continued for 12 consecutive months. The technical advance allowed studying the kinetics of tumor repopulation after treatment, a topic that was often neglected so far, but is relevant to understand real world treatment outcomes^45^. In contrast to findings in solid tumors, where tumor growth was accelerated after chemotherapy^44,45^, we observed slower growth after treatment. Our results are consistent with the well-known fact that chemotherapy is more effective in proliferating cells than in resting cells^42,62^. Reduced tumor growth kinetics might be explained by efficient eradication of proliferating cells and exhaustion of cells in the stem cell compartment. However, these results need to be validated in additional PDX models and future projects are required to uncover the underlying mechanisms. Of relevance for clinical decision making, our data show that AML growth dynamics might substantially differ directly after treatment compared to disease relapse.

In summary, ^*^PDX AML models mimic the human disease more closely than cell lines and can be used in preclinical studies to generate data for subsequent clinical trials across multiple genetically distinct AML subtypes. Our vision is that ^*^PDX AML models will be globally shared analogous to cell lines, contribute to scientific progress in the field and ultimately help improving the prognosis of patients suffering from AML.

## Materials and Methods

### Ethical Statement

BM and peripheral blood (PB) samples from AML patients were collected for diagnostic purposes. Signed informed consent was obtained from the patients or from person having custody in case patients were minor. The study was performed in accordance with the Helsinki Declaration of 1975, as revised in 2013, and with the ethical standards of the responsible committees on human experimentation (Research Ethics Boards of the medical faculty of LMU Munich, #068-08, 222-10 and 20-739).

### Mouse work

Animal trials were performed in accordance with the ethical standards of the official committee on animal experimentation and according to ARRIVE guidelines^63^. Five to 35 weeks old male and female NOD.Cg-Prkdc^scid^ IL2rg^tm1Wjl^/SzJ (NSG) mice (The Jackson Laboratory, Bar Harbour, USA) were included (n=2111); these were part of diverse scientific projects (Table S3). To reduce gender bias and ensure ethical animal use, we included both female and male mice and both younger and older mice in our studies^64^. This approach minimizes the number of animals bred without scientific purpose, in line with German regulations that prohibit the killing of animals without justified scientific need. No randomization or blinding was performed. All mice, mouse-related interventions and values were documented in a customized database (My IMouse, Bioslava GbR, Berg, Germany).

### Transplantation of primary leukemia cells into NSG mice

Depending on availability of cells, 2.5E5 to 1E7 fresh or frozen/thawed primary cells were injected into the tail vein of one to eight NSG mice. In contrast to literature^15,16,22,65^ and to avoid treatment-induced bias, mice were not pre-irradiated prior cell injection. No enrichment or depletion steps were performed. Exclusion criteria were samples from EDTA or citrate anti-coagulated material, samples with less than 25% blasts, samples with less than 2.5E5 cells, and acute promyelocytic leukemias. Information on samples is listed in Table S1 according to PDX-MI guidelines^50^.

### Functional scoring system of PDX cells

PDX models were categorized into classes (Score 1 – 6) according to functional engraftment characteristics. Score 1: no primary engraftment. Score 2: primary, but no secondary engraftment. Score ≥3: serial positive engraftment; positive engraftment after freeze-thawing cycles; engraftment primarily in BM and spleen. Score ≥4: detection of >80% human cells in BM at end-stage leukemia; positive engraftment in at least 75% (Score 4 and 5) or at least 90% (Score 6) of transplanted mice; median time from transplantation to end-stage leukemia below 110, 90 or 70 days (Score 4, 5 or 6, respectively). Score ≥5: successful genetic engineering. Score 6: transduction rates above 5%.

### Therapy trials

#### Repetitive treatment cycles with cytarabine (AraC) and DaunoXome (DNX)

AML-491 PDX cells were injected into groups of mice (5E5 luc^+^ cells/mouse). Tumor burden was regularly monitored by bioluminescence *in vivo* imaging (BLI). When BLI signals reached a total flux of around 1E9 Photons/second, mice were either treated with PBS (n=7) or first cycle of chemotherapy (C1) (n=15). Treated mice showed a reduction of tumor burden followed by a re-growth of tumor cells and were again treated with a second cycle of chemotherapy (C2). When BLI signals again reached a total flux above 1E9 Photons/second, or when mice showed clinical signs of illness, the animals were sacrificed. PDX cells were re-isolated, counted, and 5E6 cells were re-injected into next recipient mice. These mice received a third cycle of chemotherapy (C3) (n=6). Chemotherapy schedule: per cycle three repetitions with four days treatment and ten days rest; C1, week 1 and 3: AraC 50 mg/kg d1-4, DNX 1 mg/kg d1+4; cycle 1, week 5: AraC 100 mg/kg d1-4; C2 and C3, week 1, 3 and 5: AraC 100 mg/kg, d1-4. AraC (Cell Pharma GmbH, Bad Vilbel, Germany) was diluted in PBS and applied i.p., DNX (Galen Ltd, Craigavon, United Kingdom) was diluted in distilled H_2_O and applied i.v.

#### Long-term treatment with Azacitidine (Aza)

AML-372 (n=3 per group) or AML-491 (n=4 per group) PDX cells were injected into groups of mice (3E5 luc+ cells/mouse). Tumor burden was regularly monitored by BLI. When BLI signals reached a total flux of around 1E8 Photons/second, mice were treated i.p. either with PBS or Azacitidine (2.5 mg/kg or 5 mg/kg, diluted in PBS) three times a week for 11 weeks. Monitoring of tumor burden was continued after end of therapy.

### Enrichment of AML PDX cells for sequencing

AML PDX cells were enriched from murine BM via negative selection by magnetic cell sorting using mouse cell depletion kit (Miltenyi Biotech, Bergisch Gladbach, Germany). 100 μl MicroBeads were used per 1E7 BM cells isolated from mice with end-stage leukemia. Enriched PDX cells were lysed in RLT-Buffer (Qiagen, Hilden, Germany) with 1% beta-mercaptoethanol.

### Prediction models

To develop a prediction model for positive engraftment, a backward selection based on the Akaike information criterion was performed. Variables were included that had less than 10% of missing values and a univariable *p*-value of ≤0.1.

### Statistics

Statistical analyses were calculated using GraphPad Prism 10 software (Graphpad Prism, La Jolla, CA, USA) and R^66^. Statistical tests used are indicated in the figure legends. No star: not significant; ^*^ *p*<0.05, ^**^ *p*<0.01, ^***^ *p*<0.001, ^****^ *p*<0.0001.

**Further methods** are provided in the supplementary information.

## Acknowledgements

We thank all team members of the Irmela Jeremias lab for helpful discussions and critically reading the manuscript; Maike Fritschle, Zahra Raffie Pour, Chantal Jekien (all Irmela Jeremias lab) and Bianka Ksienzyk (Department of Medicine III, LMU Munich) for excellent technical assistance; Stephanie Hoffmann for lab management; Markus Brielmeier and team (Research Unit Comparative Medicine) for animal care services; Véronique de Mas and François Vergez (Diagnosis laboratory, University of Toulouse) and the team of the leukemia diagnostics lab (LFL, Department of Medicine III, LMU Munich) for providing primary patient cells and diagnostics; PRoXe (Weinstock Laboratory, Dana-Farber Cancer Institute) for providing PDX models; Johannes Bagnoli and Wolfgang Enard (Faculty of Biology, LMU Munich) for primeSeq of ^*^PDX models, Marina Vogel and Andrea Weber (Genomics & Proteomics Core Facility, German Cancer Research Center (DKFZ)), and Stefan Krebs, Helmut Blum and Alexander Graf (Laboratory for Functional Genome Analysis, LMU Munich) for providing excellent sequencing services; Isolde Summerer and Claudia Haferlach (MLL Münchner Leukämielabor GmbH) for cytogenetic characterization of ^*^PDX samples.

The work was supported by grants from the European Research Council (Consolidator Grant 681524) and the Deutsche Forschungsgemeinschaft (DFG, German Research Foundation) – 537701335 (Reinhart Koselleck project) and SFB 1709/1 2025 – 533056198; a Mildred Scheel Professorship by German Cancer Aid; Bettina Bräu Stiftung and Dr. Helmut Legerlotz Stiftung (all to IJ) and by funding from the Deutsche José Carreras Leukämie-Stiftung (DJCLS 15 R/2021 and DJCLS 02 R/2023 to I.J. and P.M.)

## Author contribution

B.V., V.J., G.H., A.M., L.H., L.M., F.K., and A.F. performed experiments; B.V., V.J., K.K., L.Lagally, L.Laszko, C.A., M.R.T., A.M.D., S.S., K.H.M., P.A.G., M.S., S.V., and T.H. analyzed results; M.C.A., C.D.B., M.P.C., C.D., M.E., K.S.G., P.M., J.E.S., C.R., F.V., V.d.M., A.S., C.T., T.V.H., X.W., J.H.K., M.v.B.B., W.H., M.S., T.H., and K.S. provided material and data; B.V., K.K., and I.J. made the figures; B.V., T.H., K.S., and I.J. designed the research; B.V., K.K., and I.J. wrote the paper.

## References

1. Döhner H, Wei AH, Appelbaum FR, et al. Diagnosis and management of AML in adults: 2022 recommendations from an international expert panel on behalf of the ELN. Blood. 2022;140(12):1345–1377.

2. de Leeuw DC, Ossenkoppele GJ, Janssen J. Older Patients with Acute Myeloid Leukemia Deserve Individualized Treatment. Curr Oncol Rep. 2022;24(11):1387–1400.

3. Montesinos JJ, Sanchez-Valle E, Flores-Figueroa E, et al. Deficient proliferation and expansion in vitro of two bone marrow cell populations from patients with acute myeloid leukemia in response to hematopoietic cytokines. Leuk Lymphoma. 2006;47(7):1379–1386.

4. Stevens AM, Terrell M, Rashid R, et al. Addressing a Pre-Clinical Pipeline Gap: Development of the Pediatric Acute Myeloid Leukemia Patient-Derived Xenograft Program at Texas Children’s Hospital at Baylor College of Medicine. Biomedicines. 2024;12(2).

5. Ben-David U, Siranosian B, Ha G, et al. Genetic and transcriptional evolution alters cancer cell line drug response. Nature. 2018;560(7718):325–330.

6. Safa-Tahar-Henni S, Paez Martinez K, Gress V, et al. Comparative small molecule screening of primary human acute leukemias, engineered human leukemia and leukemia cell lines. Leukemia. 2025;39(1):29–41.

7. Sinha R, Luna A, Schultz N, Sander C. A pan-cancer survey of cell line tumor similarity by feature-weighted molecular profiles. Cell Rep Methods. 2021;1(2):100039.

8. Drexler HG. Isolation and Culture of Leukemia Cell Lines. In: Langdon SP, ed. Cancer Cell Culture: Methods and Protocols. Totowa, NJ: Humana Press; 2004:141–155.

9. Khan AO, Rodriguez-Romera A, Reyat JS, et al. Human Bone Marrow Organoids for Disease Modeling, Discovery, and Validation of Therapeutic Targets in Hematologic Malignancies. Cancer Discov. 2023;13(2):364–385.

10. Blanchard Z, Brown EA, Ghazaryan A, Welm AL. PDX models for functional precision oncology and discovery science. Nature Reviews Cancer. 2024.

11. Woo XY, Giordano J, Srivastava A, et al. Conservation of copy number profiles during engraftment and passaging of patient-derived cancer xenografts. Nat Genet. 2021;53(1):86–99.

12. Sun H, Cao S, Mashl RJ, et al. Comprehensive characterization of 536 patient-derived xenograft models prioritizes candidatesfor targeted treatment. Nat Commun. 2021;12(1):5086.

13. Bonnet D, Dick JE. Human acute myeloid leukemia is organized as a hierarchy that originates from a primitive hematopoietic cell. Nature Medicine. 1997;3(7):730–737.

14. Boyd AL, Reid JC, Salci KR, et al. Acute myeloid leukaemia disrupts endogenous myelo-erythropoiesis by compromising the adipocyte bone marrow niche. Nat Cell Biol. 2017;19(11):1336–1347.

15. Ishikawa F, Yoshida S, Saito Y, et al. Chemotherapy-resistant human AML stem cells home to and engraft within the bone-marrow endosteal region. Nat Biotechnol. 2007;25(11):1315–1321.

16. Shlush LI, Mitchell A, Heisler L, et al. Tracing the origins of relapse in acute myeloid leukaemia to stem cells. Nature. 2017;547(7661):104–108.

17. Vick B, Rothenberg M, Sandhofer N, et al. An advanced preclinical mouse model for acute myeloid leukemia using patients’ cells of various genetic subgroups and in vivo bioluminescence imaging. PLoS One. 2015;10(3):e0120925.

18. Culen M, Kosarova Z, Jeziskova I, et al. The influence of mutational status and biological characteristics of acute myeloid leukemia on xenotransplantation outcomes in NOD SCID gamma mice. J Cancer Res Clin Oncol. 2018;144(7):1239–1251.

19. Hassan N, Yang J, Wang JY. An Improved Protocol for Establishment of AML Patient-Derived Xenograft Models. STAR Protoc. 2020;1(3):100156.

20. Griessinger E, Vargaftig J, Horswell S, Taussig DC, Gribben J, Bonnet D. Acute myeloid leukemia xenograft success prediction: Saving time. Experimental Hematology. 2018;59:66-71.e64.

21. Kawashima N, Ishikawa Y, Kim JH, et al. Comparison of clonal architecture between primary and immunodeficient mouse-engrafted acute myeloid leukemia cells. Nature Communications. 2022;13(1).

22. Paczulla AM, Dirnhofer S, Konantz M, et al. Long-term observation reveals high-frequency engraftment of human acute myeloid leukemia in immunodeficient mice. Haematologica. 2017;102(5):854–864.

23. Rombouts WJC, Martens ACM, Ploemacher RE. Identification of variables determining the engraftment potential of human acute myeloid leukemia in the immunodeficient NOD/SCID human chimera model. Leukemia. 2000;14(5):889–897.

24. Sanchez PV, Perry RL, Sarry JE, et al. A robust xenotransplantation model for acute myeloid leukemia. Leukemia. 2009;23(11):2109–2117.

25. Mian SA, Ariza-McNaughton L, Anjos-Afonso F, et al. Influence of donor-recipient sex on engraftment of normal and leukemia stem cells in xenotransplantation. Hemasphere. 2024;8(5):e80.

26. Stanger AMP, Arnone M, Hanns P, et al. Recipient sex and donor leukemic cell characteristics determine leukemogenesis in patient-derived models. Haematologica. 2025.

27. Townsend EC, Murakami MA, Christodoulou A, et al. The Public Repository of Xenografts Enables Discovery and Randomized Phase II-like Trials in Mice. Cancer Cell. 2016;29(4):574–586.

28. Woiterski J, Ebinger M, Witte KE, et al. Engraftment of low numbers of pediatric acute lymphoid and myeloid leukemias into NOD/SCID/IL2Rcgammanull mice reflects individual leukemogenecity and highly correlates with clinical outcome. Int J Cancer. 2013;133(7):1547–1556.

29. Richter A, Roolf C, Sekora A, et al. The Molecular Subtype of Adult Acute Lymphoblastic Leukemia Samples Determines the Engraftment Site and Proliferation Kinetics in Patient-Derived Xenograft Models. Cells. 2022;11(1).

30. Grimwade D, Hills RK, Moorman AV, et al. Refinement of cytogenetic classification in acute myeloid leukemia: determination of prognostic significance of rare recurring chromosomal abnormalities among 5876 younger adult patients treated in the United Kingdom Medical Research Council trials. Blood. 2010;116(3):354–365.

31. Khoury JD, Solary E, Abla O, et al. The 5th edition of the World Health Organization Classification of Haematolymphoid Tumours: Myeloid and Histiocytic/Dendritic Neoplasms. Leukemia. 2022;36(7):1703–1719.

32. Ranieri R, Pianigiani G, Sciabolacci S, et al. Current status and future perspectives in targeted therapy of NPM1-mutated AML. Leukemia. 2022;36(10):2351–2367.

33. van Weelderen RE, Klein K, Harrison CJ, et al. Measurable Residual Disease and Fusion Partner Independently Predict Survival and Relapse Risk in Childhood KMT2A-Rearranged Acute Myeloid Leukemia: A Study by the International Berlin-Frankfurt-Munster Study Group. J Clin Oncol. 2023;41(16):2963–2974.

34. Vetro C, Haferlach T, Meggendorfer M, et al. Cytogenetic and molecular genetic characterization of KMT2A-PTD positive acute myeloid leukemia in comparison to KMT2A-Rearranged acute myeloid leukemia. Cancer Genet. 2020;240:15–22.

35. Bottomly D, Long N, Schultz AR, et al. Integrative analysis of drug response and clinical outcome in acute myeloid leukemia. Cancer Cell. 2022;40(8):850–864 e859.

36. Pettirossi V, Venanzi A, Spanhol-Rosseto A, et al. The gene mutation landscape of acute myeloid leukemia cell lines and its exemplar use to study the BCOR tumor suppressor. Leukemia. 2023;37(2):473–477.

37. Greif PA, Hartmann L, Vosberg S, et al. Evolution of Cytogenetically Normal Acute Myeloid Leukemia During Therapy and Relapse: An Exome Sequencing Study of 50 Patients. Clin Cancer Res. 2018;24(7):1716–1726.

38. Rausch C, Rothenberg-Thurley M, Dufour A, et al. Validation and refinement of the 2022 European LeukemiaNet genetic risk stratification of acute myeloid leukemia. Leukemia. 2023;37(6):1234–1244.

39. Moser C, Jurinovic V, Sagebiel-Kohler S, et al. A clinically applicable gene expression-based score predicts resistance to induction treatment in acute myeloid leukemia. Blood Adv. 2021;5(22):4752–4761.

40. Herold T, Jurinovic V, Batcha AMN, et al. A 29-gene and cytogenetic score for the prediction of resistance to induction treatment in acute myeloid leukemia. Haematologica. 2018;103(3):456–465.

41. Perova Z, Martinez M, Mandloi T, et al. PDCM Finder: an open global research platform for patient-derived cancer models. Nucleic Acids Res. 2023;51(D1):D1360–D1366.

42. Ebinger S, Zeller C, Carlet M, et al. Plasticity in growth behavior of patients’ acute myeloid leukemia stem cells growing in mice. Haematologica. 2020;105(12):2855–2860.

43. Terziyska N, Castro Alves C, Groiss V, et al. In vivo imaging enables high resolution preclinical trials on patients’ leukemia cells growing in mice. PLoS One. 2012;7(12):e52798.

44. El Sharouni SY, Kal HB, Battermann JJ. Accelerated regrowth of non-small-cell lung tumours after induction chemotherapy. Br J Cancer. 2003;89(12):2184–2189.

45. Kim JJ, Tannock IF. Repopulation of cancer cells during therapy: an important cause of treatment failure. Nat Rev Cancer. 2005;5(7):516–525.

46. Forsberg M, Konopleva M. AML treatment: conventional chemotherapy and emerging novel agents. Trends Pharmacol Sci. 2024.

47. Wunderlich M, Mizukawa B, Chou FS, et al. AML cells are differentially sensitive to chemotherapy treatment in a human xenograft model. Blood. 2013;121(12):e90–97.

48. Farge T, Saland E, de Toni F, et al. Chemotherapy-Resistant Human Acute Myeloid Leukemia Cells Are Not Enriched for Leukemic Stem Cells but Require Oxidative Metabolism. Cancer Discov. 2017;7(7):716–735.

49. DiNardo CD, Pratz K, Pullarkat V, et al. Venetoclax combined with decitabine or azacitidine in treatment-naive, elderly patients with acute myeloid leukemia. Blood. 2019;133(1):7–17.

50. Meehan TF, Conte N, Goldstein T, et al. PDX-MI: Minimal Information for Patient-Derived Tumor Xenograft Models. Cancer Res. 2017;77(21):e62–e66.

51. Ben-David U, Ha G, Tseng YY, et al. Patient-derived xenografts undergo mouse-specific tumor evolution. Nat Genet. 2017;49(11):1567–1575.

52. Eirew P, Steif A, Khattra J, et al. Dynamics of genomic clones in breast cancer patient xenografts at single-cell resolution. Nature. 2015;518(7539):422–426.

53. Klco JM, Spencer DH, Miller CA, et al. Functional heterogeneity of genetically defined subclones in acute myeloid leukemia. Cancer Cell. 2014;25(3):379–392.

54. Janjic A, Wange LE, Bagnoli JW, et al. Prime-seq, efficient and powerful bulk RNA sequencing. Genome Biol. 2022;23(1):88.

55. Roas M, Vick B, Kasper MA, et al. Targeting FLT3 with a new-generation antibody-drug conjugate in combination with kinase inhibitors for treatment of AML. Blood. 2023;141(9):1023–1035.

56. Mattar M, McCarthy CR, Kulick AR, Qeriqi B, Guzman S, de Stanchina E. Establishing and Maintaining an Extensive Library of Patient-Derived Xenograft Models. Front Oncol. 2018;8:19.

57. Sajjad H, Imtiaz S, Noor T, Siddiqui YH, Sajjad A, Zia M. Cancer models in preclinical research: A chronicle review of advancement in effective cancer research. Animal Model Exp Med. 2021;4(2):87–103.

58. Lai Y, Wei X, Lin S, Qin L, Cheng L, Li P. Current status and perspectives of patient-derived xenograft models in cancer research. J Hematol Oncol. 2017;10(1):106.

59. Boutzen H, Madani Tonekaboni SA, Chan-Seng-Yue M, et al. A primary hierarchically organized patient-derived model enables in depth interrogation of stemness driven by the coding and non-coding genome. Leukemia. 2022;36(11):2690–2704.

60. Velasco-Hernandez T, Trincado JL, Vinyoles M, et al. Integrative single-cell expression and functional studies unravels a sensitization to cytarabine-based chemotherapy through HIF pathway inhibition in AML leukemia stem cells. Hemasphere. 2024;8(2):e45.

61. Qin R, Liang Y, Zhou F. Advances in the application of patient-derived xenograft models in acute leukemia resistance. Cancer Drug Resist. 2025;8:23.

62. Tilsed CM, Fisher SA, Nowak AK, Lake RA, Lesterhuis WJ. Cancer chemotherapy: insights into cellular and tumor microenvironmental mechanisms of action. Front Oncol. 2022;12:960317.

63. Percie du Sert N, Hurst V, Ahluwalia A, et al. The ARRIVE guidelines 2.0: Updated guidelines for reporting animal research. PLoS Biol. 2020;18(7):e3000410.

64. Karp NA, Berdoy M, Gray K, et al. The Sex Inclusive Research Framework to address sex bias in preclinical research proposals. Nat Commun. 2025;16(1):3763.

65. Bonnet D. In vivo evaluation of leukemic stem cells through the xenotransplantation model. Curr Protoc Stem Cell Biol. 2008;Chapter 3:Unit 3 2.

66. R Core Team (2021). R: A language and environment for statistical computing. R Foundation for Statistical Computing, Vienna, Austria. https://www.R-project.org/.

