## Supplementary information for "A platform of robust patient-derived leukemia models covering subgroups for which no cell lines exist"

#### Supplemental Materials and Methods

##### Sample Cohorts

Primary adult patient samples of **cohort 1** (n=74) were collected between 2010 and 2019 in the Department of Internal Medicine III, Ludwig-Maximilians-Universität (LMU), Munich, Germany. Details on clinical parameters such as treatment regimens, survival, mutational status etc. were available for samples of this cohort.

Primary adult patient samples of **cohort 2** (n=28) were collected between 2012 and 2019 in the Department of Hematology, Oncology and Cancer Immunology, Charité, Berlin, Germany (n=9); Department of Medicine III, Technical University of Munich (TUM), Munich, Germany (n=6); Division of Hematology & Oncology, Department of Medicine, University of Pennsylvania, Philadelphia, USA (n=5); and Department of Internal Medicine I, University Hospital Carl Gustav Carus, University of Technology, Dresden, Germany (n=2).

Patient-derived xenograft (PDX) samples of **cohort 3** (n=22) were established at Inserm, Cancer Research Center of Toulouse, Toulouse, France (n=9)[1]; Department of Hematology and Oncology, Freiburg University Medical Center, Albert-Ludwigs-University of Freiburg, Breisgau, Germany (n=7); Department of General Pediatrics, Hematology and Oncology, University Children's Hospital, Tuebingen, Germany (n=5)[2]; Josep Carreras Leukemia Research Institute, Department of Biomedicine, School of Medicine, University of Barcelona, Barcelona, Spain (n=4)[3] or from the PRoXe consortium, Boston, USA[4] (n=2) and originated from adult or pediatric patients.

Primary pediatric patient samples of **cohort 4** (n=13) were collected between 2001 and 2014 in children's hospitals throughout Germany; PDX models thereof were established at Hannover Medical School and University Hospital Halle (Saale), Germany.

##### Mouse housing

All animal trials were performed in accordance with the current ethical standards of the official committee on animal experimentation (written approval by Regierung von Oberbayern,; 55.2-1-54-2531.6-10-10, 55.2-1-54-2531-95-10, 55.2-2532.Vet_03-16-56, 55.2-2532.Vet_02-16-7, 55.2-2532.Vet_02-20-159, 55.2-2532.Vet_03-21-9) and according to ARRIVE guidelines[5]. Mice were kept in animal rooms of the Laboratory Animal Breeding and Husbandry Unit of Helmholtz Munich under specified pathogen-free (SPF) conditions with a 12/12-hour light cycle. The animal rooms of the barriers were fully air-conditioned with a temperature of 20-24°C and 45-65% humidity according to Annex A of the European Convention 2007/526 EC. The maximum stocking density of the cages corresponds to Annex III of the 2010/63 EU. The cages were constantly filled with structural enrichment and the animals had unlimited access to food and water. During the experiment, mice were kept in individually ventilated cages (IVC). Hygiene monitoring was carried out at least quarterly in accordance with the FELASA recommendation: In the animal housing areas equipped with IVC systems, exhaust dust from the IVC ventilation units was tested for all FELASA-listed pathogens by PCR. Mice were examined daily, and health status was scored once a week (mice used for cell amplifications) or daily (mice receiving therapy).

**Isolation of primary AML blasts**

Human mononuclear cells were isolated from heparinized BM aspirates or PB by density gradient centrifugation (Ficoll-PaqueTM PLUS, GE Healthcare, Uppsala, Sweden). Cells were washed twice in phosphate buffered saline (PBS; Life Technologies, Darmstadt, Germany) and either viable frozen in 90% FCS (Biochrom AG, Berlin, Germany) / 10% DMSO (Sigma-Aldrich, St. Louis, MO, USA) or used for transplantation. Frozen cells were thawed according to published protocols[6].

**Monitoring engraftment of cells**

To monitor engraftment and growth of human cells in NSG mice, 50 µl of PB was repetitively collected by tail vein aspiration every other week starting from week four after cell injection. Depending on growth kinetics, sampling frequency was increased or decreased. BM was isolated after sacrifice of mice (see below). PB and BM were analyzed by flow cytometry after staining with antibodies against human CD45 (APC Mouse Anti-Human CD45 Clone HI30, BD Biosciences) and human CD33 (PE Mouse Anti-Human CD33 Clone WM53, BD Biosciences). Antibodies were added to 50 µL of PB or BM cell suspension, and incubated for 30 min. For PB, FACS lysing solution (BD Biosciences, Heidelberg, Germany) was added afterwards and cells were incubated for another 15 min at room temperature. Cells were washed twice with PBS. Flow cytometry was performed with a BD LSRFortessa (BD Biosciences) using FACS Diva software (BD Biosciences). Flow cytometry data was analyzed using FlowJo software (TreeStar Inc., Ashland, OR, USA). Percentage of hCD45+ hCD33+ cells were analyzed within the “lymphocyte gate” in FSC/SSC. Positive engraftment was defined as >1% hDC33+ cells in the BM within 20 weeks after transplantation.

**Experimental end-points**

The majority of mice used for PDX cell amplification were sacrificed at advanced leukemic disease (more than 50% leukemic cells within PB) or when mild clinical signs of illness appeared (e.g. rough fur, mild hunchback, or slightly slowed movement); few mice developed moderate or severe symptoms (e.g. reduced motility, lameness, increased breathing frequency). Despite close monitoring of mouse health status by daily examinations and flow cytometry of PB, individual mice were found dead, either due to high leukemia cell engraftment, or due to leukemia-unrelated reasons (n=21, 1% of mice included in the study). If 20-30 weeks after cell injection no human cells were detectable in PB and no clinical signs of illness arose, mice were sacrificed, and BM was analyzed by flow cytometry for human cell engraftment. Mice showing leukemia-unrelated illness or peculiarities were sacrificed and excluded from the studies. In experimental trials, specific endpoints were defined (i.e. defined bioluminescence signal). Mice which died in inhalation narcosis were excluded from further analyses. Mice showing therapy-related toxicity (i.e. loss of body weight beyond 15%) were sacrificed. Mice were sacrificed by exposure to $CO_2$ or cervical dislocation.

**Recovering PDX cells from mice and serial transplantation**

PDX cells were reisolated from bones and spleen if this organ was macroscopically enlarged. Femurs, tibiae, hips, sternum and spine were crushed using a mortar and pistil in 10 ml PBS and filtered through a 70 µm cell strainer (EASYstrainer, Greiner Bio-One, Frickenhausen, Germany). Cell pellet was incubated with red blood cell lysis buffer (Sigma Aldrich) for 10 minutes and washed three times in PBS. Spleens were smashed through a 70µm cell strainer and were ficollized. Cells were counted with Trypan-blue exclusion using a Neubauer counting chamber, analyzed by flow cytometry, and frozen in 90% FCS / 10% DMSO (until 2022) or in Bambanker (Nippon Genetics, Düren, Germany) in a density of 5E6-1E7 cells/ml and stored at -80°C or liquid nitrogen for long-term storage. Furthermore, 1E5 to 5E6 PDX cells were reinjected into next recipient mice for expansion or *in vivo* trials. Experimental mice were transplanted either with cells isolated freshly from donor mice, or with frozen/thawed cells from previous donor mice. Specimens were named with a specific nomenclature to be able to trace back donor cells and re-passaging steps.

**Authentication of passaged PDX cells**

Accuracy of model identity was regularly verified by repetitive finger printing using PCR of mitochondrial DNA[7]. Primary patient cells (if available) or early passage PDX cells were used as reference sequence.

**Lentiviral transduction and enrichment of genetically engineered PDX (GEPDX) cells**

Lentiviruses were produced with a third-generation lentiviral system as published[8]. In this study, AML PDX cells were genetically engineered to express enhanced firefly luciferase and mCherry as marker fluorochrome, Cre-ER$^{T2}$ and mCherry (Plasmids #104833 or #178185, Addgene, Watertown, Massachusetts, USA), or Cas9[9-11].

PDX cells were isolated from donor mice and resuspended in StemPro-34 medium (Thermo Fisher Scientific Waltham, MA, USA) supplemented with 1% L-Glutamin, 1% Penicillin/Streptomycin, 2% FCS (all Gibco, San Diego, CA, USA), 10 ng/ml hrFLT3L (R&D Systems, Minneapolis, MN, USA), 10 ng/ml hrSCF, 10 ng/ml hrTPO, and 10 ng/ml hrIL3 (all Peprotech, Rocky Hill, NJ, USA) as proposed recently[12] at a density of 1E7 cells per ml, and kept at 37°C, 5% $CO_2$. Cells were transduced overnight with lentiviral constructs in the presence of 8 µg/ml polybrene (Sigma-Aldrich). Transductions were performed at low MOI to guarantee single integrations. After 24 h to 48 h, cells were washed three times with PBS and injected into recipient mice for amplification. At advanced leukemic disease, mice were sacrificed and PDX cells were isolated from murine BM or spleen. GEPDX cells were enriched by sorting for mCherry or GFP positive cells on a FACSAria II (BD). Enriched GEPDX cells were re-injected into mice for amplification. All therapy trials were performed with GEPDX cells purified to levels beyond 90% mCherry+ cells.

**Bioluminescence *in vivo* imaging (BLI)**

Bioluminescence was measured using an IVIS Lumina 2 (Perkin Elmer) to repetitively visualize outgrowth of luciferase+ GEPDX cells in living mice. Mice were anesthetized by isoflurane inhalation. For cells expressing a recombinant codon-optimized form of firefly luciferase, 150 mg/kg D-Luciferin (BIOMOL GmbH, Hamburg, Germany) was injected into the tail vein. Images were taken immediately for 30 sec or up to 2 min using a field of view of 12.5 cm with binning 8, f/stop 1 and open filter setting. Images with more than 1000 counts were considered within the linear range; saturated images were repeated with reduced exposure time or reduced f-stop.

**Quantification of BLI pictures**

Quantification of light emission was performed using the Living Image software (Caliper Life Sciences, Mainz, Germany). A region of interest (ROI) covering the whole mouse was used and total flux was determined.

**Calculation of tumor growth rate by analyzing doubling time of BLI signals**

To calculate growth rate of tumor cells, an exponential trend line was adjusted within the logarithmic growth phase of PDX cells in individual mice (total flux between 1E7 and 1E10 Photons / second). BLI doubling time in days was calculated by dividing ln(2)/k, with k being the exponential growth coefficient $u(t)=-c*e^{k*t}$.

**Blood cell count**

Blood cell count was performed on an XN9000 automated analyzer (Sysmex, Norderstedt, Germany). Analyses were performed in a 1:7 dilution of EDTA-anticoagulated whole blood with Cellpack (Sysmex).

**Sequencing**

**DNA and RNA preparation**

gDNA was prepared using the QIAamp DNA Mini Kit (Qiagen), RNA was prepared using the Direct-zol RNA Miniprep (Zymo Research, Freiburg, Germany). Concentration of dsDNA was determined via Qubit dsDNA BR Assay Kit (Life Technologies); concentration of RNA was determined via NanoDrop, quality of RNA was determined via Agilent RNA 6000 Nano Kit (Agilent Technologies, Santa Clara, CA, USA).

**Targeted Sequencing of recurrently mutated genes**

68 genes recurrently mutated in myeloid malignancies were sequenced using a targeted amplicon-based enrichment assay (Haloplex, Agilent, Boeblingen, Germany) as previously described[13].

**Low coverage whole genome sequencing and data analysis**

DNA was prepared for whole genome sequencing using SureSelect WGS kit (Agilent Technologies, Santa Clara, CA, USA). Libraries were 100 bp paired-end sequenced using a NovaSeq 6000 sequencer (Illumina, San Diego, CA, USA). Median coverage was 1.60 fold.

**RNA sequencing and data analysis**

RNA was prepared using TruSeq Stranded Total RNA (Agilent Technologies). Libraries were 100 bp paired-end sequenced using a NovaSeq 6000 sequencer (Illumina, San Diego, CA, USA).

**Analysis of copy number variants (CNVs) from methylome**

CNVs from methylation data were inferred with the conumee2 package (v2.1.2) in R (v4.4.2). Raw IDAT files were processed using minfi (v1.52.1). Reference control sample of normal tissue was obtained from the minfiDataEPIC(v1.32.0). Processing followed documentation[14]. Probe-level annotations were generated with a minimum bin size of 500 kb and minimum probe count of 15.

**Analysis of CNVs from lcWGS**

CNVs from lcWGS were inferred using QDNAseq (v1.42.0/ R v4.4.2)[15] and ichorCNA (v0.3.2)[16], following each tools documentation. A bin size of 500 kb was used in both methods.

**CNV calling**

CNV calls from methylation data were considered high evidence if detected with a $\log_2$ ratio $\geq 0.5$ and confirmed by lcWGS data with a $\log_2$ ratio of $\geq 0.5$ based on one of the two tools plus $\geq 0.2$ based on the second tool (QDNAseq or ichorCNA). CNV calls were considered low evidence if not supported by lcWGS data.

In lcWGS we distinguish between high- and low-quality data. If the standard deviation of $log_2$ ratios in lcWGS data was ≥ 0.5 and in methylation data ≤ 0.25, samples were considered low quality. Data from samples AML-573, AML-346, AML-640, and AML-372 was considered low quality.

CNV calls from high quality lcWGS data were considered high evidence if detected with a $log_2$ ratio ≥ 0.5 by at least one of the two tools (QDNAseq or ichorCNA), a $log_2$ ratio of ≥ 0.2 in the second tool plus supported by methylation data with a $log_2$ ratio ≥ 0.2. Calls were considered low evidence if not supported by methylation data but detected using QDNAseq and ichorCNA, both with a $log_2$ ratio ≥ 0.5.

CNV calls from low quality lcWGS samples were only considered if detected with a $log_2$ ratio ≥ 0.5 in both tools (QDNAseq or ichorCNA) and supported by a $log_2$ ratio ≥ 0.4 in methylation data.

Calls are required to span at least two consecutive 500 kb bins.

**Analysis of fusion transcripts**

Fastq files were processed with the RIFTT pipeline (https://github.com/pkerbs/RIFTT), providing an evidence level in the range 1-6. Known fusions, that are listed in ChimerDB (here called predescribed), were considered if evidence level was five or six. Unknown fusions, without a corresponding entry in this database (here called undescribed), were considered if evidence level was six.

**Gene expression profiling, t-SNE plots**

Bulk RNA sequencing was performed as described in detail in[17]. Gene expression data were processed with the R packages *limma* (version 3.44.3) and *edgeR* (version 3.30.3). Samples with a library size of less than 500,000 were excluded from the analysis. Only genes that had more than one read per million in at least 10% of samples were included. The R package *Rtsne* (version 0.15) that implements the t-distributed stochastic neighbor embedding (t-SNE) method was used to cluster the data.

Gene set enrichment analysis was used to identify significantly enriched pathways in the Gene Ontology, KEGG and Hallmarks of Cancer databases. The normalized values of 18959 transcripts were used for the analysis, and the annotation file Human_Ensembl_Gene_ID_MSigDB.v2024.1.Hs.chip from Broad Institute was used to collapse Ensembl IDs to gene symbols. A q-value of <0.25 was considered significant.

**Methylome, t-SNE plots**

Methylation data was preprocessed with the R package *minfi* (version 1.50.0). Probes not mapped to a genomic region, probes measuring SNPs and non-CpG methylation, probes with SNPs and probes with missing values were removed from the analysis. M-values were used to cluster the data with the R package *Rtsne* (version 0.15). AML cell lines EoL-1, KASUMI-1, MOLM-13, OCI-AML-3, SKM-1, and THP-1 were included in the analysis.

**Clinical data sets**

Data generated with AML PDX models were compared to the following data sets: AML-CG-1999 (n=864)[18], AML-CG-2008 (n=274)[19], AMLHD98A (n=557)[20], AMLHD98B (n=160)[21], AML-SG07-04 (n=724)[22], AML-CG Registry (n=271)[23], primary AML samples of adult patients at relapsed disease of BEAT cohort (n=53)[24], primary AML samples of adult patients at initial diagnosis or relapse CN cohort (n=50)[25], AML cell lines generated from adult patients (n=54), eventually focusing on models with > 500 articles on Google Scholar as of March 2025 (n=19; AML-14, EoL-1, HEL, HL-60, KASUMI-3, KG-1, Me-1, ML-2, MOLM-13, MONO-MAC-6, NOMO-1, OCI-AML-2, OCI-AML-3, OCI-AML-5, SKM-1, SKNO-1, TF-1, U-937, UT-7)[26].

**Prediction analysis for microarrays (PAM) analysis**

Samples from the AML-CG-1999 and AML-CG-2008 study (training set) were used to build a classifier for genes mutated in at least 20% of PDX models. For each gene, the prediction analysis for microarrays (PAM)[27] was used to build a model that classifies samples into a non-mutated or a mutated group. The classifier was then tested on PDX models and human samples from the AML-CG Registry. To evaluate the performance, the true positive (TPR), true negative rate (TNR), positive predictive (PPV) and negative predictive value (NPV) were calculated.

The gene expression training set contained 218, and the methylation training set contained 209 samples[28]. The classifiers were tested on 271 gene expression and 224 methylation samples from the AML-CG Registry, and on 18 gene expression and 19 methylation PDX models.

#### Flow cytometry of surface antigens

Surface expression of CD14 (clone: RMO52), CD33 (D3HL60.251), CD34 (581), CD38 (LS198.4.3), CD64 (22), CD65 (88H7), CD117 (104D2D1), CD123 (7G3), CD135 (SF1.340), HLA-DR (Immu-357, all Beckman Coulter, Marseille, France), CD7 (M-T701), CD13 (L138), CD56 (NCAM16.2, all BD Biosciences, Milpitas, CA, USA), CD11b (ICRF44, Biolegend), CD15 (W6D3, both Biolegend, San Diego, CA, USA) was assessed on a Navios flow cytometer (Beckman Coulter, Krefeld, Germany).

#### Data Availability

Sequencing and methylome data are accessible through Gene Expression Omnibus: GSE307747 and GSE308997, respectively. Information on PDX models is shared via CancerModels.org [29].

For all other original data, please contact.

#### Illustrations

Schemes were created in BioRender. Klein, F. (2025) https://BioRender.com/ffj84eg

#### Supplemental Tables

**Supplemental Table 1: All samples**

| AHS-ID | Score | patient data | | | | | | | | | |
|---|---|---|---|---|---|---|---|---|---|---|---|
| | | origin, external ID | material | cohort | age | sex | Disease status | risk group ELN 2022 | FAB Subtyp | Cytogenetics | Fusions | panel seq (mutations) |
| 295 | 2 | KUM | primary | 1 | 20 | f | ID | Adverse | M5 | t(9;11)(p22;q23) | KMT2A::MLLT3 | BCOR, NRAS |
| 343 | 2 | KUM | primary | 1 | 80 | m | ID | Adverse | unknown | CN | unknown | FLT3-ITD, NPM1, IDH2, SRSF2 |
| 346 | 5 | Tübingen (P18R) | PDX | 3 | 1 | f | R | Adverse | unknown | unknown | unknown | cKIT |
| 356 | 6 | Tübingen (P17R) | PDX | 3 | 5 | m | R | Intermediate | unknown | unknown | unknown | unknown |
| 358 | 5 | Tübingen (P49S) | PDX | 3 | 9 | m | R | Adverse | unknown | unknown | unknown | unknown |
| 361 | 4 | KUM | primary | 1 | 40 | f | ID | Adverse | M4 | CN | unknown | DNMT3A, NPM1, FLT3-ITD, BCOR |
| 362 | 2 | KUM | primary | 1 | 71 | m | ID | Favorable | M4 | CN | unknown | DNMT3A, NPM1, CEBPA, IDH2 |
| 372 | 6 | KUM | primary | 1 | 41 | m | R | Adverse | M1 | complex including -17, -7, ETV-Del, ATM-Del | unknown | TP53, KRAS Monosomie 17, Monosmie 7, ETV-Deletion, ATM-Deletion, NRAS, KRAS, TP53 |
| 373 | 2 | KUM | primary | 1 | 79 | f | ID | Adverse | unknown | CN | unknown | DNMT3A, FLT3-ITD, moCEBPA, NRAS, BCOR, IDH1 |
| 388 | 5 | KUM | primary | 1 | 57 | m | ID, refractory | Adverse | M4 | t(6;11)(q27;q23) | KMT2A::AFDN | KRAS Q61H |
| 392 | 2 | KUM | primary | 1 | 65 | f | ID | Favorable | M2 | not done | unknown | not done; NPM1 |
| 393 | 6 | KUM | primary | 1 | 47 | f | R1 | Adverse | M4 | ins(10;11)(p12;q23q23) | KMT2A::MLLT10 | BCOR, KRAS |
| 403 | 2 | KUM | primary | 1 | 32 | f | ID | Adverse | M1 | complex, including ETV6-Del | unknown | PHF6, ASXL2 |
| 407 | 3 | KUM | primary | 1 | 58 | f | R1 | Adverse | unknown | t(4;8)(p1?5;q2?2),+12 | unknown | DNMT3A, SRSF2, NRAS, IDH1, NOTCH1 |
| 412 | 2 | KUM | primary | 1 | 65 | f | ID | Intermediate | M1 | CN | unknown | TET2, NPM1, FLT3-ITD |
| 415 | 3 | KUM | primary | 1 | 68 | f | R2 | Intermediate | unknown | CN | unknown | DNMT3A, FLT3-ITD, NPM1, IDH1 |
| 426 | 3 | KUM | primary | 1 | 55 | f | R1 | Adverse | M1 | ´+8 +13 | unknown | FLT3-ITD, FLT3-TKD, RUNX1, TET2 |
| 448 | 3 | KUM | primary | 1 | 61 | m | ID | Intermediate | M4 | CN | unknown | DNMT3A, NPM1, FLT3-ITD, FLT3-TKD |
| 452 | 2 | KUM | primary | 1 | 81 | f | ID | Intermediate | M1 | CN | unknown | TET2, NPM1, FLT3-ITD, MLL-PTD |
| 461 | 2 | KUM | primary | 1 | 65 | f | ID | Intermediate | M1 | CN | unknown | NPM1, FLT3-ITD |
| 479 | 3 | TUM | primary | 2 | 31 | f | refractory | unknown | unknown | unknown | unknown | not done |
| 485 | 3 | Dresden | primary | 2 | 46 | f | R | unknown | unknown | unknown | unknown | not done |
| 489 | 2 | KUM | primary | 1 | 31 | f | ID | Favorable | M2 | CN | unknown | NPM1, IDH2, TET2, TET2-AS1 |
| 491 | 6 | KUM | primary | 1 | 53 | f | R1 of 661 | Adverse | M2 | del(7)(q2?1) / s1,t(2;11)(p11;q2?3) | unknown | DNMT3A, BCOR, NRAS, KRAS, ETV6, PTPN11, RUNX1 |
| 506 | 3 | KUM | primary | 1 | 29 | m | R1 | Adverse | M5 | complex, including +8, t(9;11)(p22;q23) | KMT2A::MLLT3 | ASXL1 |
| 530 | 3 | Berlin | primary | 2 | 54 | m | R | unknown | unknown | CN at ID | unknown | NPM1, FLT3-ITD |
| 538 | 5 | Berlin | primary | 2 | 68 | f | R | Adverse | unknown | CN at ID | unknown | DNMT3A, IDH1 |
| 573 | 5 | KUM | primary | 1 | 63 | f | R1, sAML | Intermediate | M1 | t(5;11) at ID | unknown | DNMT3A, IDH2, FLT3-ITD, WT1, MLL-PTD |
| 579 | 5 | KUM | primary | 1 | 50 | m | R1 | Intermediate | M5 | CN | unknown | DNMT3A, NPM1, FLT3-ITD, IDH1 |
| 596 | 3 | TUM | primary | 2 | 55 | m | ID | unknown | unknown | CN | unknown | NPM1 mut, FLT3 wt |
| 601 | 2 | Berlin | primary | 2 | 69 | m | R | unknown | unknown | inv(16)(p13q22) at ID | unknown | not done |
| 602 | 5 | Berlin | primary | 2 | 40 | f | R1 | Intermediate | unknown | unknown | unknown | TET2, NPM1, FLT3-ITD, CEBPA, JAK3 |
| 613 | 2 | Berlin | primary | 2 | 66 | f | R | unknown | unknown | complex at ID | unknown | not done |
| 625 | 2 | KUM | primary | 1 | 36 | f | R1 | Adverse | M4 | CN | unknown | FLT3-ITD, DNMT3A, NPM1, BCOR, IDH2 |
| 640 | 6 | Berlin | primary | 2 | 79 | m | R | Intermediate | unknown | t(11;15) | unknown | NPM1, FLT3-ITD, IDH1 |
| 661 | 5 | KUM | primary | 1 | 54 | f | R2 of 491 | Adverse | M2 | del7q (7q21.13 q36.3); del6p (6p25.3 p21.1) | unknown | DNMT3A, BCOR, NRAS, ETV6, PTPN11, RUNX1, EZH2 |
| 663 | 6 | Tübingen (P84D/P21) | PDX | 3 | 13 | f | R | Intermediate | unknown | unknown | unknown | FLT3-ITD, WT1 |
| 664 | 5 | Tübingen (P93A) | PDX | 3 | 6 | f | R | Intermediate | unknown | unknown | unknown | WT1 |
| 669 | 5 | KUM | primary | 1 | 48 | f | R2 | Adverse | M5 | t(9;11)(p22;q23) | KMT2A::MLLT3 | none detected |
| 760 | 2 | Harlaching | primary | 2 | adult | unknown | unknown | unknown | unknown | not done | unknown | not done |
| 788 | 3 | KUM | primary | 1 | 55 | f | ID of 979 | Intermediate | M4 | CN | unknown | DNMT3A, NPM1, FLT3-ITD |
| 810 | 3 | KUM | primary | 1 | 38 | f | ID of 842 | Intermediate | M2 | ´+8 | unknown | DNMT3A, NPM1, FLT3-ITD, DCLK1 |
| 842 | 4 | KUM | primary | 1 | 39 | f | R1 of 810 | Adverse | M4 | complex, including +8 | unknown | NPM1, FLT3-ITD, DCLK1, DNMT3A, |
| 896 | 4 | TUM | primary | 2 | 52 | f | ID | Intermediate | unknown | CN | unknown | DNMT3A, IDH2, NPM1, FLT3-ITD, ASXL2 |
| 954 | 2 | Freiburg (197) | PDX | 3 | adult | unknown | unknown | unknown | unknown | not done | unknown | not done |
| 955 | 5 | Freiburg (342) | PDX | 3 | adult | f | unknown | Favorable | unknown | CN | unknown | DNMT3A, NPM1, WT1 |
| 957 | 2 | Toulouse (IM06b) | PDX | 3 | adult | unknown | unknown | unknown | unknown | not done | unknown | not done |
| 958 | 3 | Toulouse (IM10b) | PDX | 3 | adult | unknown | unknown | unknown | unknown | not done | unknown | not done |
| 959 | 3 | Toulouse (IM13b) | PDX | 3 | adult | unknown | unknown | unknown | unknown | not done | unknown | NPM1, FLT3-ITD, DNMT3A, TET2 (2x) |
| 960 | 3 | Toulouse (IM17) | PDX | 3 | adult | unknown | unknown | unknown | unknown | not done | unknown | not done |
| 961 | 2 | Toulouse (IM18) | PDX | 3 | adult | unknown | unknown | unknown | unknown | not done | unknown | not done |
| 962 | 2 | Toulouse (IM16) | PDX | 3 | adult | unknown | unknown | unknown | unknown | not done | unknown | not done |
| 979 | 6 | KUM | primary | 1 | 56 | f | R2 of 788 | Intermediate | M4 | CN at ID | unknown | DNMT3A, NPM1, FLT3-ITD, IDH2, WT1 |
| 980 | 3 | KUM | primary | 1 | 76 | m | ID of 981 | Adverse | M1 | CN | unknown | DNMT3A, TET2, NPM1, FLT3-TKD, ASXL1, FLT3-ITD |
| 981 | 5 | KUM | primary | 1 | 76 | m | R1 of 980 | Adverse | M1 | CN at ID | unknown | DNMT3A, TET2, NPM1, FLT3-TKD, ASXL1, FLT3-ITD |
| 1176 | 2 | ProXe (DFAM- | PDX | 3 | 67 | m | R | Intermediate | M4 | ´+8,+1,del(1)(p22p36) | unknown | none detected |
| 1177 | 5 | ProXe (DFAM- | PDX | 3 | 58 | m | ID (response unclear) | Intermediate | M4 | CN | unknown | NPM1, FLT3-ITD |
| 1183 | 2 | BAR (14011) | PDX | 3 | 1,5 | m | ID | unknown | M5 | Inv16 | CBFb::MYH11 | not done |
| 1184 | 2 | BAR (14030) | PDX | 3 | 18 | m | ID | unknown | unknown | CN | unknown | not done |
| 1186 | 5 | BAR (14048) | PDX | 3 | 63 | f | ID (response unclear) | Intermediate | M4 | t(9;11)(p22;q23) | KMT2A::MLLT3 | not done |
| 1191 | 2 | BAR (14186) | PDX | 3 | <1 | m | ID | unknown | M5b | t(11;17)(q23;q12) | KMT2A::MLLT6 | not done |
| 1221 | 4 | Toulouse (IM28) | PDX | 3 | >70 | f | ID | unknown | unknown | not done | unknown | FLT3-TKD, FLT3-ITD, WT1, TET2 (2x) |
| 1222 | 5 | Toulouse (IM40) | PDX | 3 | >30 | m | R | Intermediate | unknown | not done | unknown | RAD21, DNMT3A, NPM1, FLT3-ITD |
| 1223 | 2 | Toulouse (IM48) | PDX | 3 | >40 | m | ID | unknown | unknown | not done | unknown | not done |
| 1224 | 3 | Philadelphia | primary | 2 | adult | unknown | ID | Adverse | M4 | complex | unknown | not done |
| 1225 | 4 | Philadelphia | primary | 2 | adult | unknown | ID | Intermediate | unknown | t(9;11)(p22;q23),?add(10)(q?24 | KMT2A::MLLT3 | ASXL1, U2AF1 |
| 1226 | 5 | Philadelphia | primary | 2 | adult | f | ID, refractory | Intermediate | M4 | not done | unknown | TET2 (2x), WT1 (2x), KIT |
| 1227 | 2 | Philadelphia | primary | 2 | adult | unknown | R | unknown | unknown | not done | unknown | FLT3-TKD, STAG2 |
| 1228 | 4 | Philadelphia | primary | 2 | adult | unknown | tAML | unknown | unknown | 45,XY,add(3)(q13.3),-7[2]/45,idem,ins(8;7)(p23;q11.2q32)[19] | unknown | not done |
| R139 | ≥4 | FFM | PDX | 4 | 5 | f | ID | unknown | unknown | ´+19 | unknown | unknown |
| H615 | ≥4 | FFM | PDX | 4 | 16 | m | ID | unknown | unknown | t(6;11)(q27;q23) | unknown | NRAS |
| H826 | ≥4 | FFM | PDX | 4 | 7 | f | ID | unknown | unknown | t(9;11)(p22;q23) | unknown | unknown |
| H958 | ≥4 | FFM | PDX | 4 | 5 | f | ID | unknown | unknown | t(10;11)(p12;q14) | unknown | unknown |
| H1454 | ≥4 | FFM | PDX | 4 | 1 | f | ID | unknown | unknown | t(9;11)(p22;q23) | unknown | unknown |
| H1555 | 3 | FFM | PDX | 4 | 13 | m | unknown | unknown | unknown | t(9;11)(p22;q23) | unknown | unknown |
| H2788 | ≥4 | FFM | PDX | 4 | 1 | m | ID | unknown | unknown | t(9;11)(p22;q23) | unknown | unknown |

Table S1A. Score 2-6 Samples. Patient data

| AHS-ID | Score | PDX data | | | | | |
|---|---|---|---|---|---|---|---|
| | | Cytogenetics | Transcriptome (predescribed fusion genes) | Transcriptome (Undescribed fusion genes) | panel seq (mutations) | lcWGS + Methylome (CNAs) | WHO 2022 group |
| **295** | 2 | *not done* | *not done* | *not done* | *not done* | *not done* | |
| **343** | 2 | *not done* | *not done* | *not done* | *not done* | *not done* | |
| **346** | **5** | interstitial 5q Del / interstitial 13q Del | CBFA2T3::GLIS2 (inv16) | AFF1::LINC00863 | cKIT | del(5q), del(13q), amp(13q) -> follows after the deletion | AML, myelodysplasia-related |
| **356** | **6** | *not done* | KMT2A::MLLT3 | none detected | U2AF1 S34Y (.49), KRAS A146T (.51) | del(1p), amp(2q), amp(3q), amp(6p), +8, del(10p),amp(11q), amp(19q) | AML with KMT2A rearrangement |
| **358** | **5** | 46,XY,del(7)(p22p11),der(20)t(17;20)(q22;q13) | *not done* | *not done* | FLT3 N839E (.46) | del(7p), amp(17q) | AML, defined by differentiation |
| **361** | 4 | *not done* | *not done* | *not done* | DNMT3A R882H (.5), NPM1 W288Cfs* (.5), FLT3-ITD (1), BCOR D551N (.5) | *not done* | |
| **362** | 2 | *not done* | *not done* | *not done* | *not done* | *not done* | |
| **372** | **6** | complex, including 5q13 Del/ETV6-Del/-7/-17 | none detected | PPFIA1::GRIK4, FAM168A::DLG2, ZBTB44-DT::ZBTB44, PEX1::TMUB2, FMN1::SIRT1, CTD-2537O9.1::IFT46, SUFU::UQCC1, SGMS1::FMN1, JMJD1C::FMN1, CPEB3::RALGAPA2, AC007566.10::BECN1, FN3KRP::SUPT6H | TP53 R248Q (.99), KRAS G12V (.33) | del(5q), -7, amp(8q), del(10q), del(11q), del(12p), del(13q), del(14q) del(15p), del(16q), del(17p), del(20p) | AML, myelodysplasia-related |
| **373** | 2 | *not done* | *not done* | *not done* | DNMT3A R882H (.5), DNMT3A S663P (.5), FLT3-ITD (.3), CEBPA P23fs (.5), BCOR R810* (.5) | *not done* | |
| **388** | **5** | t(6;11)(q27;q23) | KMT2A::AFDN | PVT1::RP11-367L7.1, RP11-770J1.8::AFDN | KRAS Q61H (.38) | none detected | AML with KMT2A rearrangement |
| **392** | 2 | *not done* | *not done* | *not done* | *not done* | *not done* | |
| **393** | **6** | ins(10;11)(p12;q23q23) | KMT2A::MLLT10 | MLLT10::CUL5 | KRAS G12A (.46), BCOR P1012Lfs*8 (.44) | amp(1p), amp(2q), amp(3q), amp(5q), amp(6q), del(7p), amp(8q), del(10q), amp(13q), amp(14q), del(14q), amp(17q), amp(21q) | AML with KMT2A rearrangement |
| **403** | 2 | *not done* | *not done* | *not done* | *not done* | *not done* | |
| **407** | 3 | *not done* | *not done* | *not done* | DNMT3A R882C (.99), NRAS Q61H (.4), SRSF2 P95H (.5), NOTCH1 S1690L (.4) | *not done* | |
| **412** | 2 | *not done* | *not done* | *not done* | TET2 A1097Vfs* (.5), NPM1 W288Cfs* (.5), FLT3-ITD (.1) | *not done* | |
| **415** | 3 | *not done* | *not done* | *not done* | DNMT3A V527I (.5), DNMT3A S714C (.5), NPM1 W288Cfs* (.5), FLT3-ITD (.99), IDH1 R132H (.5) | del(15q) | |
| **426** | 3 | *not done* | *not done* | *not done* | *not done* | *not done* | |
| **448** | 3 | *not done* | *not done* | *not done* | *not done* | *not done* | |
| **452** | 2 | *not done* | *not done* | *not done* | TET2 S1582Lfs* (.5), NPM1 W288Cfs* (.4), FLT3-ITD (.4) | *not done* | |
| **461** | 2 | *not done* | *not done* | *not done* | *not done* | *not done* | |
| **479** | 3 | *not done* | *not done* | *not done* | *not done* | *not done* | |
| **485** | 3 | *not done* | *not done* | *not done* | *not done* | *not done* | |
| **489** | 2 | *not done* | *not done* | *not done* | *not done* | *not done* | |
| **491** | **6** | del7q (7q21.13 q36.3) | none detected | none detected | DNMT3A R882S (.47), BCOR P683Qfs*32 (.52), NRAS Q61K (.05), KRAS G12A (.43), ETV6 P214L (.43), RUNX1 N136K (.55), PTPN11 D61H (.46), JAK1 V658F (.005) | amp(2q), amp(4q), amp(6q), del(7q), del(10q) | AML, myelodysplasia-related |
| **506** | 3 | *not done* | *not done* | *not done* | *not done* | *not done* | |
| **530** | 3 | *not done* | *not done* | *not done* | *not done* | *not done* | |
| **538** | **5** | *not done* | BCR::ABL1; KMT2A-PTD | none detected | DNMT3A R836Efs*19 (.46), DNMT3A G381Pfs*8 (.43), IDH1 R132C (.50) | amp(2q), amp(4q), amp(5q), amp(13q), amp(18q) | AML with BCR::ABL1 fusion |
| **573** | **5** | CN | none detected | none detected | DNMT3A S663L (.39), FLT3-ITD (.34), WT1 H257* (.47), WT1 R157PRG (.47), IDH2 R140Q (.49) | none detected | AML, myelodysplasia-related |
| **579** | **5** | CN | none detected | none detected | DNMT3A R882C (.48), DNMT3A F868L (.50), NPM1 W288Cfs*12 (.51), FLT3-ITD (.99), IDH1 R132H (.52) | del(7q), del(11p) | AML with NPM1 mutation |
| **596** | 3 | *not done* | *not done* | *not done* | *not done* | *not done* | |
| **601** | 2 | *not done* | *not done* | *not done* | *not done* | *not done* | |
| **602** | **5** | *not done* | t(4;12) involving ETV6 | CYRIB::GRID1 | NPM1 W288Cfs*12 (.45), FLT3-ITD (.29), TET2 N281* (.40), TET2 S1369* (.14), CEBPA D262Rfs*59 (.51), JAK3 V722I (.55) | del(3q), del(11p) | AML with NPM1 mutation |
| **613** | 2 | *not done* | *not done* | *not done* | *not done* | *not done* | |
| **625** | 2 | *not done* | *not done* | *not done* | *not done* | *not done* | |
| **640** | **6** | t(11;15)(p1?1;q?22) | none detected | WT1::LDLRAD3, YWHAE::UBE2Q2 | NPM1 W288Cfs*12 (.47), FLT3-ITD (.52), IDH1 R132H (.36) | none detected | AML with NPM1 mutation |
| **661** | **5** | del7q (7q21.13 q36.3) | none detected | none detected | DNMT3A R882S (.48), BCOR P683Qfs*32 (.48), ETV6 P214L (.49), RUNX1 N136K (.52), PTPN11 D61H (.51), EZH2 A692G | del(7q) | AML, myelodysplasia-related |
| **663** | **6** | CN | none detected | none detected | FLT3-ITD (.48), WT1 A170Rfs* (.47), WT1 C241Y (.55) | none detected *(no lcWGS)* | AML, defined by differentiation |
| **664** | **5** | *not done* | KMT2A::MLLT3 | none detected | WT1 D469V (.62) | del(2q), +21 | AML with KMT2A rearrangement |
| **669** | **5** | *not done* | KMT2A::MLLT3 | CACNA1C-AS1::ITFG2-AS1 | KRAS G13D (.03) | del(12q) | AML with KMT2A rearrangement |
| **760** | 2 | *not done* | *not done* | *not done* | *not done* | *not done* | |
| **788** | 3 | *not done* | *not done* | *not done* | DNMT3A R882H (.45), NPM1 W288Cfs* (.5), FLT3-ITD (.05) | del(2q) | |
| **810** | 3 | *not done* | *not done* | *not done* | DNMT3A R882C (.5), NPM1 W288Cfs* (.3), FLT3-ITD (.45) | *not done* | |
| **842** | 4 | *not done* | *not done* | *not done* | DNMT3A R882C (.44), NPM1 W288Cfs* (.5), FLT3-ITD (1.0 / .5) | -8 *(no lcWGS)* | |
| **896** | 4 | *not done* | *not done* | *not done* | DNMT3A R882H (.45), NPM1 W288Cfs* (.5), FLT3-ITD (.99), IDH2 R140Q (.5), ASXL2 G1057R (.5) | none detected *(no lcWGS)* | |
| **954** | 2 | *not done* | *not done* | *not done* | *not done* | *not done* | |
| **955** | **5** | CN | none detected | PHF3::CDKAL1 | DNMT3A S714C (.35), NPM1 W288Cfs*12 (.63), WT1 P381Q (.54) | del(13q) | AML with NPM1 mutation |
| **957** | 2 | *not done* | *not done* | *not done* | *not done* | *not done* | |
| **958** | 3 | *not done* | *not done* | *not done* | *not done* | *not done* | |
| **959** | 3 | *not done* | *not done* | *not done* | DNMT3A R882H (.5), NPM1 W288Cfs* (.5), FLT3-ITD (.5), TET2 Y1294C (.5) | del(2q) *(no lcWGS)* | |
| **960** | 3 | *not done* | *not done* | *not done* | *not done* | *not done* | |
| **961** | 2 | *not done* | *not done* | *not done* | *not done* | *not done* | |
| **962** | 2 | *not done* | *not done* | *not done* | *not done* | *not done* | |
| **979** | **6** | complex; 46,XX,t(11;16)(p12;p13),t(11;22)(p13;q13)[2]/46,idem,t(4;20)(q28;q13)[15]/46,idem,inv(1)(p36q12),t(4;20)(q28;q13)[3] | none detected | none detected | DNMT3A R882H (.41), NPM1 W288Cfs*12 (.49), IDH2 R140Q (.50), FLT3-ITD (?) | none detected *(no lcWGS)* | AML with NPM1 mutation |
| **980** | 3 | *not done* | *not done* | *not done* | DNMT3A Y448* (.36), NPM1 W288Cfs*12 (.49), FLT3-ITD (.42), ASXL1 P783A (.49), FLT3 D835H (.49), TET2 R1404* (1.0) | none detected *(no lcWGS)* | AML with NPM1 mutation |
| **981** | **5** | 46,XY,t(16;17)(q24;p12)[6] | none detected | GSE1::ELAC2 | DNMT3A Y448* (.41), NPM1 W288Cfs*12 (.52), FLT3-ITD (.96), ASXL1 P783A (.50), FLT3 D835H (.99), TET2 R1404* (.99), WT1 T165Pfs*9 (.39) | none detected | AML with NPM1 mutation |
| **1176** | 2 | *not done* | *not done* | *not done* | *not done* | *not done* | |
| **1177** | **5** | del(9)(q21q34) | none detected | none detected | DNMT3A R882H (.48), NPM1 W288Cfs*12 (.56), FLT3-ITD (pos.) | del(9q) *(no methylome)* | AML with NPM1 mutation |
| **1183** | 2 | *not done* | *not done* | *not done* | *not done* | *not done* | |
| **1184** | 2 | *not done* | *not done* | *not done* | *not done* | *not done* | |
| **1186** | **5** | *not done* | KMT2A::MLLT3 | none detected | KRAS G13dup (.44) | none detected *(no methylome)* | AML with KMT2A rearrangement |
| **1191** | 2 | *not done* | *not done* | *not done* | *not done* | *not done* | |
| **1221** | 4 | *not done* | none detected | none detected | TET2 L957Yfs*50 (.5), TET2 Y1337* (.3), WT1 R458* (1.0), FLT3-ITD (pos.), FLT3 D835H (.2) | none detected *(no methylome)* | |
| **1222** | **5** | *not done* | none detected | MAPKAPK5::ACAD10 | DNMT3A R882H (.46), NPM1 W288Cfs*12 (.47), FLT3-ITD (pos.), RAD21 R146Gfs*21 (.46) | del(11p) *(no methylome)* | AML with NPM1 mutation |
| **1223** | 2 | *not done* | *not done* | *not done* | *not done* | *not done* | |
| **1224** | 3 | *not done* | *not done* | *not done* | *not done* | *not done* | |
| **1225** | **4** | *not done* | KMT2A::MLLT3 | RP11-770J1.8::MLLT3 | ASXL1 G646Wfs*12 (.27), U2AF1 Q157P (.5) | none detected *(no methylome)* | AML with KMT2A rearrangement |
| **1226** | **5** | 47,XX,+19[5]/47,XX,del(9)(p24p13),+19[15] | none detected | TRIP12::SLC16A14 | TET2 N439Tfs*8 (.48), TET2 C1289Y (.42), KIT D816V (.45), WT1 A113Rfs*86 (.44), WT1 Y295* (.45) | none detected *(no methylome)* | AML, defined by differentiation |
| **1227** | 2 | *not done* | *not done* | *not done* | *not done* | del(11p) *(no methylome)* | |
| **1228** | 4 | *not done* | none detected | none detected | PTPN11 Q79R (.42), CCND2 T282_D283insE (.2) | -7 *(no methylome)* | |

Table S1B. Score 2-6 Samples. PDX data

| AHS-ID | Score | mouse data | | | | | | | | | | | | |
|---|---|---|---|---|---|---|---|---|---|---|---|---|---|---|
| | | last P | Mean passaging time [d] (min-max) | robustness of engraftment | % hCD33+ in PB (highest) | mice with enlarged spleen (>2e7) | preferred mouse gender | # PDX cells from spleen *1e6 (median) | % hCD33+ in spleen (median) | # PDX cells from BM *1e6 (median) | % hCD33+ in BM (median) | organ for Re-Tx | cell number, state | problems in vivo |
| **295** | 2 | 1 | 98 - 147 | 80% | *not done* | none | | 30 | 50% | 60 | 40% | BM | | caused extramedullary tumors |
| **343** | 2 | 1 | 150 | 25% | *not done* | none | | *not done* | *not done* | 40 | 25% | BM | | no re-engraftment |
| **346** | **5** | 16 | 47 (34 - 126) | 89,55% | 50% | **69%** | female | 161 | 94% | 32 | 87% | BM or Spl | >0,5Mio cells | engraftment <90% |
| **356** | **6** | 18 | 42 (26 - 82) | 97,37% | 60% | **69%** | male or female | 43 | 88% | 34 | 83% | BM or Spl | fresh if possible | fine |
| **358** | **5** | 10 | 71 (42 - 91) | 93,10% | 80% | **54%** | male or female | 31 | 52% | 18 | 86% | BM or Spl | | >70d; low transduction efficiency |
| **361** | 4 | 6 | 83 (43 - 158) | 75% | 85% | none | | *not done* | *not done* | 50 | 90% | BM or Spl | | no transduction possible |
| **362** | 2 | 1 | 60 - 150 | 10% | *not done* | none | | *not done* | *not done* | 83 | 18% | BM | | no re-engraftment |
| **372** | **6** | 10 | 66 (34 - 118) | 98,54% | 80% | **16%** | male or female | 25 | 83% | 43 | 95% | BM or Spl | >0,5Mio cells; fresh if possible | fine |
| **373** | 2 | 2 | 87 (66 - 95) | 53% | 60% | none | | *not done* | *not done* | 35 | 95% | BM | | no re-engraftment after P1 |
| **388** | **5** | 14 | 42 (24 - 89) | 87,50% | 90% | **56%** | female | 39 | 89% | 41 | 82% | BM or Spl | >0,5Mio cells | engraftment <90% |
| **392** | 2 | 1 | 150 | 25% | *not done* | none | | *not done* | *not done* | *not done* | 1% | BM | | no re-engraftment |
| **393** | **6** | 15 | 42 (25 - 99) | 91,76% | 90% | **69%** | male or female | 170 | 85% | 35 | 80% | BM or Spl | >0,5Mio cells | mice might die from one day to the other |
| **403** | 2 | 1 | 150 | 25% | *not done* | none | | *not done* | *not done* | 20 | 2% | BM | | no re-engraftment |
| **407** | 3 | 7 | 120 (85 - 160) | 75% | 60% | none | | *not done* | *not done* | 44,5 | 73% | BM | | slow, no transduction possible |
| **412** | 2 | 5 | 113 (71 - 160) | 50% | 87% | 20% | | 80 | 94% | 43 | 76% | BM or Spl | | no re-engraftment after thawing, slow, no transduction possible |
| **415** | 3 | 10 | 124 (70 - 160) | 60% | 75% | none | | *not done* | *not done* | 48,5 | 85% | BM | | slow, inhomogeneous growth |
| **426** | 3 | 2 | 135 (90 - 150) | 38% | 40% | none | | *not done* | *not done* | 36 | 31% | BM | | slow, low and inconsistent engraftment |
| **448** | 3 | 1 | 160 (140-180) | 67% | 10% | none | | *not done* | *not done* | 50 | 83% | BM | | slow |
| **452** | 2 | 3 | 105 (55-150) | 43% | 94% | 43% | | 65 | 95% | 47 | 69% | BM | | T cell engraftment might influence blast engraftment |
| **461** | 2 | 1 | 150 | 50% | *not done* | none | | *not done* | *not done* | 20 | 2% | BM | | no re-engraftment |
| **479** | 3 | 2 | 150 | 44% | 30% | none | | *not done* | *not done* | 76 | 55% | BM | | slow, low and inconsistent engraftment |
| **485** | 3 | 2 | 130 | 63% | 30% | none | | *not done* | *not done* | 72 | 91% | BM | | slow, inconsistent engraftment |
| **489** | 2 | 1 | 150 | 50% | *not done* | none | | *not done* | *not done* | 30 | 9,5% | BM | | no re-engraftment |
| **491** | **6** | 12 | 57 (32 - 161) | 96,84% | 80% | none | male or female | *not done* | *not done* | 50 | 86% | BM | | fine |
| **506** | 3 | 2 | 130 (100 - 150) | 100% | 5% | none | | *not done* | *not done* | 17 | 25% | BM | | slow, very low engraftment |
| **530** | 3 | 1 | 170 | 100% | 5% | none | | *not done* | *not done* | 59 | 42% | BM | | slow, low engraftment |
| **538** | **5** | 7 | 72 (55 - 132) | 92,86% | 50% | **7%** | female | 35 | *not done* | 76 | 86% | BM | >0,5Mio cells; fresh if possible | >70d; low transduction efficiency |
| **573** | **5** | 10 | 76 (53 - 152) | 92,86% | 90% | none | female | *not done* | *not done* | 60 | 95% | BM or Spl | >0,5Mio cells | >70d; low transduction efficiency |
| **579** | **5** | 7 | 75 (40 - 188) | 91,96% | 80% | none | female | *not done* | *not done* | 56 | 90% | BM or Spl | >0,5Mio cells; fresh if possible | >70d; low transduction efficiency |
| **596** | 3 | 1 | 120 | 67% | 70% | none | | *not done* | *not done* | 50 | 53% | BM | | slow, low BM engraftment; decreased in P1 |
| **601** | 2 | 1 | 150 | 25% | *not done* | none | | *not done* | *not done* | *not done* | 15% | BM | | no re-engraftment |
| **602** | **5** | 11 | 61 (41 - 96) | 87,50% | 60% | **16%** | female | 45 | 71% | 35 | 92% | BM or Spl | fresh if possible | engraftment <90% |
| **613** | 2 | 1 | 150 | 50% | *not done* | none | | *not done* | *not done* | 84 | 83% | BM | | no re-engraftment |
| **625** | 2 | 1 | 150 | 50% | *not done* | none | | *not done* | *not done* | 40 | 27% | BM | | no re-engraftment |
| **640** | **6** | 13 | 49 (38 - 133) | 96,10% | 70% | **23%** | female | 37 | 85% | 39 | 90% | BM or Spl | >0,5Mio cells; fresh if possible | fine |
| **661** | **5** | 11 | 50 (35 - 96) | 100% | 80% | none | male or female | *not done* | *not done* | 41 | 84% | BM or Spl | | low transduction efficiency |
| **663** | **6** | 8 | 41 (30 - 71) | 100% | 80% | **59%** | male or female | 75 | 59% | 61 | 92% | BM | fresh if possible | fine |
| **664** | **5** | 4 | 61 (39 - 89) | 84,21% | 70% | **12%** | male or female | 19 | *not done* | 62 | 93% | BM | >0,5Mio cells | engraftment <90% |
| **669** | **5** | 8 | 105 (36 - 158) | 72% | 60% | **22%** | female | 71 | 75% | 61 | 89% | BM | fresh if possible | >70d; engraftment <90% |
| **760** | 2 | 1 | 150 | 75% | *not done* | none | | *not done* | *not done* | 28 | 74% | BM | | slow, low engraftment; decreased in P1 |
| **788** | 3 | 1 | 175 | 57% | 30% | none | | *not done* | *not done* | 49 | 67% | BM | | slow |
| **810** | 3 | 1 | 150 (130 - 170) | 50% | 20% | none | | *not done* | *not done* | 53 | 81% | BM | | slow, inconsistent engraftment |
| **842** | 4 | 5 | 110 (70 - 160) | 72% | 60% | none | | *not done* | *not done* | 49 | 79% | BM | | rather slow, even slower if transduced |
| **896** | 4 | 4 | 120 (80 - 170) | 87% | 40% | none | | *not done* | *not done* | 40 | 84% | BM | | rather slow, transduced even slower/inconsistent; seem to lose transgene over time |
| **954** | 2 | 1 | *no re-engraftment* | *not done* | 0% | none | | *not done* | *not done* | *not done* | 0% | | | |
| **955** | **5** | 7 | 81 (64 - 97) | 100% | 80% | **85%** | male or female | 52 | 99% | 38 | 93% | BM or Spl | >0,5Mio cells | >70d |
| **957** | 2 | 1 | *no re-engraftment* | *not done* | 0% | none | | *not done* | *not done* | *not done* | 0% | | | |
| **958** | 3 | 3 | 190 | 75% | 5% | none | | *not done* | *not done* | 46 | 33% | BM | | slow, low engraftment |
| **959** | 3 | 6 | 130 (85 - 200) | 82% | 30% | none | | *not done* | *not done* | 55 | 72% | BM | | slow |
| **960** | 3 | 3 | 150 | 83% | 50% | none | | *not done* | *not done* | 41 | 90% | BM | | slow, low engraftment |
| **961** | 2 | 1 | *no re-engraftment* | *not done* | 0% | none | | *not done* | *not done* | *not done* | 0% | | | |
| **962** | 2 | 1 | *no re-engraftment* | *not done* | 0% | none | | *not done* | *not done* | *not done* | 0% | | | |
| **979** | **6** | 4 | 69 (49 - 86) | 100% | 70% | none | male or female | *not done* | *not done* | 57 | 93% | BM | fresh if possible | >70d |
| **980** | 3 | 3 | 130 (78 - 190) | 56% | 30% | none | | *not done* | *not done* | 46 | 76% | BM | | slow, inconsistent engraftment |
| **981** | **5** | 5 | 88 (54 - 133) | 96% | 70% | **79%** | male or female | 43 | 90% | 46 | 90% | BM or Spl | | >70d; low transduction efficiency |
| **1176** | 2 | 5 | 50 (47 - 73) | 71% | 12% | none | | *not done* | *not done* | 24 | 25% | BM | | induces paralysis in mice, inconsistent bm engraftment |
| **1177** | **5** | 8 | 55 (37 - 137) | 100% | 70% | none | female | *not done* | *not done* | 62 | 96% | BM | >0,5Mio cells | low transduction efficiency |
| **1183** | 2 | 1 | *no re-engraftment* | *not done* | 0% | none | | *not done* | *not done* | *not done* | 0% | | | |
| **1184** | 2 | 1 | *no re-engraftment* | *not done* | 0% | none | | *not done* | *not done* | *not done* | 0% | | | |
| **1186** | **5** | 5 | 84 (55 - 147) | 70% | 70% | **76%** | male or female | 62 | 93% | 39 | 93% | BM or Spl | >0,5Mio cells | .25+ cells cannot be amplified |
| **1191** | 2 | 1 | *no re-engraftment* | *not done* | 0% | none | | *not done* | *not done* | *not done* | 0% | | | |
| **1221** | 4 | 6 | 105 (74 - 133) | 87% | 60% | none | | *not done* | *not done* | 39 | 87% | BM | | rather slow, low transuction, sorted did not engraft |
| **1222** | **5** | 7 | 50 (35 - 68) | 100% | 90% | **33%** | male or female | 70 | 91% | 64 | 88% | BM or Spl | >0,5Mio cells | low transduction efficiency |
| **1223** | 2 | 3 | 174 | *not done* | 5% | none | | *not done* | *not done* | *not done* | 18% | BM | | no re-engraftment |
| **1224** | 3 | 2 | 144 (93 - 184) | *not done* | 15% | none | | *not done* | *not done* | *not done* | 29% | BM | | slow; no transduction feasible |
| **1225** | **4** | 6 | 104 (73 - 146) | 100% | 50% | none | | *not done* | *not done* | 83 | 90% | BM | | slow |
| **1226** | **5** | 5 | 63 (48 - 114) | 100% | 60% | none | female | *not done* | *not done* | 35 | 93% | BM | >0,5Mio cells | low transduction efficiency |
| **1227** | 2 | 2 | 110 (90 - 130) | 25% | 50% | none | | *not done* | *not done* | 53 | 45% | BM | | no re-engraftment |
| **1228** | 4 | 5 | 86 (70 - 133) | 83% | 80% | none | | *not done* | *not done* | 44 | 83% | BM | | rather slow, transduction difficult |
| **R139** | ≥4 | 5 | 50 (28 - 76) | | | | | | | | | | | |
| **H615** | ≥4 | 4 | 67 (47 - 97) | | | | | | | | | | | |
| **H826** | ≥4 | 5 | 39 (15 - 47) | | | | | | | | | | | |
| **H958** | ≥4 | 4 | 53 (26 - 71) | | | | | | | | | | | |
| **H1454** | ≥4 | 4 | 89 (56 - 167) | | | | | | | | | | | |
| **H1555** | 3 | 5 | 141 (50 - 174) | | | | | | | | | | | |
| **H2788** | ≥4 | 4 | 49 (33 - 74) | | | | | | | | | | | |

Table S1C. Score 2-6 Samples. Mouse data

| AHS-ID | Score | genetic modification | | | |
|---|---|---|---|---|---|
| | | Transduction efficiency | Luc mCherry | Cre-ERT2 dsRed | splitCas9 |
| **295** | 2 | *not done* | *not done* | *not done* | *not done* |
| **343** | 2 | *not done* | *not done* | *not done* | *not done* |
| **346** | **5** | 2 - 50% | *not done* | TRUE | TRUE |
| **356** | **6** | 90% | TRUE | TRUE | TRUE |
| **358** | **5** | 0 - 3% | TRUE | TRUE | TRUE |
| **361** | 4 | FAIL | FAIL | *not done* | *not done* |
| **362** | 2 | *not done* | *not done* | *not done* | *not done* |
| **372** | **6** | 1 - 25% | TRUE | TRUE | FAIL |
| **373** | 2 | *not done* | *not done* | *not done* | *not done* |
| **388** | **5** | 2,30% | TRUE | TRUE | TRUE |
| **392** | 2 | *not done* | *not done* | *not done* | *not done* |
| **393** | **6** | 12-73% | TRUE | TRUE | TRUE |
| **403** | 2 | *not done* | *not done* | *not done* | *not done* |
| **407** | 3 | *not done* | FAIL | *not done* | *not done* |
| **412** | 2 | FAIL | FAIL | *not done* | *not done* |
| **415** | 3 | 0 - 6% | TRUE | *not done* | *not done* |
| **426** | 3 | *not done* | *not done* | *not done* | *not done* |
| **448** | 3 | *not done* | *not done* | *not done* | *not done* |
| **452** | 2 | *not done* | *not done* | *not done* | *not done* |
| **461** | 2 | *not done* | *not done* | *not done* | *not done* |
| **479** | 3 | *not done* | *not done* | *not done* | *not done* |
| **485** | 3 | *not done* | *not done* | *not done* | *not done* |
| **489** | 2 | *not done* | *not done* | *not done* | *not done* |
| **491** | **6** | 11% | TRUE | TRUE | TRUE |
| **506** | 3 | *not done* | *not done* | *not done* | *not done* |
| **530** | 3 | *not done* | *not done* | *not done* | *not done* |
| **538** | **5** | 3,50% | TRUE | TRUE | FAIL |
| **573** | **5** | 0,2 - 0,5% | TRUE | *not done* | FAIL |
| **579** | **5** | 0,1% - 2,5% | TRUE | *not done* | FAIL |
| **596** | 3 | *not done* | *not done* | *not done* | *not done* |
| **601** | 2 | *not done* | *not done* | *not done* | *not done* |
| **602** | **5** | 7 - 30% | TRUE | TRUE | TRUE |
| **613** | 2 | *not done* | *not done* | *not done* | *not done* |
| **625** | 2 | *not done* | *not done* | *not done* | *not done* |
| **640** | **6** | 8 - 9% | TRUE | TRUE | TRUE (unstable) |
| **661** | **5** | 0,7-2,5% | TRUE | TRUE | TRUE |
| **663** | **6** | 22% | TRUE | TRUE | FAIL |
| **664** | **5** | 33% | TRUE | *not done* | *not done* |
| **669** | **5** | 12-20% | TRUE | *not done* | FAIL |
| **760** | 2 | *not done* | *not done* | *not done* | *not done* |
| **788** | 3 | *not done* | *not done* | *not done* | *not done* |
| **810** | 3 | *not done* | *not done* | *not done* | *not done* |
| **842** | 4 | 2% | *not done* | *not done* | *not done* |
| **896** | 4 | 5% | TRUE | *not done* | *not done* |
| **954** | 2 | *not done* | *not done* | *not done* | *not done* |
| **955** | **5** | 5% | TRUE | *not done* | FAIL |
| **957** | 2 | *not done* | *not done* | *not done* | *not done* |
| **958** | 3 | *not done* | *not done* | *not done* | *not done* |
| **959** | 3 | <30% | TRUE | *not done* | *not done* |
| **960** | 3 | *not done* | *not done* | *not done* | *not done* |
| **961** | 2 | *not done* | *not done* | *not done* | *not done* |
| **962** | 2 | *not done* | *not done* | *not done* | *not done* |
| **979** | **6** | $ after 1st sort | TRUE | TRUE | FAIL |
| **980** | 3 | 14-67% after 1st sort | TRUE | *not done* | *not done* |
| **981** | **5** | 1 - 5% | TRUE | TRUE | FAIL |
| **1176** | 2 | *not done* | *not done* | *not done* | *not done* |
| **1177** | **5** | 0 - 2% | TRUE | TRUE | *not done* |
| **1183** | 2 | *not done* | *not done* | *not done* | *not done* |
| **1184** | 2 | *not done* | *not done* | *not done* | *not done* |
| **1186** | **5** | 6% | TRUE | TRUE | FAIL |
| **1191** | 2 | *not done* | *not done* | *not done* | *not done* |
| **1221** | 4 | <1% | FAIL | TRUE | TRUE |
| **1222** | **5** | 2% | TRUE | TRUE | TRUE |
| **1223** | 2 | *not done* | *not done* | *not done* | *not done* |
| **1224** | 3 | *not done* | *not done* | *not done* | *not done* |
| **1225** | **4** | 7% | TRUE | TRUE | TRUE |
| **1226** | **5** | 1% | TRUE | TRUE | TRUE |
| **1227** | 2 | FAIL | FAIL | *not done* | *not done* |
| **1228** | 4 | FAIL | FAIL | TRUE | FAIL |

Table S1D. Score 2-6 Samples. PDX genetic modification

| AHS-ID | Score | molecular characterization (Passage analyzed, PDX or GEPDX) | | | | | | |
|---|---|---|---|---|---|---|---|---|
| | | Cytogenetics | Panel seq | lcWGS | MethArray | scrbSeq Transcript. | RNA seq | Immuno-phenotyp |
| 295 | 2 | not done | not done | not done | not done | not done | not done | not done |
| 343 | 2 | not done | not done | not done | not done | not done | not done | not done |
| 346 | 5 | P5(GEPDX) | P6(GEPDX) | P13(GEPDX) | P11(GEPDX) | not done | P12(GEPDX) | P8(PDX) |
| 356 | 6 | not done | P7(GEPDX) | P9(GEPDX) | P7(GEPDX) | not done | P10(GEPDX) | P8(PDX) |
| 358 | 5 | P11(GEPDX) | P8(PDX) | P8(PDX) | P8(PDX) | not done | not done | P8(PDX) |
| 361 | 4 | not done | P0,1,4(PDX) | not done | not done | P1,2,3,4,5,6(PDX) | not done | not done |
| 362 | 2 | not done | not done | not done | not done | not done | not done | not done |
| 372 | 6 | P4(PDX) | P0(PDX), P6(GEPDX) | P3,5(GEPDX) | P7(GEPDX) | P0,1,2(PDX), P3,4,5,6(GEPDX) | P6(PDX) | P5(PDX) |
| 373 | 2 | not done | P0(PDX) | not done | not done | not done | not done | not done |
| 388 | 5 | P4(PDX) | P0(PDX), P6(GEPDX) | P3,4(GEPDX) | P6(GEPDX) | P0,1,2,3,4(PDX), P4(GEPDX) | P4(PDX) | P6(PDX) |
| 392 | 2 | not done | not done | not done | not done | not done | not done | not done |
| 393 | 6 | P5(PDX) | P0(PDX), P5(GEPDX) | P8,9(GEPDX) | P8(GEPDX) | P0,1,2,3,4,5(PDX), P4,6,7(GEPD | P3(PDX) | P4(PDX) |
| 403 | 2 | not done | not done | not done | not done | not done | not done | not done |
| 407 | 3 | not done | P0(PDX) | not done | not done | P0,1,2,3,4,6(PDX) | not done | not done |
| 412 | 2 | not done | P0,1,2(PDX) | not done | not done | not done | not done | not done |
| 415 | 3 | not done | P1(PDX), P8(GEPDX) | not done | P8(GEPDX) | P0,1,2,3,4,5(PDX) | not done | not done |
| 426 | 3 | not done | not done | not done | not done | not done | not done | not done |
| 448 | 3 | not done | not done | not done | not done | not done | not done | not done |
| 452 | 2 | not done | not done | not done | not done | not done | not done | not done |
| 461 | 2 | not done | not done | not done | not done | not done | not done | not done |
| 479 | 3 | not done | not done | not done | not done | not done | not done | not done |
| 485 | 3 | not done | not done | not done | not done | not done | not done | not done |
| 489 | 2 | not done | not done | not done | not done | not done | not done | not done |
| 491 | 6 | P2(PDX) | P0(PDX), P3,5(GEPDX) | P2,5(GEPDX) | P5(GEPDX) | P0,1,2,3(PDX), P3,4,5,6(GEPDX | P4(PDX) | P3(PDX) |
| 506 | 3 | not done | not done | not done | not done | not done | not done | not done |
| 530 | 3 | not done | not done | not done | not done | not done | not done | not done |
| 538 | 5 | not done | P0(PDX), P3(GEPDX) | P4,6(GEPDX) | P3(GEPDX) | P0,1,2,3(PDX), P3(GEPDX) | P3(PDX) | P3(PDX) |
| 573 | 5 | P8(GEPDX) | P1,2(PDX), P9(GEPDX) | P6,7(GEPDX) | P9(GEPDX) | P0,1,2,3,4,5(PDX) | P2(PDX) | P1(PDX) |
| 579 | 5 | P6(PDX) | P1(PDX), P4(GEPDX) | P2,5(GEPDX) | P4(GEPDX) | P0,1,2,3(PDX), P4,5(GEPDX) | P4,5(PDX) | P4(PDX) |
| 596 | 3 | not done | not done | not done | not done | not done | not done | not done |
| 601 | 2 | not done | not done | not done | not done | not done | not done | not done |
| 602 | 5 | not done | P0(PDX), P8(GEPDX) | P6(PDX), P7(GEPDX) | P8(GEPDX) | P0,1,2,3(PDX), P4,5,6(GEPDX) | P7(GEPDX) | P2(PDX) |
| 613 | 2 | not done | not done | not done | not done | not done | not done | not done |
| 625 | 2 | not done | not done | not done | not done | not done | not done | not done |
| 640 | 6 | P5(GEPDX) | P2(PDX), P4(GEPDX) | P4,7(GEPDX) | P4(GEPDX) | P2(GEPDX) | P2,4(PDX) | P4(PDX) |
| 661 | 5 | P4(PDX) | P0(PDX), P1(GEPDX) | P3,5(GEPDX) | P5(GEPDX) | P1(GEPDX) | P3(PDX) | P3(PDX) |
| 663 | 6 | P6(GEPDX) | P3(GEPDX) | not done | P5(GEPDX) | P4(PDX) | P4(PDX) | P4(PDX) |
| 664 | 5 | not done | P4(GEPDX) | P2(PDX), P4(GEPDX) | P4(GEPDX) | P2(PDX) | P2(PDX) | P2(PDX) |
| 669 | 5 | not done | P2(GEPDX) | P1(PDX), P2(GEPDX) | P2(GEPDX) | P2(PDX) | P1(PDX) | P1(PDX) |
| 760 | 2 | not done | not done | not done | not done | not done | not done | not done |
| 788 | 3 | not done | P1(PDX) | not done | P1(PDX) | not done | not done | not done |
| 810 | 3 | not done | P1(PDX) | not done | not done | not done | not done | not done |
| 842 | 4 | not done | P2(GEPDX) | not done | P2(GEPDX) | not done | not done | not done |
| 896 | 4 | not done | P1(GEPDX) | not done | P1(GEPDX) | P1(GEPDX) | not done | not done |
| 954 | 2 | not done | not done | not done | not done | not done | not done | not done |
| 955 | 5 | P5(GEPDX) | P3(GEPDX) | P2(PDX), P2(GEPDX) | P3(GEPDX) | P2(PDX) | P4(GEPDX) | P4(PDX) |
| 957 | 2 | not done | not done | not done | not done | not done | not done | not done |
| 958 | 3 | not done | not done | not done | not done | not done | not done | not done |
| 959 | 3 | not done | P4(GEPDX) | not done | P4(GEPDX) | P3(PDX) | not done | not done |
| 960 | 3 | not done | not done | not done | not done | not done | not done | not done |
| 961 | 2 | not done | not done | not done | not done | not done | not done | not done |
| 962 | 2 | not done | not done | not done | not done | not done | not done | not done |
| 979 | 6 | P3(GEPDX) | P1(GEPDX) | not done | P1(GEPDX) | P1(PDX) | P2(PDX) | P3(PDX) |
| 980 | 3 | not done | P2(GEPDX) | not done | P2(GEPDX) | not done | not done | not done |
| 981 | 5 | P5(GEPDX) | P2(GEPDX) | P3(GEPDX) | P2(GEPDX) | P1(GEPDX) | P3(PDX), P5(G | P3(PDX) |
| 1176 | 2 | not done | not done | not done | not done | not done | not done | not done |
| 1177 | 5 | P10(GEPDX) | P4(GEPDX) | P4(PDX), P6(GEPDX) | not done | not done | P4(PDX) | P4(PDX) |
| 1183 | 2 | not done | not done | not done | not done | not done | not done | not done |
| 1184 | 2 | not done | not done | not done | not done | not done | not done | not done |
| 1186 | 5 | not done | P3(GEPDX) | P3(PDX) | not done | not done | P3(PDX) | P2(PDX) |
| 1191 | 2 | not done | not done | not done | not done | not done | not done | not done |
| 1221 | 4 | not done | P5(PDX) | P5(PDX) | not done | not done | P4(PDX) | not done |
| 1222 | 5 | not done | P5(PDX) | P5(PDX) | not done | not done | P7(GEPDX) | P4(PDX) |
| 1223 | 2 | not done | not done | not done | not done | not done | not done | not done |
| 1224 | 3 | not done | not done | not done | not done | not done | not done | not done |
| 1225 | 4 | not done | P1(PDX) | P1(PDX) | not done | not done | P1(PDX) | not done |
| 1226 | 5 | P6(GEPDX) | P1,3(GEPDX) | P3,4(PDX) | not done | not done | P3,4(PDX) | P3(PDX) |
| 1227 | 2 | not done | not done | not done | not done | not done | not done | not done |
| 1228 | 4 | not done | P1,4(GEPDX) | P1,4(GEPDX) | not done | not done | P5(GEPDX) | not done |

Table S1E. Score 2-6 Samples. PDX molecular characterization

| AHS-ID | Score | patient data | | | | | | | | | | |
|---|---|---|---|---|---|---|---|---|---|---|---|---|
| | | origin | material | cohort | age | sex | Disease status | risk group ELN 2022 | FAB Subtype | Cytogenetics | Fusions | panel seq (mutations) |
| **281** | 1 | KUM | primary | 1 | 35 | f | R2 | Adverse | M0 | complex | | NA |
| **283** | 1 | KUM | primary | 1 | 68 | m | ID | Intermediate | M5 | CN | | FLT3-ITD, NPM1, DNMT3A |
| **286** | 1 | KUM | primary | 1 | 59 | m | ID | Favorable | M4 | del(13)(q12q22) | CBFß::MYH11 | FLT3-ITD, KIT |
| **294** | 1 | KUM | primary | 1 | 80 | m | ID | Adverse | M1 | CN | | IDH1, SRSF2, MFSD11 |
| **296** | 1 | KUM | primary | 1 | 72 | m | R1, sAML of MPS | Adverse | NA | complex | | IDH1, JAK2, SRSF2, MFSD11 |
| **299** | 1 | KUM | primary | 1 | 60 | f | R1 | Intermediate | M1 | CN | | IDH2, FLT3-ITD, NPM1, DNMT3A |
| **306** | 1 | KUM | primary | 1 | 42 | m | ID | Adverse | M1 | CN | | NPM1, EZH2, IDH2, SRSF2 |
| **308** | 1 | KUM | primary | 1 | 66 | f | ID | Favorable | NA | CN | | FLT3-TDK, NPM1, DNMT3A, IDH1 |
| **330** | 1 | KUM | primary | 1 | 73 | f | R1 | Intermediate | M2 | CN | | NPM1, FLT3-ITD, DNMT3A |
| **331** | 1 | KUM | primary | 1 | 81 | f | ID | Adverse | M2 | complex | | TP53 |
| **333** | 1 | KUM | primary | 1 | 64 | f | ID | Intermediate | M1 | CN | | none identified |
| **344** | 1 | KUM | primary | 1 | 48 | f | ID, tAML | Adverse | M5 | t(9;11)(p22;q23) | KMT2A::MLLT3 | BRAF |
| **345** | 1 | KUM | primary | 1 | 54 | f | ID, tAML | Adverse | M1 | CN | | FLT3-ITD, RUNX1, TET2 |
| **348** | 1 | KUM | primary | 1 | 58 | m | ID | Intermediate | M4 | CN | | NPM1, FLT3-ITD, DNMT3A |
| **391** | 1 | KUM | primary | 1 | 62 | m | ID, sAML of MDS/MPS | Adverse | M1 | ´+13 | | SRSF2, SETBP1, ASXL1, RAD21, NF1, CBL, NF1, NF1, RUNX1, ASXL2 |
| **396** | 1 | KUM | primary | 1 | 73 | m | ID /CMML | Intermediate | M4 / M5 | ´+8 | | NA |
| **401** | 1 | KUM | primary | 1 | 73 | m | ID | Favorable | M1 | der(14)t(11;14)(q1?3;q3?2) | | FLT3-TDK, NPM1, DNMT3A |
| **404** | 1 | KUM | primary | 1 | 59 | m | ID | Intermediate | M4 | CN | | FLT3-ITD, NPM1, DNMT3A |
| **405** | 1 | KUM | primary | 1 | 51 | m | ID | Intermediate | M1 | CN | | FLT3-ITD, FLT3-TDK, NPM1, DNMT3A, NFE2, WT1, PHF6 |
| **408** | 1 | KUM | primary | 1 | 72 | f | ID | Favorable | M4 | CN | | NPM1, DNMT3A, TET2, TET2-AS1 |
| **410** | 1 | KUM | primary | 1 | 35 | m | ID | Favorable | M4 | CN | | NPM1, DNMT3A |
| **421** | 1 | KUM | primary | 1 | 47 | f | ID | Favorable | M1 | CN | | NPM1, ZBTB7A, NRAS, FLT3-TKD |
| **433** | 1 | KUM | primary | 1 | 48 | f | ID | Favorable | M4 | CN | | NPM1, DNMT3A, IDH2 |
| **443** | 1 | Dresden | primary | 2 | *unkn* | m | R | *unknown* | *unknown* | *unknown* | *unknown* | *unknown* |
| **478** | 1 | KUM | primary | 1 | 72 | m | ID | Favorable | M5 | CN | | NPM1, DNMT3A, NRAS |
| **493** | 1 | KUM | primary | 1 | 21 | f | ID | Intermediate | M5 | CN | | NPM1, FLT3-ITD, DNMT3A |
| **505** | 1 | Freiburg | primary | 2 | *unkn* | f | *unknown* | *unknown* | *unknown* | +8, 5q Del, -20, TP53-Del | *unknown* | NPM1, FLT3-ITD, CEBPA |
| **518** | 1 | KUM | primary | 1 | 75 | f | ID | Favorable | M1 | CN | | NPM1, FLT3, DNMT3A, TET2, TET2-AS1 |
| **522** | 1 | KUM | primary | 1 | 67 | f | ID | Favorable | M4 | CN | | NPM1, TET2, DNMT3A, KRAS, PTPN11 |
| **525** | 1 | KUM | primary | 1 | 57 | m | primary refractory | Adverse | NA | 5q31-Deletion | | RUNX1, PTPN11, SF3B1 |
| **534** | 1 | Freiburg | primary | 2 | 70 | m | *unknown* | *unknown* | *unknown* | complex, including del5q31, del13q14 | *unknown* | *unknown* |
| **551** | 1 | TUM | primary | 2 | 54 | f | ID | *unknown* | *unknown* | *unknown* | *unknown* | FLT3-TKD, NPM1 |
| **552** | 1 | TUM | primary | 2 | 31 | f | ID | *unknown* | *unknown* | 47, XX, +19 | none detected | RUNX1 |
| **577** | 1 | KUM | primary | 1 | 74 | f | primary refractory | Adverse | M1 | NA | | DNMT3A, IDH2, STAG2, PHF6, BCOR |
| **580** | 1 | KUM | primary | 1 | 24 | m | R1 | Adverse | M4 | complex | KMT2A::MLLT3 | KRAS, WT1 |
| **587** | 1 | Freiburg | primary | 2 | 70 | m | *unknown* | *unknown* | *unknown* | *unknown* | *unknown* | *unknown* |
| **609** | 1 | Freiburg | primary | 2 | 68 | m | *unknown* | *unknown* | *unknown* | *unknown* | *unknown* | *unknown* |
| **616** | 1 | Berlin | primary | 2 | 63 | f | R1 | *unknown* | *unknown* | *unknown* | *unknown* | *unknown* |
| **618** | 1 | KUM | primary | 1 | 29 | m | ID | Adverse | M5 | ´+8,t(9;11)(p22;q23) | KMT2A::MLLT3 | FLT3-TDK, KRAS |
| **619** | 1 | KUM | primary | 1 | 44 | f | ID | Adverse | M5 | t(9;11)(p22;q23) | KMT2A::MLLT3 | none identified |
| **620** | 1 | Berlin | primary | 2 | 62 | m | primary refractory | *unknown* | *unknown* | CN | *unknown* | *unknown* |
| **623** | 1 | KUM | primary | 1 | 24 | f | ID | Adverse | M5 | t(9;11)(p22;q23) | KMT2A::MLLT3 | KRAS, NRAS |
| **635** | 1 | Freiburg | primary | 2 | *unkn* | m | primary refractory | *unknown* | *unknown* | *unknown* | *unknown* | *unknown* |
| **662** | 1 | KUM | primary | 1 | 30 | f | primary refractory | Intermediate | M2 | CN | | none identified |
| **667** | 1 | KUM | primary | 1 | 78 | f | ID | Adverse | NA | ´-7 | | TET2, CEBPA |
| **676** | 1 | Berlin | primary | 2 | 70 | 7 | R1 | *unknown* | *unknown* | *unknown* | *unknown* | *unknown* |
| **683** | 1 | KUM | primary | 1 | 66 | f | ID | Adverse | M5 | t(9;11)(p22;q23) | KMT2A::MLLT3 | KRAS, BCOR, SRSF2 |
| **689** | 1 | KUM | primary | 1 | 73 | m | ID | Adverse | M7 | complex including +8, +19, +21 | | JAK2, MPL, DNMT3A, ZRSR2, TET2 |
| **744** | 1 | KUM | primary | 1 | 36 | m | ID | Favorable | M2 | t(8;21)(q22;q22) | RUNX1::RUNX1T1 | RAD21, ASXL2, KIT |
| **771** | 1 | KUM | primary | 1 | 57 | f | ID | Favorable | M1 | CN | | DCLK1, CUX1, STAG2, GATA2, NPM1, TET2 |
| **785** | 1 | KUM | primary | 1 | 70 | m | ID | Intermediate | M1 | ´+13 | | DUP-MLL |
| **898** | 1 | TUM | primary | 2 | 52 | f | refractory | *unknown* | *unknown* | CN | *unknown* | DNMT3A, IDH2, NPM1, FLT3-ITD, ASXL2 |
| **942** | 1 | KUM | primary | 1 | 59 | f | R1 | Favorable | M1 | CN | | STAG2, NPM1, TET2 |
| **952** | 1 | KUM | primary | 1 | 52 | f | ID | Adverse | M2 | CN | | ETV6, DNMT3A, CUX1, BCOR, RUNX1, PTPN11 |
| **1140** | 1 | KUM | primary | 1 | 42 | f | ID | Adverse | NA | t(9;11)(p22;q23) | KMT2A::MLLT3 | NRAS, PTPN11, TET2 |
| **H141** | 1 | FFM | PDX | 4 | 6 | m | ID | | | t(10;11) | | |
| **H945** | 1 | FFM | PDX | 4 | 2 | m | ID | | | t(9;11)(p22;q23) | | |
| **H1063** | 1 | FFM | PDX | 4 | 11 | m | ID | | | 11q23-aberration (except of t(4;11), t(11;19), t(9;11)) | | |
| **H1169** | 1 | FFM | PDX | 4 | 4 | f | ID | | | t(9;11)(p22;q23) | | |
| **H1369** | 1 | FFM | PDX | 4 | 1 | f | ID | | | 11q23-aberration (except of t(4;11), t(11;19), t(9;11)) | | |
| **H1569** | 1 | FFM | PDX | 4 | 3 | f | ID | | | t(10;11) | | |

Table S1F. Score 1 Samples. Patient data

**Supplemental Table 2: Patient characteristics of cohort 1**

Continuous variables were analyzed with the Mann-Whitney-U-Test, and the categorical variables with Fisher's exact text.

| **Parameter** | **Value** | **Engrafted (n = 31)** | **Not engrafted (n = 43)** | ***p*-value** |
|---|---|---|---|---|
| **Age** | Median [range] | 55 [20 - 81] | 59 [21 - 81] | 0.5 |
| **Gender female** | n (%) | 22 (71) | 24 (56) | 0.23 |
| **Relapse** | n (%) | 15 (48) | 9 (21) | 0.022 |
| **WBC (G/L)** | Median [range] | 21.2 [0.2 - 288] | 25.5 [0.1 – 233.0] | 0.97 |
| **Hb (g/dL)** | Median [range] | 10.0 [5.2 – 14.4] | 9.3 [5.4 – 15.6] | 0.24 |
| **Platelets (G/L)** | Median [range] | 46.0 [5.0 – 380.0] | 54 [7.0 – 190.0] | 0.25 |
| **Cytogenetically normal** | n (%) | 13 (57) | 23 (56) | >0.999 |
| **Complex karyotype** | n (%) | 3 (12) | 5 (12) | >0.999 |
| **FAB M0** | n (%) | 0 (0) | 1 (3) | 0.89 |
| **FAB M1** | n (%) | 9 (33) | 14 (37) | |
| **FAB M2** | n (%) | 5 (19) | 5 (13) | |
| **FAB M4** | n (%) | 9 (33) | 9 (24) | |
| **FAB M5** | n (%) | 4 (15) | 8 (21) | |
| **FAB M7** | n (%) | 0 (0) | 1 (3) | |
| **ELN 2022 favorable** | n (%) | 3 (10) | 13 (30) | 0.11 |
| **ELN 2022 intermediate** | n (%) | 10 (32) | 11 (26) | |
| **ELN 2022 adverse** | n (%) | 18 (58) | 19 (44) | |

**Supplemental Table 3: Publications with *PDX models**

| Year | first author(s) | last author(s) | Journal | PMID: | included AML PDX samples | Figures / Tables | ex vivo trials | | | | | | functional in vivo trials | | | | molecular in vivo trials | | | | | | in vivo therapy trials | | | |
|---|---|---|---|---|---|---|---|---|---|---|---|---|---|---|---|---|---|---|---|---|---|---|---|---|---|---|
| | | | | | | | phenotypic characterization | drug testing | CFU assay | growth characteristics | knockdown of GOI | knockout of GOI | LIC assays | re-engr. | dormancy | homing | barcoding | competitive growth | expression of GOI | knockdown of GOI | knockout of GOI | CRISPR screen | chemo-therapy | targeted therapy | immuno-therapy | ADC |
| 2025 | Damaskou | Vassiliou | Blood | 40561247 | 1222 | F4, FS6 | | | | | | | | | | | | | | | | | DONE | | | |
| 2025 | Hosseini | Shi | Nature | 40240608 | 372, 579 | 5i, S9b-g | | | | | | | | | | | | | | | | | | DONE | | |
| 2025 | Huber | Pabst | Hemasphere | 40265168 | 346, 372, 491, 602, 661, 663 | 2A-H, 6B-D | | | | | | | | | | | | | | DONE | | | | | | |
| 2025 | Sollier, Riedel, Toprak | Plass, Lipka | Blood Cancer Discovery | 40162972 | 661 | F7, S12 | | | | | DONE | | | DONE | | | | DONE | | DONE | | | | | | |
| 2024 | Able | Spiekermann | Leukemia | 39870768 | 388, 393, 669, 1186 | F5, F6, S5, S6, TS4, TS5 | | DONE | DONE | | | | | DONE | | | | | | | | | | | | DONE |
| 2024 | Atar | Seitz | Leukemia | 39095503 | 663 (P21) | F6 | | | | | | | | | | | | | | | | | | | DONE | |
| 2024 | Barbosa, Deshpande | Deshpande | Blood | 38048593 | 393 | F7D,E,F. FS7 | | | | DONE | | | | | | | | | | | DONE | | | | | |
| 2024 | Sperk | Eichner, Bassermann | Leukemia | 39014197 | 372, 388, 393, 491,640, 661, 669 | 2C, S5D | DONE | | | | | | | | | | | | | | | | | | | |
| 2023 | Bahrami, Schmid | Jayavelu, Jeremias | Molecular Cancer | 37422628 | 415, 579, 573, 491, 661, 388, 393, 372, 602, 356, 663 | 3D-G, 6D-K | | | DONE | DONE | | DONE | | | | | | DONE | | | DONE | | DONE | | | |
| 2023 | Ghalandary, Gao | Jeremias | Blood | 36256915 | 346, 356, 388, 393, 661, 602, 640 | F1D-F, F2, FS2-S6, FS10-S15 | | | | DONE | | DONE | DONE | DONE | | DONE | | DONE | | | DONE | DONE | DONE | | | |
| 2023 | Gottschlich, Thomas, Grünmeier | Marr, Kobold | Nature Biotechnology | 36914885 | 372, 388, 573 | F5g-m, FS5h-m, FS6a-e, TS1b | DONE | | | | | | | | | | | | | | | | | | DONE | |
| 2023 | Roas | Jeremias, Leonhardt, Spiekermann | Blood | 35981498 | 573, 579, 640 | F4 / F6 / FS6 | DONE | DONE | | | | | | | | | | | | | | | | | | DONE |
| 2022 | Janjic | Enard | Genome Biology | 35361256 | 361, 372, 388, 393, 407, 415, 491, 538, 573, 579, 602 | F5 | DONE | | | | | | | | | | | | | | | | | | | |
| 2022 | Janssen | Dietrich | Blood | 35857899 | 372 | F6ab | | | | | | | | | | | | | | | | | | DONE | | |
| 2022 | Koschade | Brandts | Leukemia | 35999260 | 573, 579, 981 | FS16 | | DONE | | | | | | | | | | | | | | | | | | |
| 2022 | Lopez-Millan | Menéndez | Cancers | 35326743 | 640 | F3 c-e | | | | | | | | | | | | | | | | | | DONE | | |
| 2022 | Sun | Niehrs | Cell Reports | 34407399 | 661 | F4A-E / F6G-J | | | | DONE | | | | | | | | | | DONE | | | | | | |
| 2022 | Zeller | Jeremias | J Hematol Oncol | 35279202 | 491, 661 | F1 / F2 / TS1 | DONE | | | | | | DONE | | | | DONE | DONE | | | | | DONE | | | |
| 2022 | Zhou, Aroua | Müller-Tidow | Cancer | 36259929 | 491, 661 | 2G-I | | | | | | | DONE | | | | | | DONE | | | | | | | |
| 2021 | Augsberger | Subklewe | Blood | 34280257 | 573 | F7D,E | | | | | | | | | | | | | | | | | | | DONE | |
| 2021 | Carlet, Völse | Jeremias | Nat Commun | 34580292 | 388, 393, 491 | F1-3 | | | | | DONE | | | | | | | DONE | | DONE | | | DONE | DONE | | |
| 2021 | Chen | Deshpande | Blood | 33690798 | 393 (patient-derived xenograft with MLL-AF10) | F7E,F (CFU) | | DONE | | | | | | | | | | | | | | | | | | |
| 2021 | Díaz de la Guardia | Menéndez | Blood Adv | 34470043 | 1186 (14048) | F1 / F2 / | | | | | | | | | | | | | | | | | | | | |
| 2021 | Kempf | Spiekermann | Sci Rep | 33712646 | 491, 661 | F7 / F8 | DONE | DONE | | | | | | | | | | | | | | | DONE | | | |
| 2021 | Khateb | Deshpande, Ronai | Nat Commun | 34518534 | 669 | F2J,K | | | | | | | | | | | | | | DONE | | | | | | |
| 2021 | Meßner | Pachmayr | Sci Rep | 34045646 | 346, 372, 388, 393, 491 | F3 / TS1 | | DONE | | | | | | | | | | | | | | | | | | |
| 2021 | Tahk | Hopfner | J Hematol Oncol | 34579739 | 491, 579 | F5F / F6 / F7 / FS3 / FS4 / FS5 | | DONE | | | | | DONE | | | | | | | | | | | | | |
| 2021 | Yankova | Tzelepis, Kouzarides | Nature | 33902106 | 579 (PDX-1), 388 (PDX-2), 393 (PDX-3) | F4 | | | | | | | | DONE | | | | | | | | | | DONE | | |
| 2020 | Baroni | Menéndez | J Immunother | 32527933 | 579 | F2G-J | | | | | | | | | | | | | | | | | | | DONE | |
| 2020 | Ebinger | Jeremias | Haematologica | 32029508 | 346, 356, 358, 372, 388, 393, 491, 538, 579, 661 | F1 - 3 / TS1 - 3 / FS1 - 5 | | | | | | | DONE | | DONE | DONE | | | | | | | DONE | | | |
| 2020 | Jensen | Fröhling | Leukemia | 32591645 | 388, 393 | F2H / F4D | | | DONE | | | | | | | | | | | DONE | | | | | | |
| 2020 | Redondo-Monte | Greif | Oncogene | 32115572 | 346, 393, 491, 573, 579, 640 | FS5 | | DONE | | | | | | | | | | | | | | | | | | |
| 2020 | Salik | Wang | Cancer Cell | 32559496 | 491 | F3A-E, FS4H-J, FS6 | DONE | DONE | | | | | | | | | | | | | DONE | | | DONE | | |
| 2020 | Stief | Spiekermann | Leukemia | 31201358 | 372, 393, 407, 415, 491, 538, 573, 579 | F2 | DONE | DONE | | | | | | | | | | | | | | | DONE | | | |
| 2019 | Garg | Pabst | Blood | 31076446 | 491 | F5D / FS7 | DONE | | | | | | | DONE | | | | | | DONE | | | | | | |
| 2019 | Lynch | Wang | Leukemia | 30622285 | 491 | F5 | | | | | | | | | | | | | | | | | | DONE | | |
| 2019 | Ruzicka | Rothenfusser | Leukemia | 31740809 | 491 | F6 | | | | | | | | | | | | | | | | | | | DONE | |
| 2018 | Koczian | Braig | Haematologica | 30309851 | 372, 393, 491 | F1 | | DONE | | | | | | | | | | | | | | | | | | |
| 2018 | Reiter | Greif | Leukemia | 28895560 | 372, 415, 491, 573, 579, 640 | F4 / F5 / F7 | DONE | DONE | | | | | | | | | | | | | | | | | | |
| 2018 | Tzelepis | Vassiliou | Nat Commun | 30568163 | 393, 491, 579 | F1J,K / F2D,E | | | DONE | | | | | | | | | | | | | | | DONE | | |
| 2017 | Garz | Götze | Oncotarget | 29312564 | 361, 602 | F6 - 7 | | | | | | | | DONE | | | | | | | | | | | | |
| 2016 | Krupka | Subklewe | Leukemia | 26239198 | 346, 361 | F3 | | | | | | | | DONE | | | | | | | | | | | | |
| 2015 | Sandhöfer | Spiekermann | Leukemia | 25322685 | 393 | F7B | | DONE | | | | | | | | | | | | | | | | | | |
| 2015 | Zhang | Liebl | Oncotarget | 26496038 | 372, 412 | F4 / F7 | | DONE | | | | | | | | | | | | | | | | | | |
| 2015 | Vick | Jeremias | PLoS One | 25793878 | 346, 356, 361, 372, 373, 390, 393, 407, 412, 415, 491 | T1 / F1 - 5 / TS 2 - 6 / FS 1 - 5 | | | | | | | DONE | | | | | | | | | | | | | |

Table S3. Publications with AML PDX models

### Supplemental Figures and Legends

#### Supplemental Figure 1

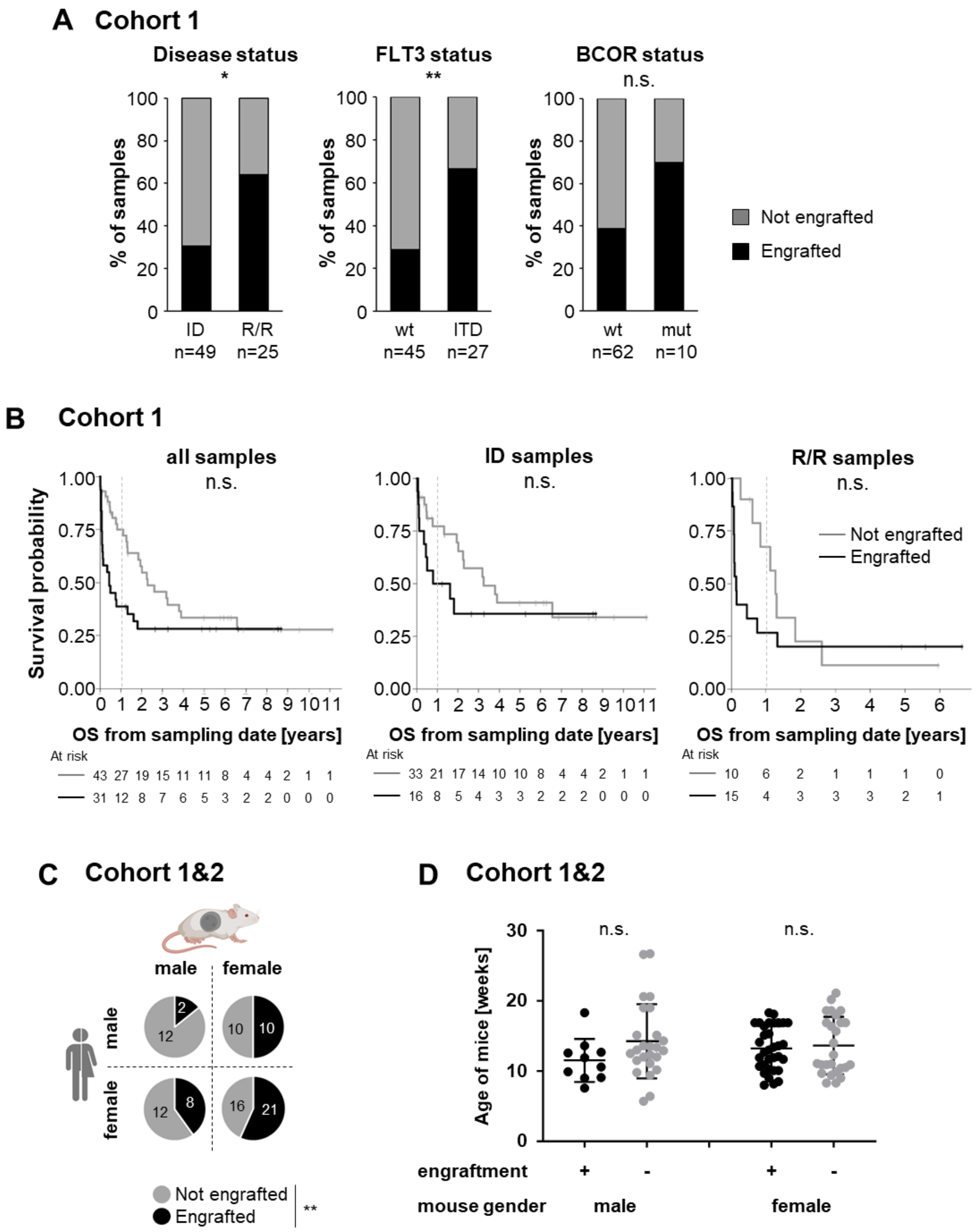

**Supplemental Figure 1. Primary engraftment of primary AML samples in NSG mice correlates with clinical parameters and sex of recipient mice; related to Figure 1A.**

**A-D** Primary patients' or PDX AML cells (2.5E5 – 2E7) were transplanted into one to eight male or female NSG mice via tail vein injection. Positive engraftment was analyzed by flow cytometry of peripheral blood and bone marrow and/or spleen at experimental end point. If at least one mouse showed positive engraftment, the sample was classified as engrafted.

**A,B** Patient samples of cohort 1 (n=74) were analyzed concerning factors influencing primary engraftment.

**A** Bar plots show positive engraftment rate according to disease status (initial diagnosis (ID) versus relapsed / refractory disease (R/R), left, *p*=0.026), *FLT3* mutation status (*FLT3* wildtype (wt) versus *FLT3*-ITD, middle, *p*=0.0034) and *BCOR* mutation status (wt versus mutated (mut), right, *p*=0.084), respectively. The *p*-values were calculated with Fisher's exact test.

**B** Kaplan-Meier graph depicting the overall survival probability stratified by positive (black) or negative (grey) engraftment of primary patient cells. Left: all samples (n=74); HR=1.73, 95%-CI [0.97, 3.07], *p*=0.063; middle: patients at ID (n=49), HR=1.55, 95%-CI [0.71, 3.], *p*=0.28; right: patients with R/R disease (n=25), HR=1.64, 95%-CI [0.66, 4.06], *p*=0.28.

**C** Pie charts displaying positive (black sector) or negative (grey sector) engraftment for primary samples of male or female patients (cohort 1 and 2) transplanted into male or female mice; only samples with known patient gender are included (n=89). Two samples were transplanted both in male and female mice and are therefore counted twice. Univariable analysis for the association between male gender (mouse) and primary engraftment, OR=0.27, *p*=0.0042.

**D** Positive engraftment (black dots) and negative engraftment (grey dots) are depicted for male and female mice of different age at time of injection. One dot represents one sample. n.s.: not significant as tested by two-sided unpaired t-test with Welch´s correction.

**Supplemental Figure 2**

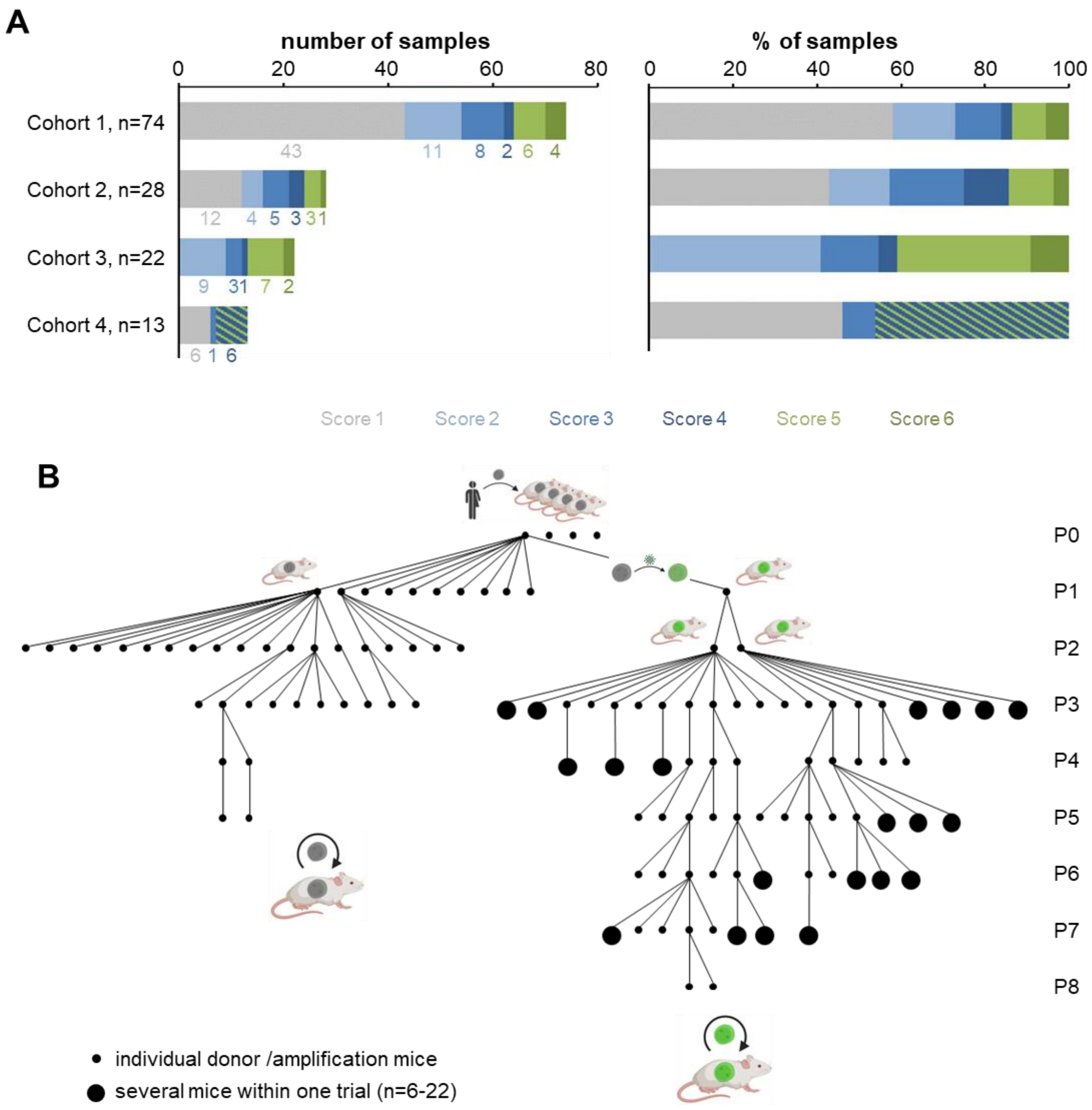

C

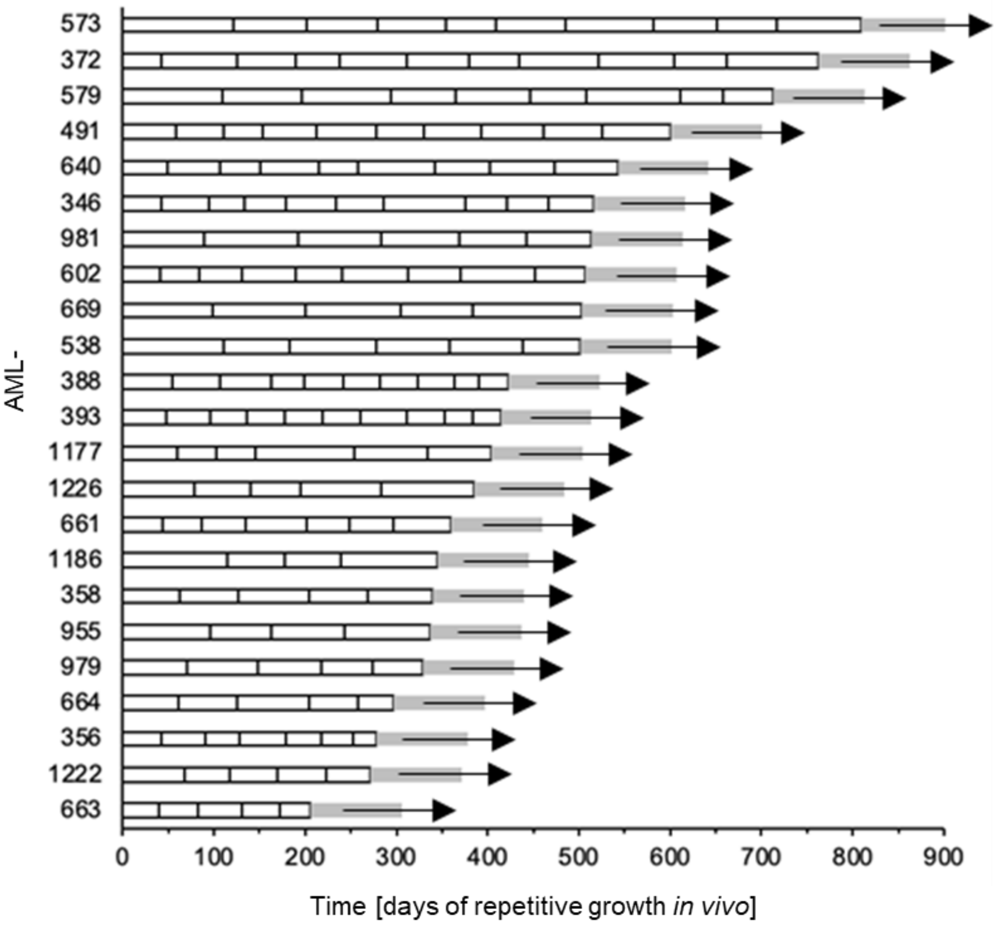
573
372
579
491
640
346
981
602
669
538
388
393
1177
1226
661
1186
358
955
979
664
356
1222
663
AML-
0
100
200
300
400
500
600
700
800
900
Time [days of repetitive growth *in vivo*]

D

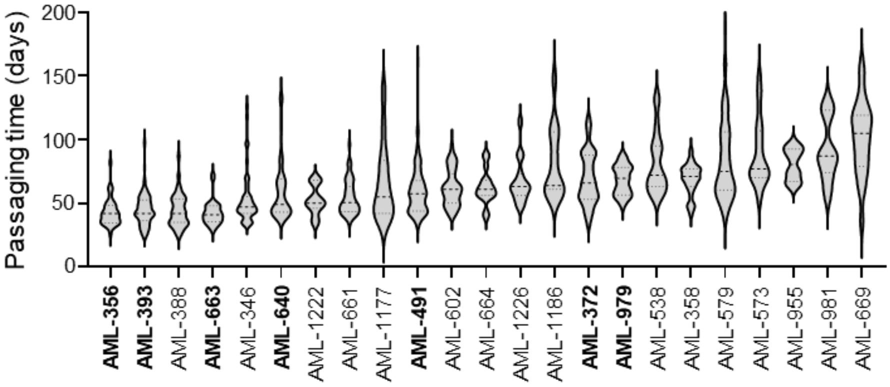
Passaging time (days)
200
150
100
50
0
**AML-356**
**AML-393**
AML-388
**AML-663**
AML-346
**AML-640**
AML-1222
AML-661
AML-1177
**AML-491**
AML-602
AML-664
AML-1226
AML-1186
**AML-372**
**AML-979**
AML-538
AML-358
AML-579
AML-573
AML-955
AML-981
AML-669

**E**

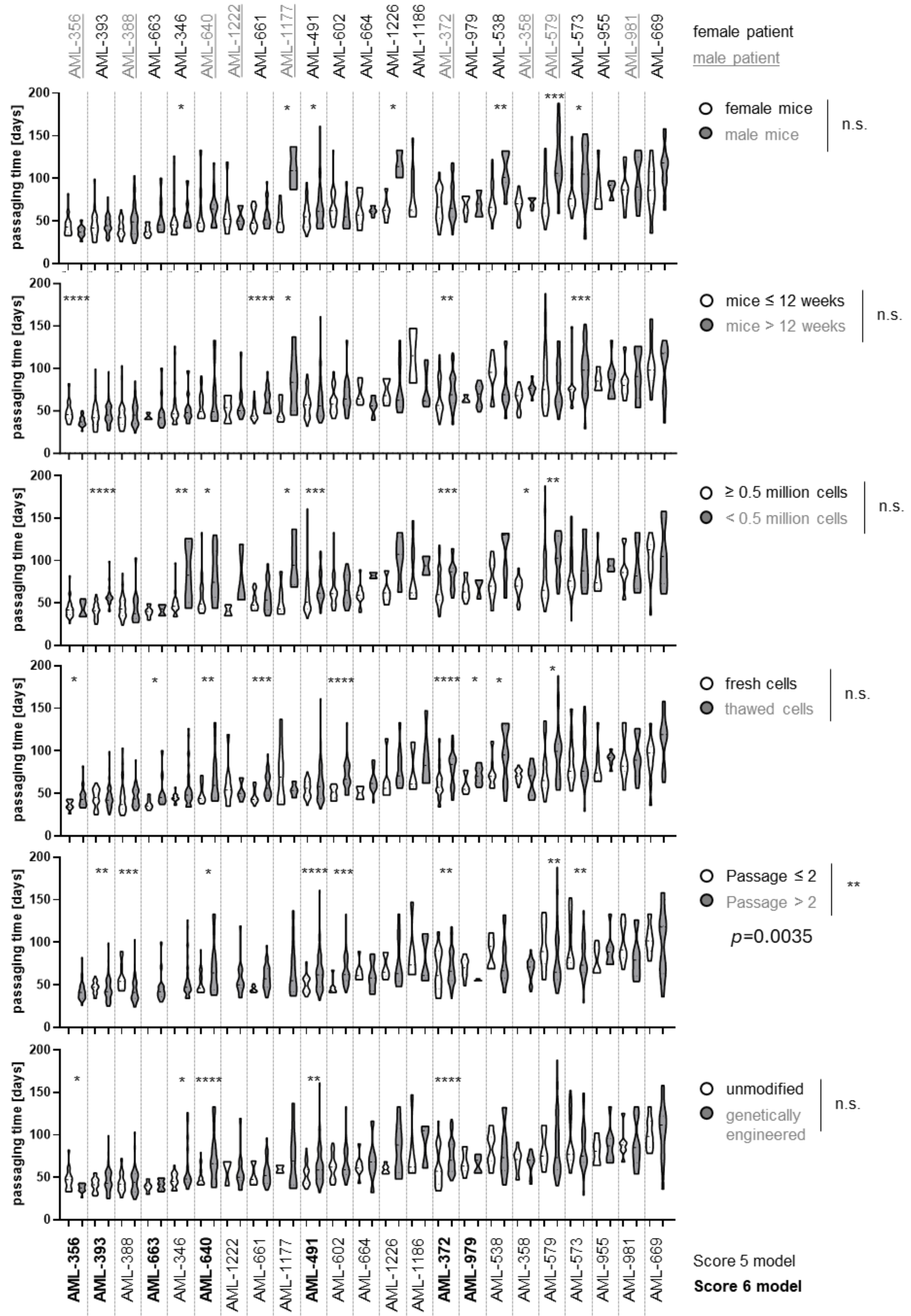

AML-356
AML-393
AML-388
AML-663
AML-346
AML-640
AML-1222
AML-661
AML-1177
AML-491
AML-602
AML-664
AML-1226
AML-1186
AML-372
AML-979
AML-538
AML-358
AML-579
AML-573
AML-955
AML-981
AML-669
female patient
male patient
passaging time [days]
female mice
male mice
n.s.
mice ≤ 12 weeks
mice > 12 weeks
n.s.
≥ 0.5 million cells
< 0.5 million cells
n.s.
fresh cells
thawed cells
n.s.
Passage ≤ 2
Passage > 2
**
p=0.0035
unmodified
genetically engineered
n.s.
Score 5 model
Score 6 model

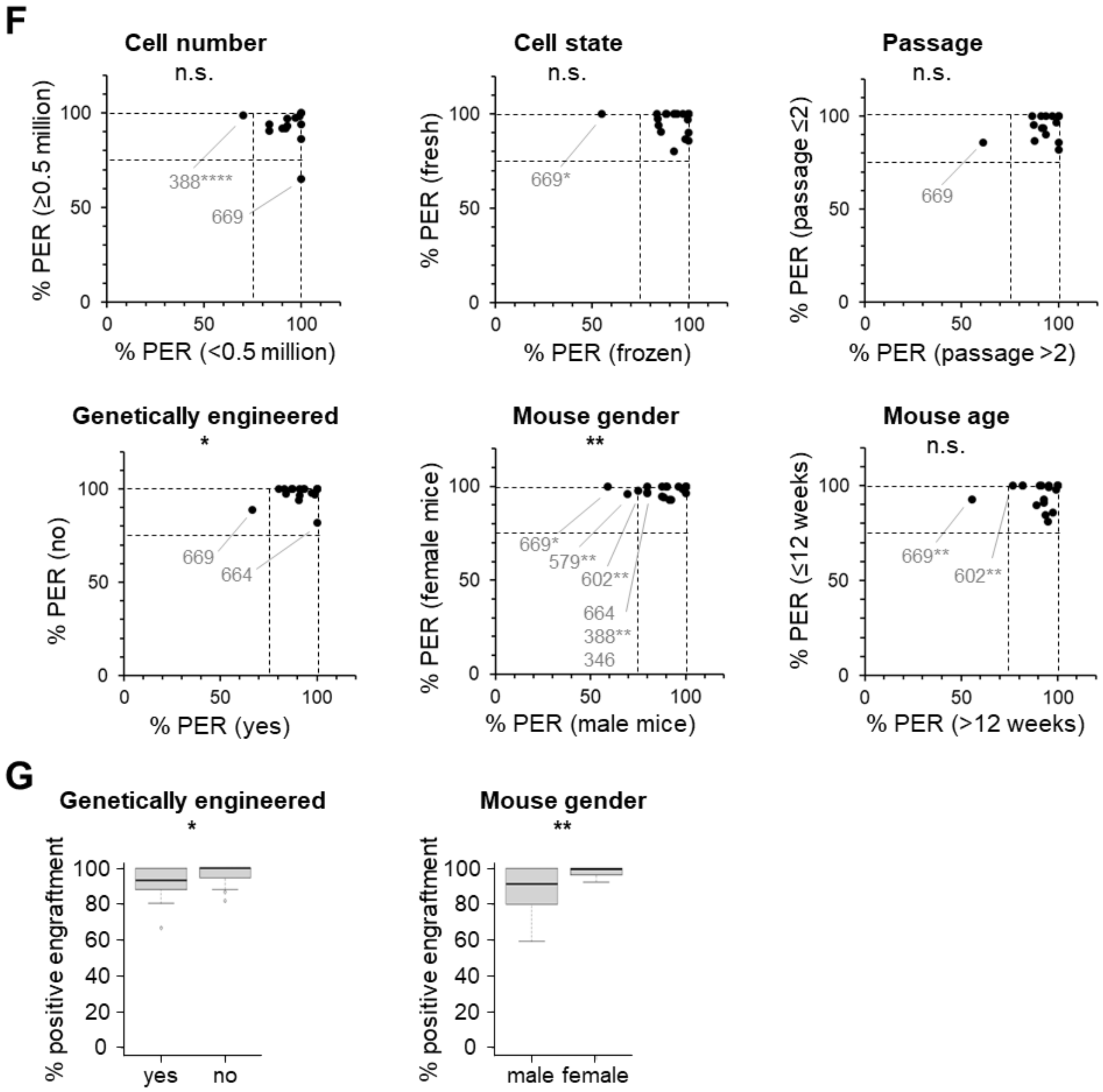
F
Cell number
n.s.
% PER (≥0.5 million)
% PER (<0.5 million)
388****
669
Cell state
n.s.
% PER (fresh)
% PER (frozen)
669*
Passage
n.s.
% PER (passage ≤2)
% PER (passage >2)
669
Genetically engineered
*
% PER (no)
% PER (yes)
669
664
Mouse gender
**
% PER (female mice)
% PER (male mice)
669*
579**
602**
664
388**
346
Mouse age
n.s.
% PER (≤12 weeks)
% PER (>12 weeks)
669**
602**
G
Genetically engineered
*
% positive engraftment
yes
no
Mouse gender
**
% positive engraftment
male
female

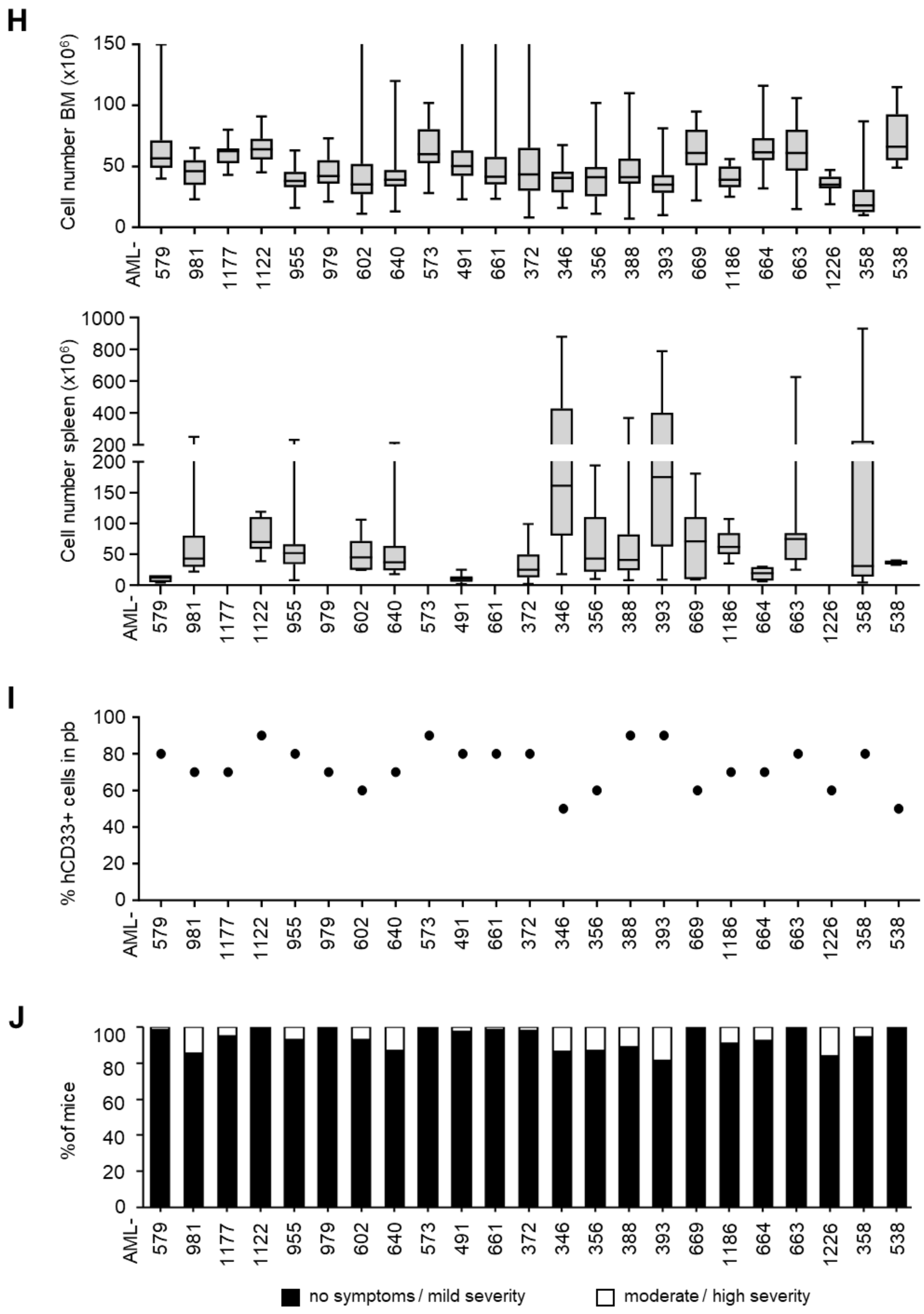

H
Cell number BM (x10⁶)
Cell number spleen (x10⁶)
AML-
579 981 1177 1122 955 979 602 640 573 491 661 372 346 356 388 393 669 1186 664 663 1226 358 538
I
% hCD33+ cells in pb
J
% of mice
no symptoms / mild severity
moderate / high severity

**Supplemental Figure 2.** ***In vivo*** **characteristics of *PDX AML models; related to Figure 1B/C.**

**A Related to Figure 1C.** Distribution of samples collected from the different cohorts within Score 1 – Score 6.

**B-J** *PDX AML models (n=23) were serially passaged in NSG mice for amplification (donor mice), to generate genetically engineered models, and used for *in vivo* trials.

**B Exemplary schematic phylogenetic tree** of one model (AML-491) re-transplanted several times into next recipient mice. Each small dot represents one donor mouse for amplification, each large dot an *in vivo* trial with several mice (n=6-22), each line a re-transplantation step of PDX cells. Left part displays passaging of unmodified PDX cells, right part displays passaging of genetically engineered PDX cells. P: passage.

**C** *PDX AML models show **indefinite growth *in vivo***. Median passaging time of each model at each re-transplantation step (passage) was calculated to depict total time each model had been growing in mice. Grey bars at the end of each time bar indicate the possibility for further re-transplantation steps. Samples are ordered according to total days of *in vitro* growth.

**D** Violin plot depicting the **passaging time** for all donor mice transplanted with Score 5 or Score 6 (bold) PDX models; median indicated by dashed line and $25^{th}$ and $75^{th}$ percentile by dotted lines. Samples are ordered according to median passaging time.

**E Passaging time** was determined in donor mice serially transplanted with *PDX AML models and is depicted for each model concerning mouse gender (female or male), mouse age at time of cell transplantation (younger or older than 12 weeks), cell number injected (more or less than 0.5 million cells), cell state (fresh isolated from donor mice or thawed), cells of low (P≤2) or high re-passaging step (P>2), and genetic engineering (unmodified or genetically engineered cells). Violin plot with median indicated by dashed line and $25^{th}$ and $75^{th}$ percentile by dotted lines. * $p$<0.05, ** $p$<0.01, *** $p$<0.001, **** $p$<0.0001 as analyzed by Fisher's exact test (binary variables) or by Mann-Whitney-U-test (continuous variables), both for individual models and all models at once.

**F Positive engraftment rate (PER)** was analyzed for mice injected with low or high cell numbers (more or less than 0.5 million cells, n.s.), freshly isolated or frozen/thawed cells (n.s.), cells at low or high passage numbers (n.s.), unmodified (no) or genetically engineered (yes) cells (*p*=0.014), and for cells injected into male or female mice (*p*=0.001) or into younger or older mice (below or above 12 weeks, n.s.). Dotted lines indicate PER between 75% and 100%. n.s. not significant, * *p*<0.05, ** *p*<0.01, **** *p*<0.0001 as analyzed by Fisher's exact test (binary variables) or by Mann-Whitney-U-test (continuous variables), both for individual models and across all models.

**G** Box plots showing **percentage of engrafted mice** transplanted with genetically engineered (yes) or unmodified (no) cells (left, n=23 models, *p*=0.042) and for male or female mice (right, n=22 models, *p*=0.001). Statistical significance was analyzed by Wilcoxon test.

**H** Cell numbers of AML PDX models re-isolated from bone marrow (BM) or spleen (only assessed when enlarged at the time point of sacrifice) of donor mice at end-stage leukemia are depicted.

**I** Highest value of hCD33+ cells in peripheral blood (pb) when mice had to be sacrificed due to mild clinical signs of illness.

**J** Percentage of mice per model with no or mild symptoms (black) or showing moderate or severe symptoms (white) at advanced leukemic stage.

#### Supplemental Figure 3

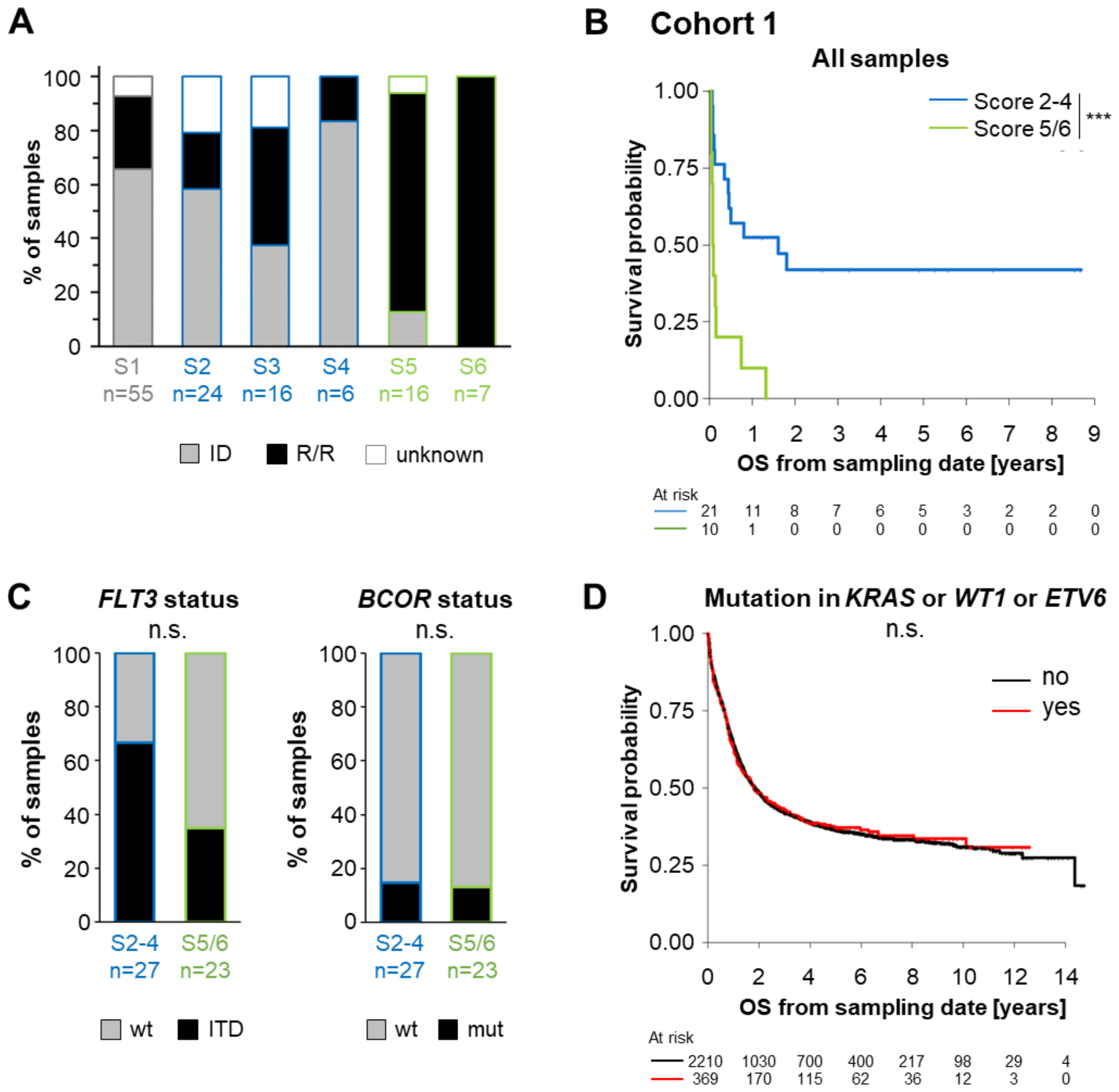

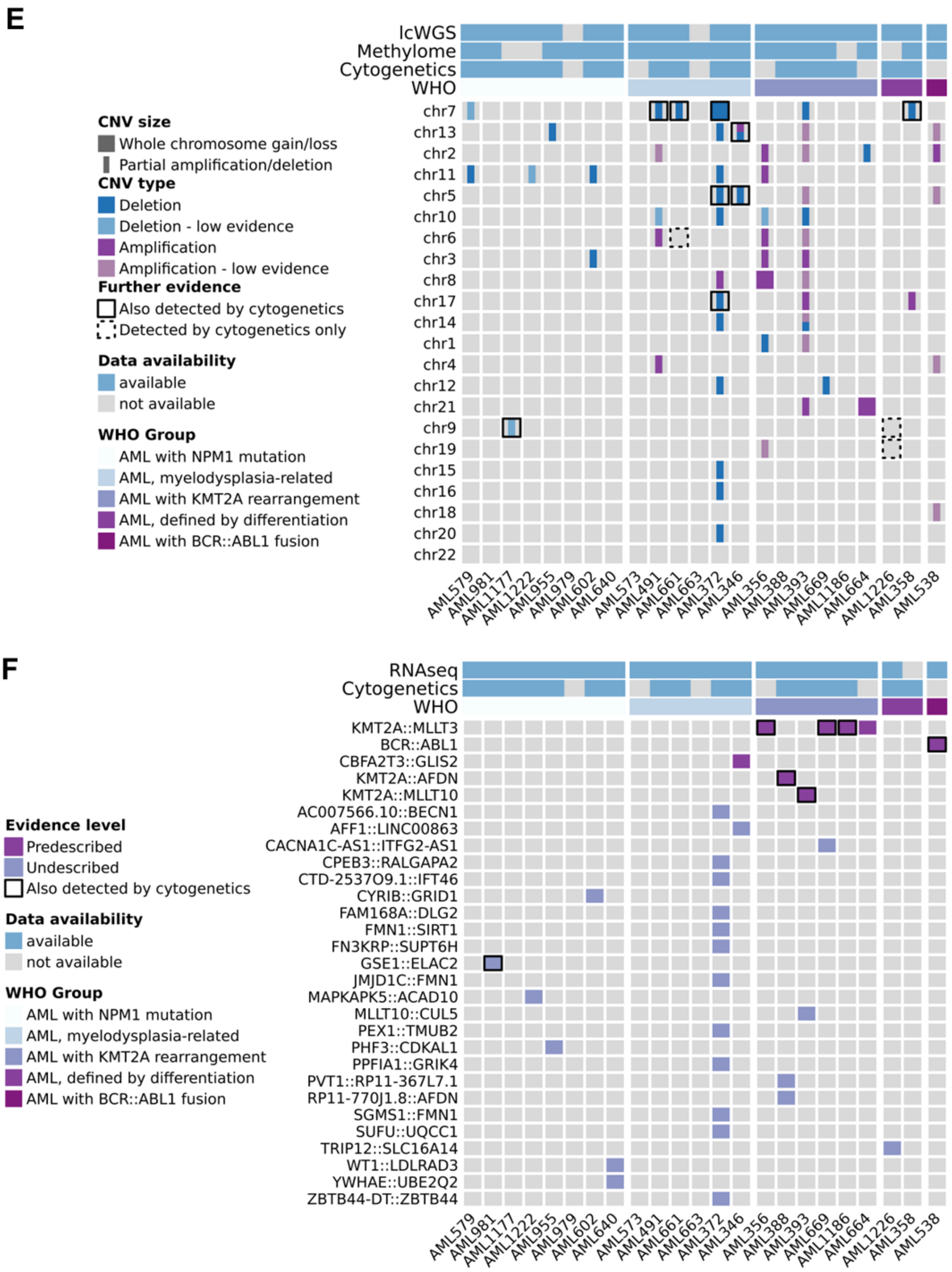

E
lcWGS
Methylome
Cytogenetics
WHO
CNV size
Whole chromosome gain/loss
Partial amplification/deletion
CNV type
Deletion
Deletion - low evidence
Amplification
Amplification - low evidence
Further evidence
Also detected by cytogenetics
Detected by cytogenetics only
Data availability
available
not available
WHO Group
AML with NPM1 mutation
AML, myelodysplasia-related
AML with KMT2A rearrangement
AML, defined by differentiation
AML with BCR::ABL1 fusion
chr7
chr13
chr2
chr11
chr5
chr10
chr6
chr3
chr8
chr17
chr14
chr1
chr4
chr12
chr21
chr9
chr19
chr15
chr16
chr18
chr20
chr22
AML579
AML981
AML1177
AML1222
AML955
AML979
AML602
AML640
AML573
AML491
AML661
AML663
AML372
AML346
AML356
AML388
AML393
AML669
AML1186
AML664
AML1226
AML358
AML538
F
RNAseq
Cytogenetics
WHO
Evidence level
Predescribed
Undescribed
Also detected by cytogenetics
Data availability
available
not available
WHO Group
AML with NPM1 mutation
AML, myelodysplasia-related
AML with KMT2A rearrangement
AML, defined by differentiation
AML with BCR::ABL1 fusion
KMT2A::MLLT3
BCR::ABL1
CBFA2T3::GLIS2
KMT2A::AFDN
KMT2A::MLLT10
AC007566.10::BECN1
AFF1::LINC00863
CACNA1C-AS1::ITFG2-AS1
CPEB3::RALGAPA2
CTD-2537O9.1::IFT46
CYRIB::GRID1
FAM168A::DLG2
FMN1::SIRT1
FN3KRP::SUPT6H
GSE1::ELAC2
JMJD1C::FMN1
MAPKAPK5::ACAD10
MLLT10::CUL5
PEX1::TMUB2
PHF3::CDKAL1
PPFIA1::GRIK4
PVT1::RP11-367L7.1
RP11-770J1.8::AFDN
SGMS1::FMN1
SUFU::UQCC1
TRIP12::SLC16A14
WT1::LDLRAD3
YWHAE::UBE2Q2
ZBTB44-DT::ZBTB44
AML579
AML981
AML1177
AML1222
AML955
AML979
AML602
AML640
AML573
AML491
AML661
AML663
AML372
AML346
AML356
AML388
AML393
AML669
AML1186
AML664
AML1226
AML358
AML538

G

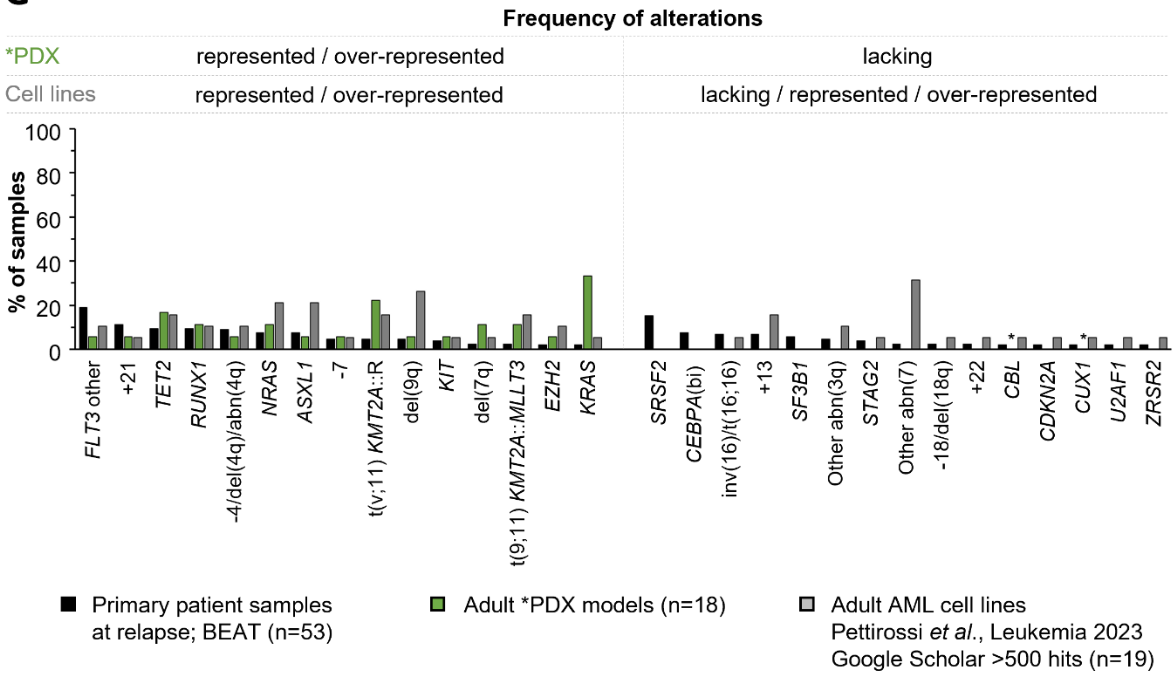
Frequency of alterations
*PDX
represented / over-represented
lacking
Cell lines
represented / over-represented
lacking / represented / over-represented
% of samples
FLT3 other
+21
TET2
RUNX1
-4/del(4q)/abn(4q)
NRAS
ASXL1
-7
t(v;11) KMT2A::R
del(9q)
KIT
del(7q)
t(9;11) KMT2A::MLLT3
EZH2
KRAS
SRSF2
CEBPA(bi)
inv(16)/t(16;16)
+13
SF3B1
Other abn(3q)
STAG2
Other abn(7)
-18/del(18q)
+22
CBL
CDKN2A
CUX1
U2AF1
ZRSR2
Primary patient samples at relapse; BEAT (n=53)
Adult *PDX models (n=18)
Adult AML cell lines Pettirossi et al., Leukemia 2023 Google Scholar >500 hits (n=19)

H

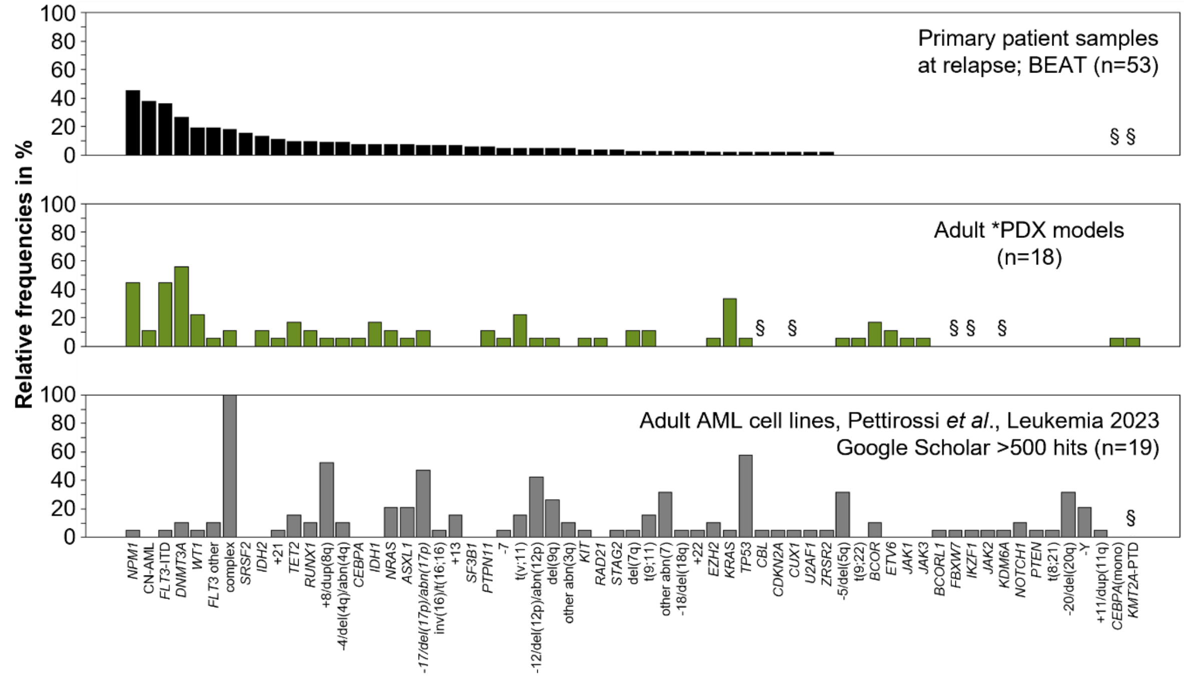
Relative frequencies in %
Primary patient samples at relapse; BEAT (n=53)
Adult *PDX models (n=18)
Adult AML cell lines, Pettirossi et al., Leukemia 2023 Google Scholar >500 hits (n=19)

I

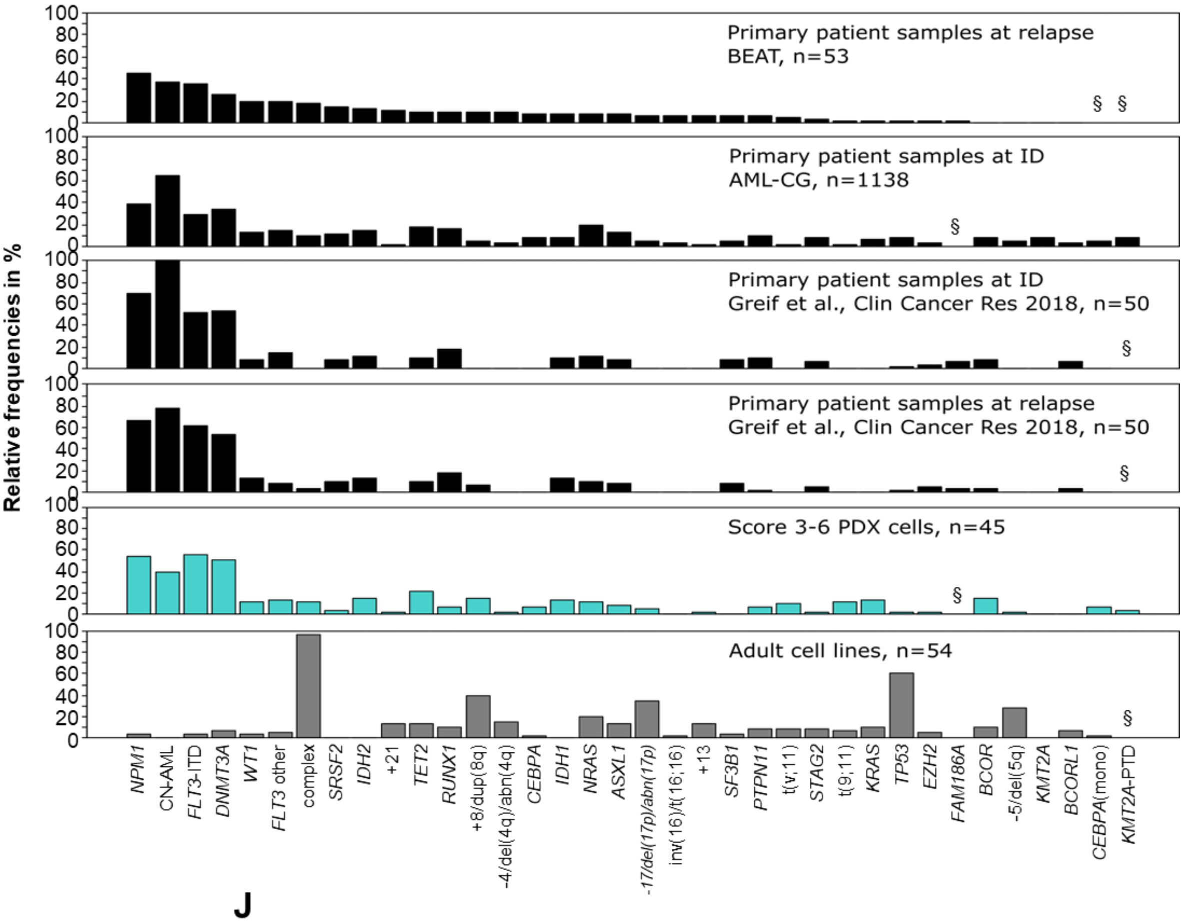
Relative frequencies in %
Primary patient samples at relapse
BEAT, n=53
Primary patient samples at ID
AML-CG, n=1138
Primary patient samples at ID
Greif et al., Clin Cancer Res 2018, n=50
Primary patient samples at relapse
Greif et al., Clin Cancer Res 2018, n=50
Score 3-6 PDX cells, n=45
Adult cell lines, n=54
NPM1
CN-AML
FLT3-ITD
DNMT3A
WT1
FLT3 other
complex
SRSF2
IDH2
+21
TET2
RUNX1
+8/dup(8q)
-4/del(4q)/abn(4q)
CEBPA
IDH1
NRAS
ASXL1
-17/del(17p)/abn(17p)
inv(16)/t(16;16)
+13
SF3B1
PTPN11
t(v;11)
STAG2
t(9;11)
KRAS
TP53
EZH2
FAM186A
BCOR
-5/del(5q)
KMT2A
BCORL1
CEBPA(mono)
KMT2A-PTD

J

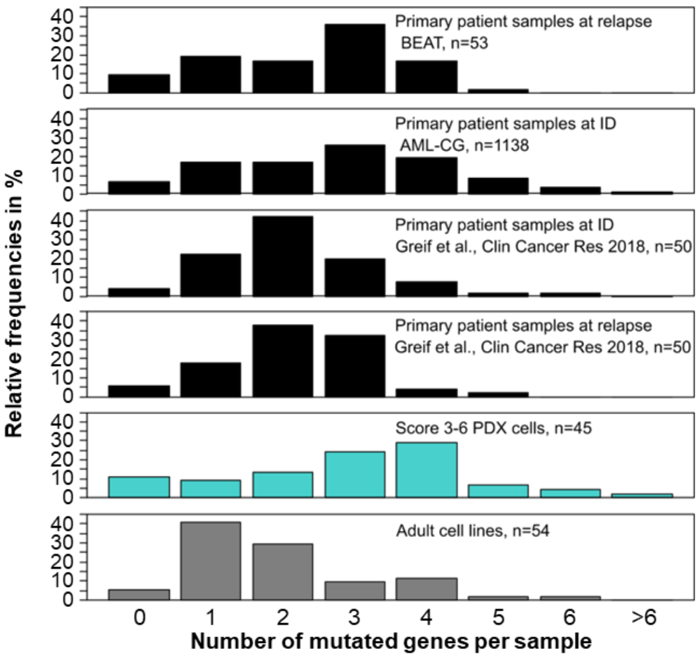
Relative frequencies in %
Primary patient samples at relapse
BEAT, n=53
Primary patient samples at ID
AML-CG, n=1138
Primary patient samples at ID
Greif et al., Clin Cancer Res 2018, n=50
Primary patient samples at relapse
Greif et al., Clin Cancer Res 2018, n=50
Score 3-6 PDX cells, n=45
Adult cell lines, n=54
Number of mutated genes per sample

**Supplemental Figure 3. Genomic characteristics of *PDX AML models better resemble primary AML cells than cell lines; related to Figure 2.**

**A-D *PDX AML models originate from patients with highly aggressive disease**

**A** Related to Figure 2A. Engrafted primary patient cells or PDX cells (n=62) were categorized into Score 2 to Score 6 as described in Figure 1B. Depicted are percentage of cases originating from ID (grey) or from R/R diseases (black).

**B** Related to Figure 2B. Kaplan-Meier curves showing the survival probability of all patients whose cells yielded *PDX models (Score 5/6, green) or did not meet the criteria of high score models (Score 2-4, blue). Log-rank test, $p$=0.00047.

**C** Related to Figure 2A. Left: percentage of models with wildtype *FLT3* (grey) or *FLT3-ITD* (black) are depicted ($p$=0.16). Right: percentage of models with wildtype *BCOR* (grey) or *BCOR* mutation (black) are depicted ($p$>0.999). The $p$-values were calculated with Fisher's exact test.

**D** Related to Figure 2A. Kaplan-Meier curves showing the survival probability of patients with (n=369) or without (n=2210) mutations in *KRAS* or *WT1* or *ETV6*. Data collected from AML-CG-1999 (n=864), AML-CG-2008 (n=274), AMLHD98A (n=557), AMLHD98B (n=160), and AML-SG07-04 (n=724).

**E,F** Related to Figure 2C. Copy Number Variants (CNVs) (**E**) and fusion transcripts (**F**) in *PDX AML samples are depicted.

**G-I** related to Figure 3B.

**G** Data were analyzed as described in Figure 2D, showing factors represented both in *PDX AML models and cell lines (left) or factors not available in *PDX AML models (right). As analyzed cell lines all show a highly complex karyotype, chromosomal alterations never occur as isolated events. §: factors not analyzed in a respective dataset.

**H** Data were analyzed as described in Figure 2D, but all analyzed alterations are depicted and sorted according to relative frequencies in relapsed patients of BEAT cohort.

**I** Relative frequencies of mutations and chromosomal alterations are depicted for primary AML samples at relapsed disease of BEAT cohort (n=53), primary AML samples at initial diagnosis of AML-CG study[23] (n=1138), primary AML samples at initial diagnosis or relapse of CN cohort (n=50), Score 3-6 PDX models generated from adult patients (n=45) and AML cell lines generated from adult patients (n=54). Events are included which are present in a minimum of 5% in one of the five cohorts. Mutations are sorted according to relative frequencies in relapsed patients of BEAT cohort.

**J** Number of detected mutations of samples/models of the cohorts listed in Figure S3I.

#### Supplemental Figure 4

A

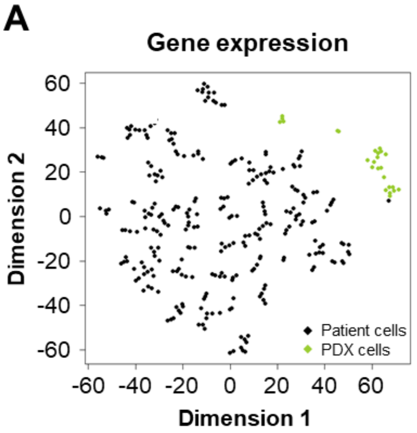


B

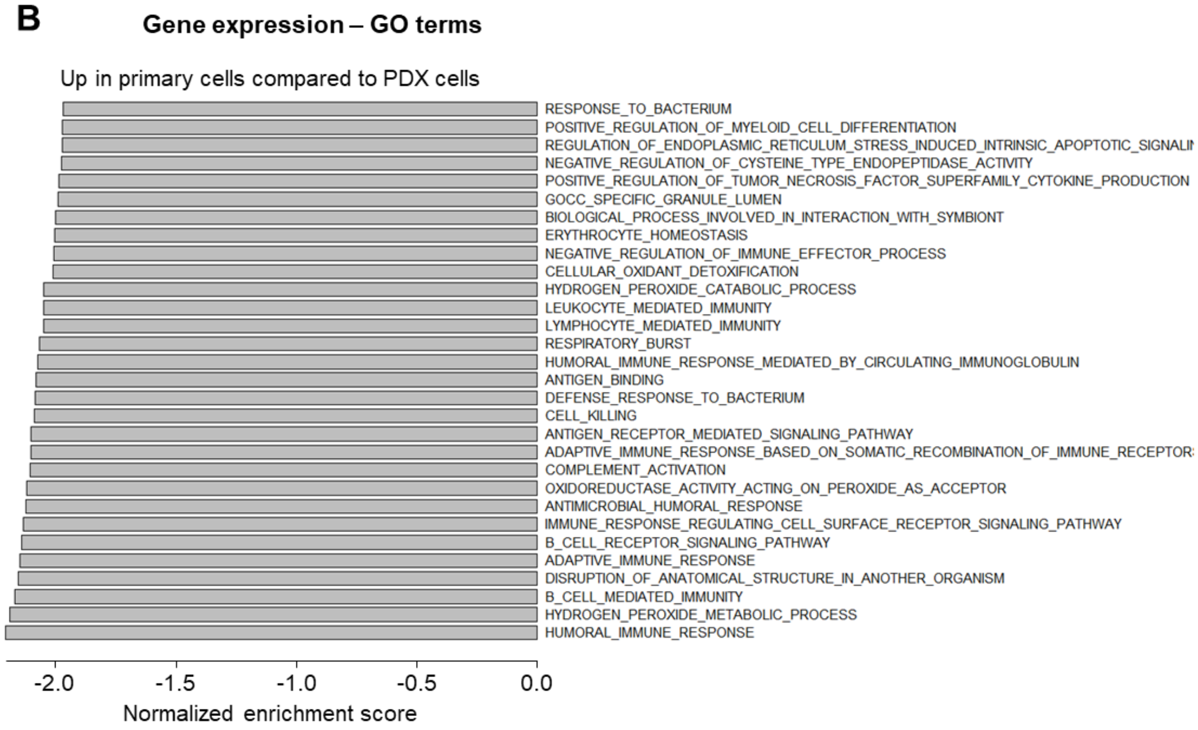

**C** **Gene expression – KEGG**

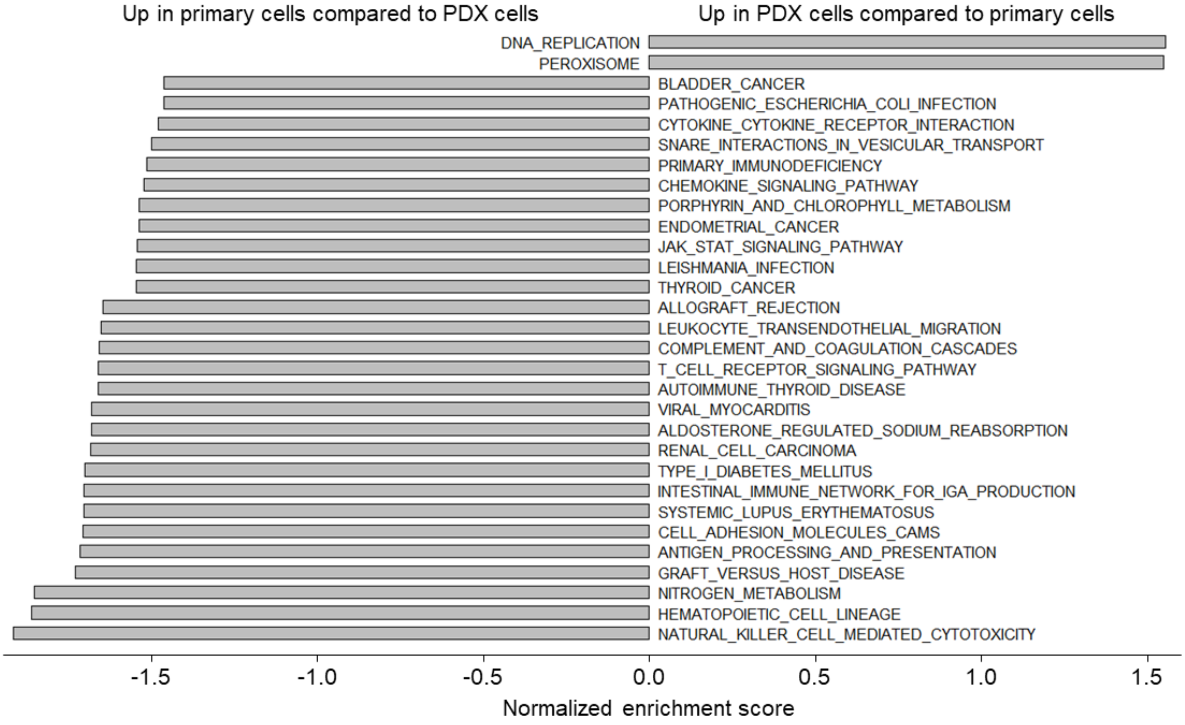


**D** **Gene expression – Hallmarks**

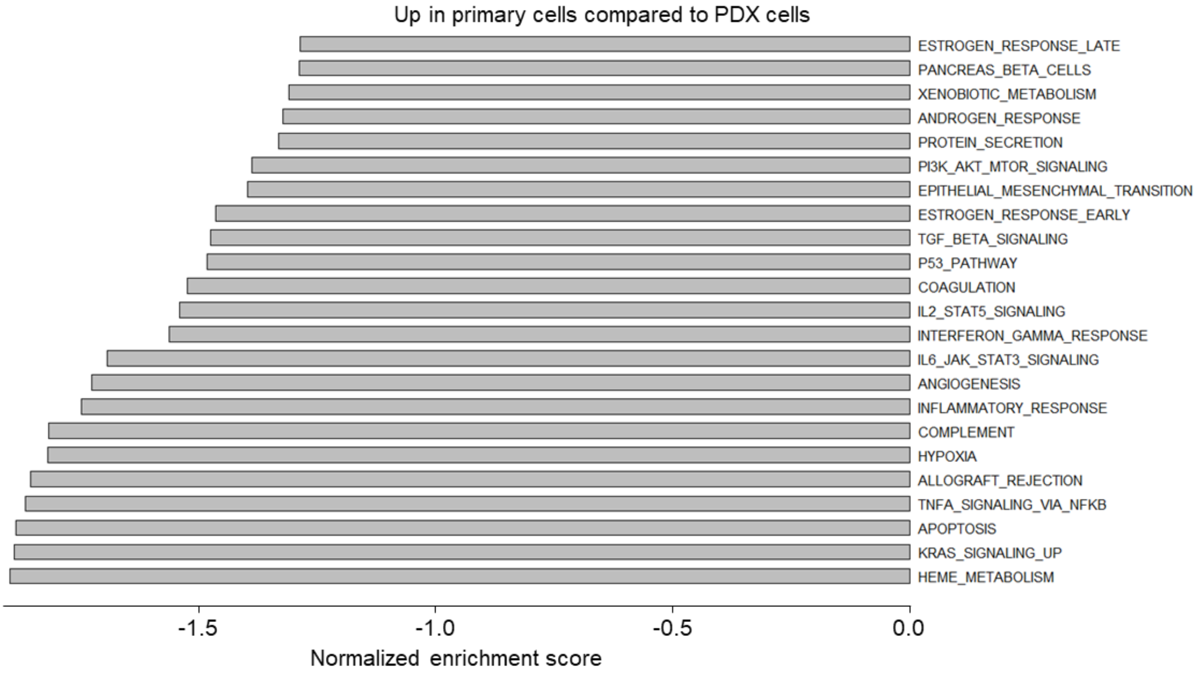

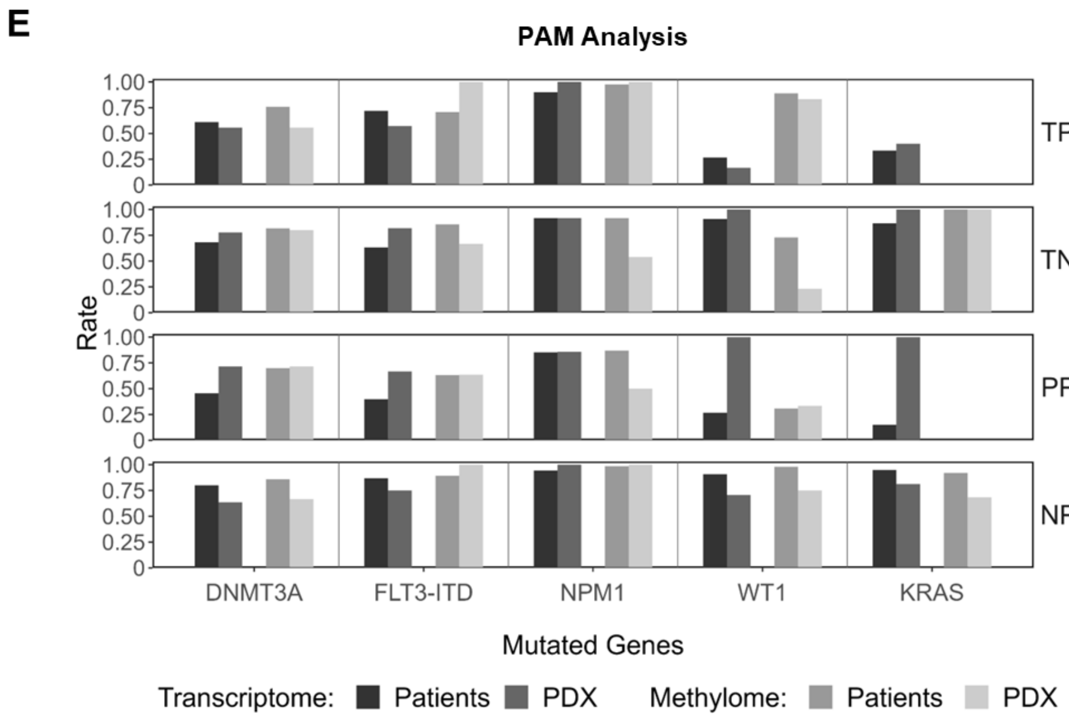
E
PAM Analysis
Rate
TPR
TNR
PPV
NPV
1.00
0.75
0.50
0.25
0
DNMT3A
FLT3-ITD
NPM1
WT1
KRAS
Mutated Genes
Transcriptome: Patients PDX Methylome: Patients PDX

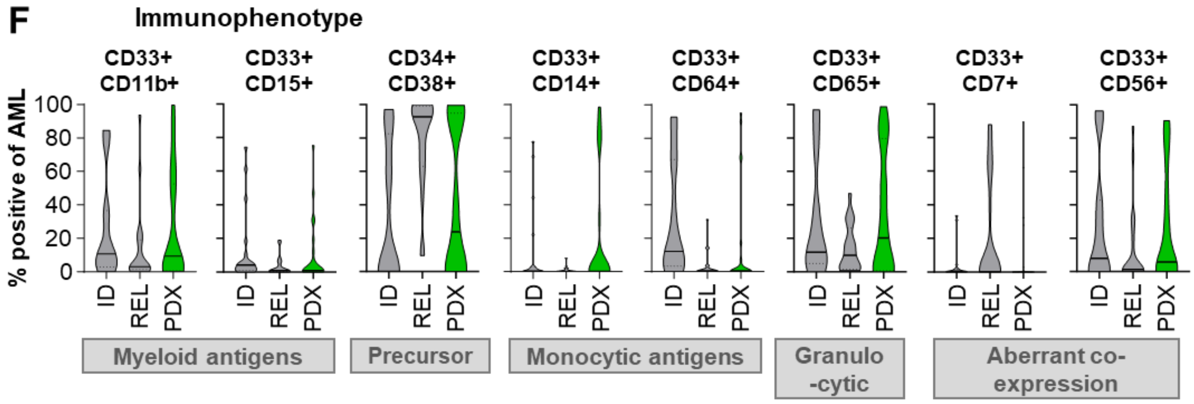
F
Immunophenotype
% positive of AML
CD33+ CD11b+
CD33+ CD15+
CD34+ CD38+
CD33+ CD14+
CD33+ CD64+
CD33+ CD65+
CD33+ CD7+
CD33+ CD56+
ID
REL
PDX
Myeloid antigens
Precursor
Monocytic antigens
Granulo -cytic
Aberrant co-expression

**Supplemental Figure 4. Methylome, transcriptome and surface protein characteristics show that *PDX AML models resemble primary AML cells, related to Figure 4**

**A** t-SNE plot of gene expression of *PDX AML models (n=23) and primary patient samples at ID (AML-CG Registry, n=271). Of note, both methylation and GE of AML-CG samples were conducted from bulk samples with variable percentage of blasts, while pure PDX AML cells were analyzed; furthermore, analyses for patient cells originating from relapsed disease were not available.

**B-D** GE data was compared between primary patient samples at ID and *PDX AML models. 30 most significantly altered pathways are depicted for GO terms (**B**) andfor KEGG (**C**), while all significantly altered pathways are depicted forHallmarks of Cancer (**D**).

**E** related to Figure 3B. PAM classifiers based on gene expression and DNA methylation profiling evaluated on *PDX AML models (n=19) and AML-CG Registry patient samples (n=224). Displayed are the true positive rate (TPR), true negative rate (TNR), positive predictive (PPV) and negative predictive value (NPV) for genes mutated in at least 20% of PDX samples.

**F** related to Figure 4C. Primary patient samples at ID (n=20), primary patient samples at relapse (n=20), and *PDX AML models were analyzed for cell surface marker expression (precursor, differentiation, aberrant co-expression) via flow cytometry. Primary patient samples are independent of PDX models. Percent antigen-positive cells within blast gate are depicted.

**Supplemental Figure 5**

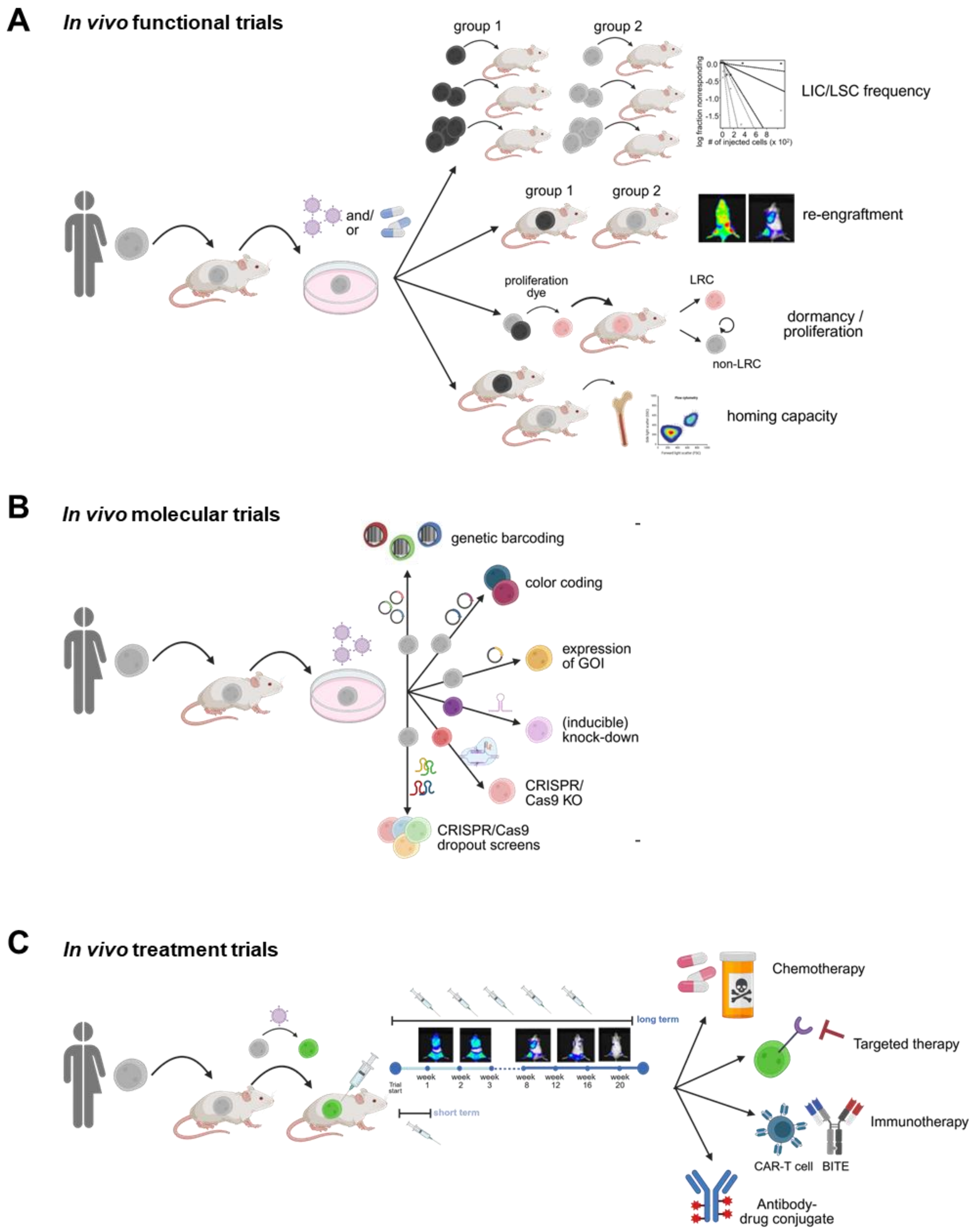

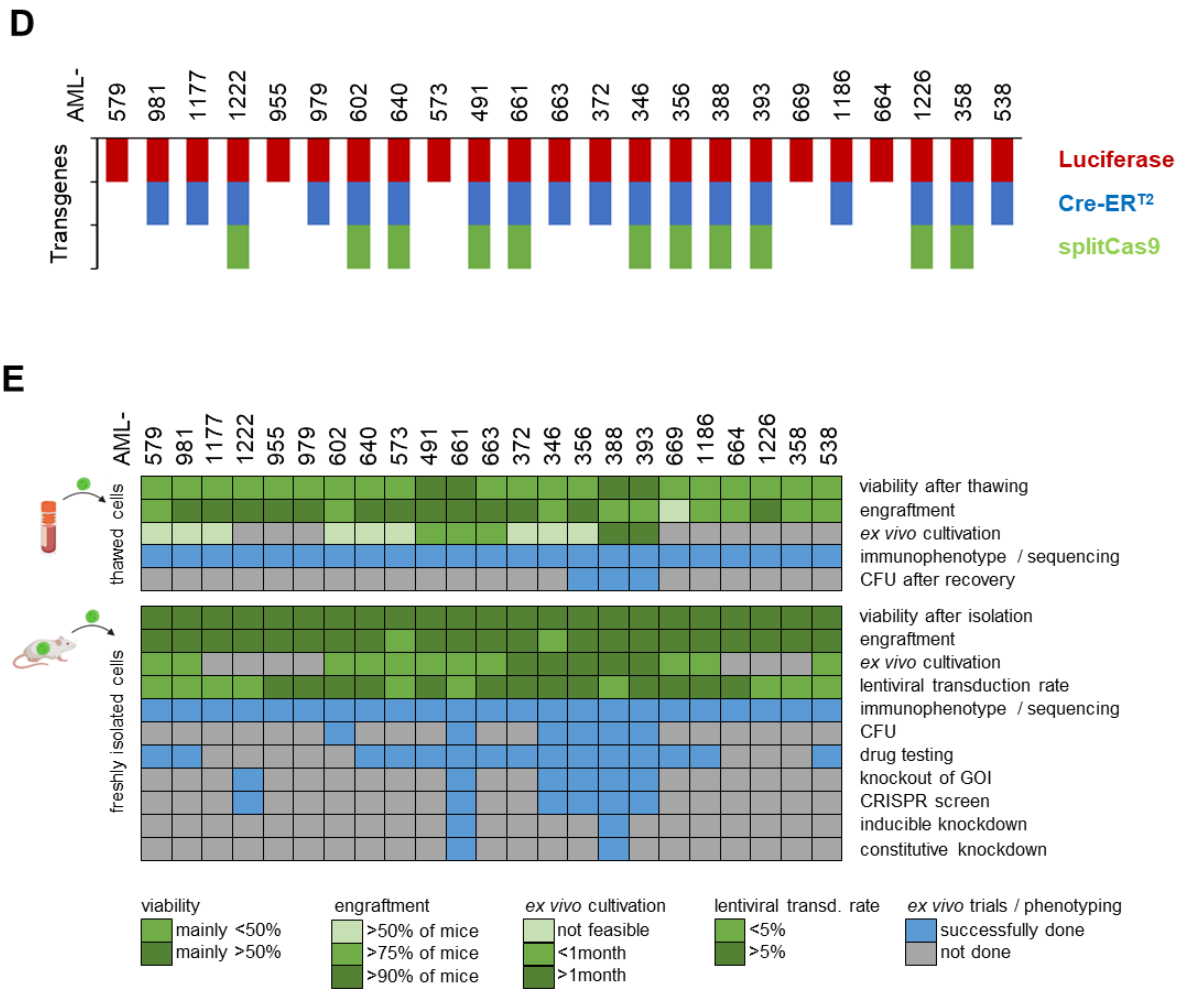

D
AML-
579 981 1177 1222 955 979 602 640 573 491 661 663 372 346 356 388 393 669 1186 664 1226 358 538
Transgenes
Luciferase
Cre-ER T2
splitCas9
E
AML-
579 981 1177 1222 955 979 602 640 573 491 661 663 372 346 356 388 393 669 1186 664 1226 358 538
thawed cells
viability after thawing
engraftment
ex vivo cultivation
immunophenotype / sequencing
CFU after recovery
freshly isolated cells
viability after isolation
engraftment
ex vivo cultivation
lentiviral transduction rate
immunophenotype / sequencing
CFU
drug testing
knockout of GOI
CRISPR screen
inducible knockdown
constitutive knockdown
viability
mainly <50%
mainly >50%
engraftment
>50% of mice
>75% of mice
>90% of mice
ex vivo cultivation
not feasible
<1month
>1month
lentiviral transd. rate
<5%
>5%
ex vivo trials / phenotyping
successfully done
not done

F

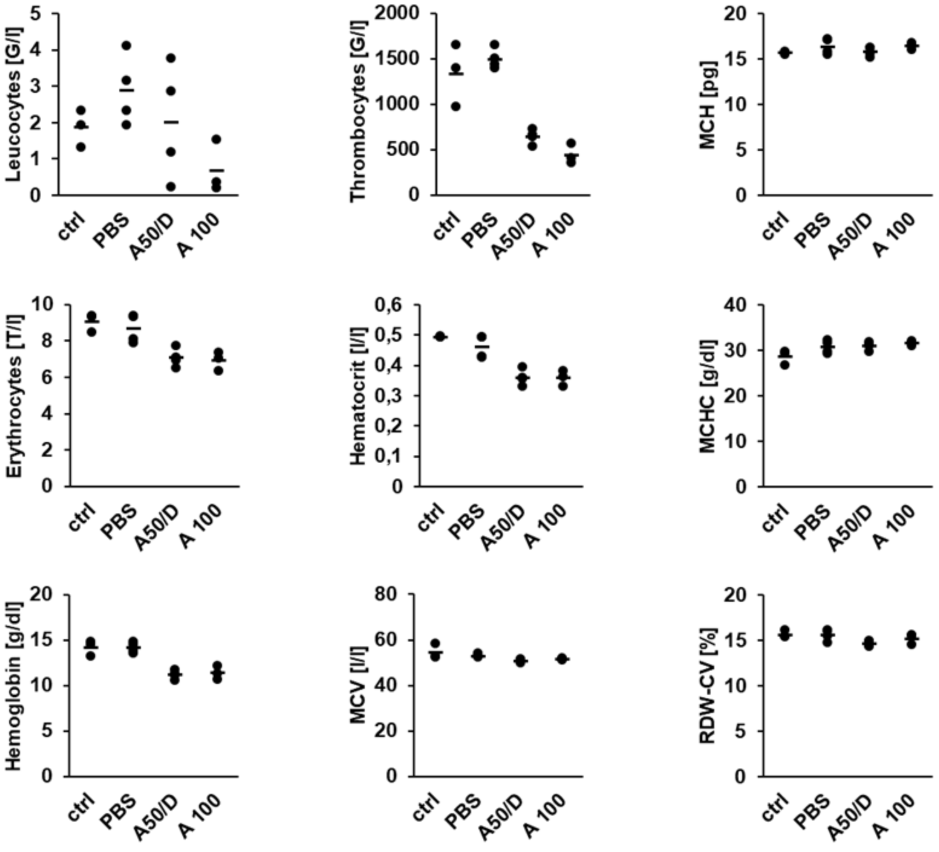
Leucocytes [G/l]
Thrombocytes [G/l]
MCH [pg]
Erythrocytes [T/l]
Hematocrit [l/l]
MCHC [g/dl]
Hemoglobin [g/dl]
MCV [l/l]
RDW-CV [%]
ctrl
PBS
A50/D
A 100

G

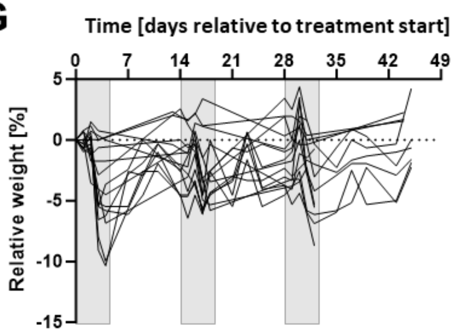
Time [days relative to treatment start]
0
7
14
21
28
35
42
49
Relative weight [%]
5
0
-5
-10
-15

H

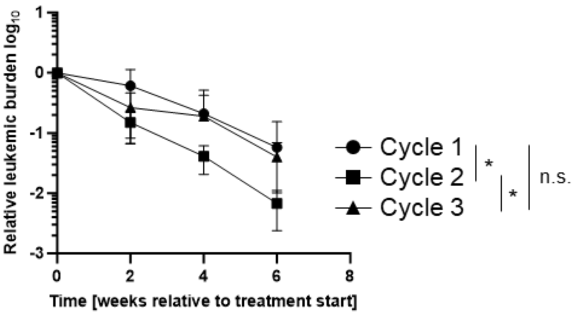
Relative leukemic burden log10
Time [weeks relative to treatment start]
Cycle 1
Cycle 2
Cycle 3
*
*
n.s.

I

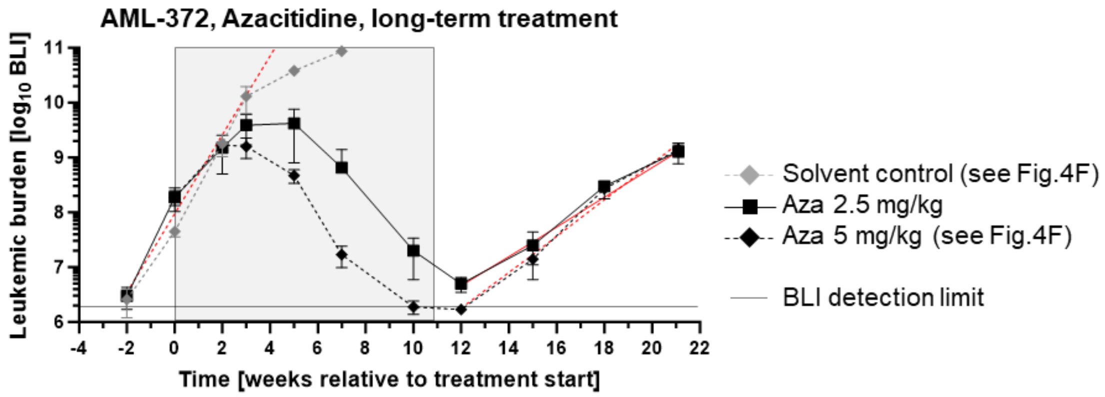


J

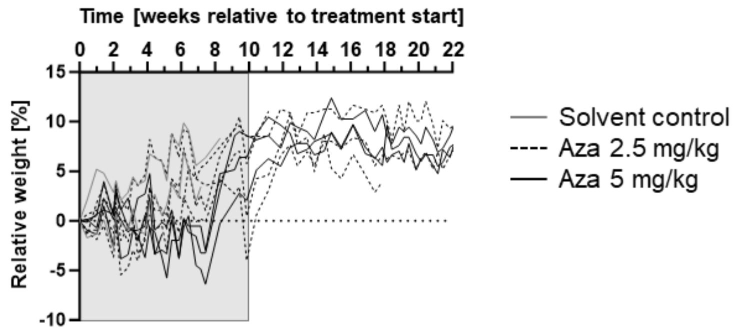


**Supplemental Figure 5. *PDX AML models allow a wide range of *ex vivo* and *in vivo* experiments, including long-term *in vivo* treatment trials, related to Figure 4.**

**A-E related to Figure 4A.**

**A** Scheme illustrating functional trials performed with *PDX AML models. Luciferase-transgenic models have been used to determine (i) leukemia-initiating cell (LIC) / leukemia stem cell (LSC) frequencies by limiting dilution transplantation assays, (ii) re-engraftment capacity of *ex vivo* or *in vivo*-treated or genetically modified cells, (iii) dormant versus proliferating cells, or (iv) homing capacity into BM.

**B** Scheme illustrating molecular trials performed with *PDX AML models. Lentiviral transduction facilitates (i) genetic barcoding to analyze functional heterogeneity and marking of single cell clones; (ii) expression of different fluorochromes for competitive *in vivo* trials; (iii) expression of mutant or wildtype gene of interest (GOI), (iv) (inducible) *knock-down* of GOI; (v) CRISPR *knock-out* of GOI, and (vi) *in vivo* CRISPR screens.

**C** Scheme illustrating preclinical therapy trials performed with *PDX AML models. Lentiviral transduction facilitates expression of enhanced firefly luciferase, enabling sensitive and repetitive disease monitoring and real-time follow-up of therapy response via bioluminescence imaging. Mice can be treated both short-term or long-term with either (i) chemotherapy, (ii) targeted therapy, (iii) immunotherapy such as CAR-T cells, bi-specific T cell engager antibodies or immune activation, or (iv) antibody-drug-conjugates.

**D** AML PDX cells were genetically modified and enriched to express enhanced firefly luciferase for bioluminescence *in vivo* imaging (red), Cre-ER$^{T2}$ for Tamoxifen-inducible expression of shRNAs and *knock-down* of target genes (blue), or split-Cas9 for *knock-out* of target genes (green).

**E** Thawed and freshly isolated cells of *PDX AML models have been used for diverse *ex vivo* trials, including phenotypic characterization, colony forming unit assays, *ex vivo* drug testing, *ex vivo* growth characteristics, *knock-down* or *knock-out* approaches (Table S3 and unpublished). Due to reduced viability after thawing, most assays can only be performed with freshly isolated cells.

**F-H Related to Figures 4B-D.**

**F Complete blood count analysis** of mice without PDX cell transplantation (ctrl, n=3), or of mice transplanted with AML-491 PDX cells treated with PBS (n=4), Cytarabine 50 mg/kg and DaunoXome 1 mg/kg (n=4), or Cytarabine 100 mg/kg (n=3) four days after last treatment.

**G Relative body weight** of mice treated with 1$^{st}$, 2$^{nd}$ or 3$^{rd}$ cycle of chemotherapy.

**H Relative drop in leukemic burden** under chemotherapy (1$^{st}$, 2$^{nd}$ or 3$^{rd}$ cycle). Relative tumor burden after start of chemotherapy is depicted for 1$^{st}$ (circle, n=24), 2$^{nd}$ (square, n=7) and 3$^{rd}$ (triangle, n=13) cycle of chemotherapy. Mean +/- standard deviation is shown. * $p$<0.01 as tested by two-sided unpaired t-test with Welch´s correction. n.s. not significant.

**I-J Related to Figure 4F, AML-372.**

**I** Experiment was performed as described in Figure 5E; however, mice were treated with 2.5 mg/kg Azacitidine.

**J Relative weight** of mice treated with Azacitidine.

#### References


1. Farge T, Saland E, de Toni F, et al. Chemotherapy-Resistant Human Acute Myeloid Leukemia Cells Are Not Enriched for Leukemic Stem Cells but Require Oxidative Metabolism. *Cancer Discov*. 2017;7(7):716-735.
2. Woiterski J, Ebinger M, Witte KE, et al. Engraftment of low numbers of pediatric acute lymphoid and myeloid leukemias into NOD/SCID/IL2Rcgammanull mice reflects individual leukemogenecity and highly correlates with clinical outcome. *Int J Cancer*. 2013;133(7):1547-1556.
3. Diaz de la Guardia R, Velasco-Hernandez T, Gutierrez-Aguera F, et al. Engraftment characterization of risk-stratified AML in NSGS mice. *Blood Adv*. 2021;5(23):4842-4854.
4. Townsend EC, Murakami MA, Christodoulou A, et al. The Public Repository of Xenografts Enables Discovery and Randomized Phase II-like Trials in Mice. *Cancer Cell*. 2016;29(4):574-586.
5. Percie du Sert N, Hurst V, Ahluwalia A, et al. The ARRIVE guidelines 2.0: Updated guidelines for reporting animal research. *PLoS Biol*. 2020;18(7):e3000410.
6. Bonnet D. In vivo evaluation of leukemic stem cells through the xenotransplantation model. *Curr Protoc Stem Cell Biol*. 2008;Chapter 3:Unit 3 2.
7. Hutter G, Nickenig C, Garritsen H, et al. Use of polymorphisms in the noncoding region of the human mitochondrial genome to identify potential contamination of human leukemia-lymphoma cell lines. *Hematol J*. 2004;5(1):61-68.
8. Terziyska N, Castro Alves C, Groiss V, et al. In vivo imaging enables high resolution preclinical trials on patients' leukemia cells growing in mice. *PLoS One*. 2012;7(12):e52798.
9. Bahrami E, Schmid JP, Jurinovic V, et al. Combined proteomics and CRISPR–Cas9 screens in PDX identify ADAM10 as essential for leukemia in vivo. *Mol Cancer*. 2023;22(1):107.
10. Carlet M, Volse K, Vergalli J, et al. In vivo inducible reverse genetics in patients' tumors to identify individual therapeutic targets. *Nat Commun*. 2021;12(1):5655.
11. Vick B, Rothenberg M, Sandhofer N, et al. An advanced preclinical mouse model for acute myeloid leukemia using patients' cells of various genetic subgroups and in vivo bioluminescence imaging. *PLoS One*. 2015;10(3):e0120925.
12. Wermke M, Camgoz A, Paszkowski-Rogacz M, et al. RNAi profiling of primary human AML cells identifies ROCK1 as a therapeutic target and nominates fasudil as an antileukemic drug. *Blood*. 2015;125(24):3760-3768.
13. Metzeler KH, Herold T, Rothenberg-Thurley M, et al. Spectrum and prognostic relevance of driver gene mutations in acute myeloid leukemia. *Blood*. 2016;128(5):686-698.
14. Daenekas B, Perez E, Boniolo F, et al. Conumee 2.0: enhanced copy-number variation analysis from DNA methylation arrays for humans and mice. *Bioinformatics*. 2024;40(2).
15. Scheinin I, Sie D, Bengtsson H, et al. DNA copy number analysis of fresh and formalin-fixed specimens by shallow whole-genome sequencing with identification and exclusion of problematic regions in the genome assembly. *Genome Res*. 2014;24(12):2022-2032.
16. Adalsteinsson VA, Ha G, Freeman SS, et al. Scalable whole-exome sequencing of cell-free DNA reveals high concordance with metastatic tumors. *Nat Commun*. 2017;8(1):1324.
17. Janjic A, Wange LE, Bagnoli JW, et al. Prime-seq, efficient and powerful bulk RNA sequencing. *Genome Biol*. 2022;23(1):88.
18. Buchner T, Berdel WE, Haferlach C, et al. Age-related risk profile and chemotherapy dose response in acute myeloid leukemia: a study by the German Acute Myeloid Leukemia Cooperative Group. *J Clin Oncol*. 2009;27(1):61-69.
19. Braess J, Amler S, Kreuzer KA, et al. Sequential high-dose cytarabine and mitoxantrone (S-HAM) versus standard double induction in acute myeloid leukemia-a phase 3 study. *Leukemia*. 2018;32(12):2558-2571.
20. Schlenk RF, Dohner K, Mack S, et al. Prospective evaluation of allogeneic hematopoietic stem-cell transplantation from matched related and matched unrelated donors in younger adults with high-risk acute myeloid leukemia: German-Austrian trial AMLHD98A. *J Clin Oncol*. 2010;28(30):4642-4648.

21. Schlenk RF, Frohling S, Hartmann F, et al. Intensive consolidation versus oral maintenance therapy in patients 61 years or older with acute myeloid leukemia in first remission: results of second randomization of the AML HD98-B treatment Trial. *Leukemia*. 2006;20(4):748-750.
22. Schlenk RF, Lubbert M, Benner A, et al. All-trans retinoic acid as adjunct to intensive treatment in younger adult patients with acute myeloid leukemia: results of the randomized AMLSG 07-04 study. *Ann Hematol*. 2016;95(12):1931-1942.
23. Rausch C, Rothenberg-Thurley M, Dufour A, et al. Validation and refinement of the 2022 European LeukemiaNet genetic risk stratification of acute myeloid leukemia. *Leukemia*. 2023;37(6):1234-1244.
24. Bottomly D, Long N, Schultz AR, et al. Integrative analysis of drug response and clinical outcome in acute myeloid leukemia. *Cancer Cell*. 2022;40(8):850-864 e859.
25. Greif PA, Hartmann L, Vosberg S, et al. Evolution of Cytogenetically Normal Acute Myeloid Leukemia During Therapy and Relapse: An Exome Sequencing Study of 50 Patients. *Clin Cancer Res*. 2018;24(7):1716-1726.
26. Pettirossi V, Venanzi A, Spanhol-Rosseto A, et al. The gene mutation landscape of acute myeloid leukemia cell lines and its exemplar use to study the BCOR tumor suppressor. *Leukemia*. 2023;37(2):473-477.
27. Tibshirani R, Hastie T, Narasimhan B, Chu G. Diagnosis of multiple cancer types by shrunken centroids of gene expression. *Proc Natl Acad Sci U S A*. 2002;99(10):6567-6572.
28. Vosberg S, Kerbs P, Jurinovic V, et al. DNA Methylation Profiling of AML Reveals Epigenetic Subgroups with Distinct Clinical Outcome. *Blood*. 2019;134.
29. Perova Z, Martinez M, Mandloi T, et al. PDCM Finder: an open global research platform for patient-derived cancer models. *Nucleic Acids Res*. 2023;51(D1):D1360-D1366.
